# Targeting Galectin-3 C-epitope oligomers associated maladaptive mechanotransductive signaling in pressure-overload induced left ventricular cardiac hypertrophy

**DOI:** 10.64898/2026.02.09.704762

**Authors:** Puja Laxmanrao Shinde, Vikas Kumar, Siddhartha Singh, K.C. Sivakumar, P Abhirami, Amit Mishra, Rashmi Mishra

**Affiliations:** Translational Mechanobiology Laboratory, BRIC-Rajiv Gandhi Centre for Biotechnology (RGCB), Thycaud PO, Poojappura, Thiruvananthapuram-695014, Kerala, India; Cardiovascular Diseases and Diabetes Biology Laboratory, BRIC-Rajiv Gandhi Centre for Biotechnology (RGCB), Thiruvananthapuram-695014, Kerala, India; Manipal Academy of Higher Education, Manipal-576104, Karnataka, India; Regional Centre for Biotechnology, Faridabad-121001, Haryana, India; Distributed Information Sub-Centre, BRIC-Rajiv Gandhi Centre for Biotechnology (RGCB), Thycaud PO, Poojappura, Thiruvananthapuram-695014, Kerala, India; Department of Medicine, New York University Grossman School of Medicine, New York, NY-10016, USA; Cellular and Molecular Neurobiology Unit, Indian Institute of Technology Jodhpur-342037, Rajasthan, India

**Keywords:** Galectin-3 C-epitope oligomers, Cardiomyocyte mechanotransduction, Signaling in cardiac hypertrophy, gallic acid

## Abstract

**Background:** Aging and various pathological conditions lead to pressure-overload in the left ventricle, promoting maladaptive hypertrophic remodeling and subsequent cardiac dysfunction, ultimately increasing the risk of heart failure. Galectin-3 (Gal-3) plays a central role in this process; however, its critical intracellular functions complicate direct therapeutic targeting. Notably, pathological microenvironments trigger the proteolytic cleavage of Gal-3 into distinct N- and C-terminal fragments. The specific contributions of these cleaved epitope forms to adverse cardiomyocyte mechanotransduction, and their potential as precision therapeutic targets in contrast to the full-length protein, remain unresolved.

**Methods:** To address this gap, we combined rodent models of aging and pressure-overload (PO) –induced cardiac hypertrophy with PO mechanobiology-driven in vitro assays and validation in human cardiac tissue and serum. Gal-3 epitope abundance, localization, phosphorylation, oligomerization, and downstream signaling were quantified using biochemical, imaging, and functional approaches.

**Results:** We found that extracellular oligomers of the Gal-3 C-terminal epitope accumulated in serum and on cardiomyocyte surfaces in hypertrophic rodents and human subjects, where they correlated with adverse remodeling and cardiomyocyte loss. Treatment with Amalaki Rasayana (AR), a standardized nutraceutical-based cardioprotective Ayurvedic phytomedicine, and its bioactive component gallic acid (GA) significantly reduced circulating and surface-associated Gal-3 C-epitope oligomers and attenuated hypertrophy-associated cytotoxic signaling. Mechanistically, AR/GA enhanced Ser6 phosphorylation of Gal-3, promoting intracellular retention, while limiting pathological secretion and deleterious extracellular oligomerization. Following AR/GA treatment, the binding of preformed Gal-3 C-epitope oligomers to cardiomyocyte surfaces were further inhibited, thereby suppressing maladaptive mechanotransductive signaling. Importantly, circulating Gal-3 C-epitope oligomers, together with atrial natriuretic peptide (ANP), constituted a drug-responsive biomarker panel that accurately tracked hypertrophy regression, serving as an indicator of drug efficacy.

**Conclusions:** In summary, Gal-3 C-epitope oligomers represent pathogenic signaling, drug-responsive therapeutic targets and circulating biomarkers of cardiac hypertrophy, with broader relevance to other Gal-3–driven neoplastic, fibrotic, and inflammatory diseases.

**Highlights:**

- Galectin-3 C-epitope forms pathogenic extracellular oligomers in pressure-overload induced cardiac hypertrophy.
- Surface binding of excess C-epitope oligomers triggers an adverse mechanotransductive remodelling in cardiomyocytes.
- Phytomedicine **-**Amalaki Rasayana (AR) and its key bioactive compound, gallic acid (GA), effectively inhibit the surface binding and mitigate harmful effects of Gal-3 C-epitope oligomers.
- AR/GA induces the phosphorylation of Gal-3, limiting pathological secretion, extracellular oligomer assembly, and glycan-mediated surface binding.
- Intracellular retention of Gal-3 preserves its crucial cellular functions.
- Circulating levels of Gal-3 C-epitope oligomers can be used to monitor the therapeutic regression of cardiac hypertrophy, serving as an indicator of drug efficacy

**GRAPHICAL ABSTRACT:** 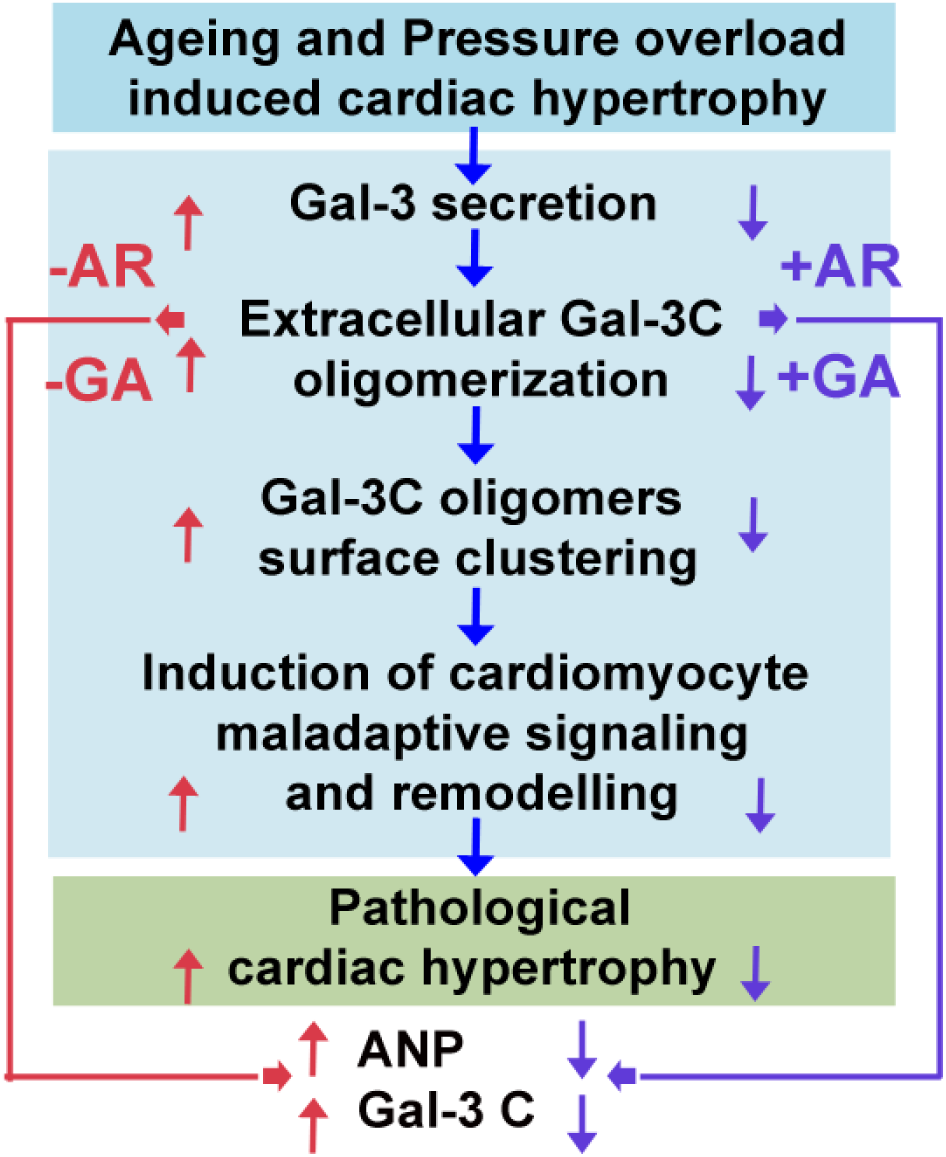

Pressure-overload induces excessive secretion of Galectin-3 and triggers its proteolytic cleavage into the N- and C-terminal epitopes. The extracellular oligomerization of the C-terminal epitope facilitates high-affinity glycan binding and promotes pathological mechanotransductive signaling in cardiomyocytes. Phytochemical modulation by Amalaki rasayana and its bioactive component, gallic acid, induces the phosphorylation of Gal-3, limiting pathological secretion, extracellular oligomer assembly, and glycan-mediated surface binding, while preserving intracellular function. Circulating levels of Gal-3 C-epitope oligomers can be used to monitor the therapeutic regression of cardiac hypertrophy, serving as an indicator of drug efficacy.

## 1. INTRODUCTION

Heart failure is a leading cause of morbidity and mortality worldwide and arises when the left ventricle (LV) fails to pump blood effectively, resulting in symptoms such as breathlessness, fatigue, and progressive multi-organ dysfunction [1]. Chronic pressure-overload, most commonly caused by hypertension and aortic stenosis, induces LV hypertrophy—characterized by increases in cardiomyocyte length, thickness, and myocardial mass. Although initially compensatory, sustained hypertrophy ultimately becomes maladaptive, leading to cardiomyocyte loss, ventricular stiffening, and progression to heart failure. A key pathological feature of this transition is myocardial fibrosis, which further compromises LV function by increasing tissue stiffness and impairing contractile efficiency [2, 3, 4, 5].

Galectin-3 (Gal-3), a β-galactoside-binding lectin, is a well-recognized mediator of cardiac remodelling, inflammation, and fibrosis, particularly under conditions of pressure-overload. Gal-3 is expressed by multiple cardiac cell types, and elevated circulating Gal-3 levels are associated with adverse cardiovascular outcomes and systemic inflammation. Importantly, proteolytic processing of Gal-3 generates a C-terminal epitope (≈10.5 kDa) containing the carbohydrate recognition domain (CRD), which can self-associate into pentameric oligomers (≈52 kDa) and higher-order multimers. Emerging evidence suggests that these Gal-3 C-epitope oligomers may preferentially engage cell-surface glycan receptors and activate hypertrophic signalling pathways, thereby exerting deleterious effects on cardiomyocytes [6, 7,8,9,10,11,12,13,14,15,16,17,18]. However, whether N- or C-epitope-specific Gal-3 pools differentially regulate cardiomyocyte fate during pressure-overload induced hypertrophy remains unclear.

In the present study, we investigated whether selective targeting of Gal-3 C-epitope oligomers, rather than global inhibition of total or full-length Gal-3, represents a more precise therapeutic strategy—one that mitigates hypertrophy-associated pathology while preserving Gal-3’s essential physiological roles in immunity and intracellular signalling [14,17]. We further examined the Gal-3 C-epitope-modulating potential of Amalaki rasayana (AR), a thoroughly standardized cardioprotective Ayurvedic formulation derived from *Phyllanthus emblica* (Indian gooseberry) and enriched in gallic acid (GA), which has demonstrated efficacy in preclinical models of cardiac hypertrophy and ageing (please see Supplementary file 1, Section B, for more details on our previously established and published protocols on AR reproducible preparation and standardization) [19, 20, 21, 22].

Using complementary *in vitro* approaches and *in vivo* models of biological ageing and pressure-overload induced LV hypertrophy, we assessed whether AR treatment reduces Gal-3 C-epitope oligomer formation and alleviates maladaptive cardiac remodelling induced cell-death. Collectively, our findings identify Gal-3 C-epitope oligomers as pathogenic mediators and drug-responsive biomarkers of cardiac hypertrophy, and support AR as a rapidly translatable phytotherapeutic strategy for precision management of pressure-overload associated heart failure.

## 2. MATERIALS AND METHODS

For details on our previously published methodologies including standardized preparation of Amalaki rasayana, its chemical fingerprinting, component quantitation, quality control, reproducibility marker identification and quality assurance regulatives; generation of the rodent model of cardiac hypertrophy, AR dose standardization [20,21] and the sources of reagents and kits used in this study, please refer to the supplementary file 1, section B.

### 2.1. Pressure-Overload Cellular Model

An in vitro pressure-overload (PO) model using weight-based method is well established [23, 24]. Cardiomyoblasts were cultured in low-glucose DMEM supplemented with 10% FBS and antibiotics in 6-well plates. After 24 hours, differentiation was induced by switching to medium containing 1% FBS for 24 hours [25, 26]. At approximately 80% confluence, the medium was replaced with CO₂-independent medium containing 5% FBS to minimize pH fluctuations under reduced gas exchange conditions. During model standardization, mechanical stress was applied by placing sterile weights (ranging from 2 to 20 g) on the medium surface to mimic pathological cardiomyocyte overload. Cells were exposed to pressure for 12–48 hours, with controls (No-PO) cultured without weights but under similar gas exchange conditions using dummy lids. A 10 g weight (∼1 g/cm²) was found to significantly enhance the cell surface area of cardiomyocytes by 3-4 fold, similar to the hypertrophy observed in *in vivo* animal models. Consequently, this weight was selected for subsequent PO-induced cardiomyocyte hypertrophy experiments. For these experiments, three independent replicates per group (No-PO and PO) were treated with either AR (100 µg/mL) or GA (100 µM), based on doses determined via viability assays. Three additional experimental sets remained untreated. Cells were then harvested for analysis of hypertrophic markers, protein expression, signaling pathways, and mechanical load-induced hypertrophy validation.

### 2.2. Dot Blot ELISA

Proteins were extracted using a lysis buffer, and concentrations were determined by BCA assay for normalization. Equal amounts (6 µg in 2.2 µ L) were spotted onto 0.45 µm nitrocellulose membranes (Amersham), air-dried for 2 hours at room temperature, and blocked overnight at 4°C with 3% BSA in TBST (0.1% Tween-20). After three washes with TBST, membranes were incubated with primary antibodies, diluted in blocking buffer, for 1 hour, followed by HRP-conjugated secondary antibodies for 1 hour at room temperature. Signals were detected using enhanced chemiluminescence and captured digitally. Signals were recorded at various time intervals until saturation was reached. The images just below signal saturation were used for densitometric quantification. Dot intensities were analyzed with Fiji software, and inverted images of the blots were included in the figures for better visualization of weaker signals. The dot blots were stained with Commassie Blue dye for the assessment of equal sample loading. Original blots and loading control images are available in the supplementary files.

### 2.3. Galectin-3 WT and Mutant Conditioned Medium Experiments

HEK293 cells were transfected with CRD-WT-Gal-3-EGFP (wild-type Gal-3) or CRD-MUT-Gal-3-EGFP (R186S mutant Gal-3) plasmids [27]. Following 24-hour pressure-overload treatment, conditioned media (CM) containing secreted WT or mutant CRD galectin-3 proteins were collected. Cardiomyocytes were incubated with the conditioned media for 10 hours. After incubation, cells were fixed with 1.5% PFA for surface imaging or stained with rhodamine phalloidin to visualize F-actin stress fibers and quantify changes in cardiomyocyte surface area.

### 2.4. Cell Surface-Bound Galectin-3 Extraction via Acid Wash

At the end of PO and AR/GA treatments, media were collected and cells were placed on ice. Cells were incubated with complete medium adjusted to pH 2.0 for several minutes to strip surface-bound gal-3. The released gal-3 was analyzed in the acid-wash medium, normalized to the normal pH, while cells were lysed separately to assess intracellular gal-3 levels.

### 2.5. Human Serum Conditioned Media Experiment

Human serum was diluted in cardiomyocyte culture medium at 1:20, 1:40, and 1:60 ratios. Differentiated cardiomyocytes were incubated with these dilutions for 10 hours before fixation. Samples were processed to assess cell surface area and to detect surface-bound gal-3 N- and C-epitope using antibody staining.

### 2.6. Immunoprecipitation, Gal-3 N- and C-Epitope Enrichment from Human Serum, and Lactose Competition Assay

Selective enrichment of gal-3 N- or C-epitope from aged human serum samples (n=3) was performed using Dynabeads Protein A immunoprecipitation according to the manufacturer’s protocol. Dynabeads (1.5 mg) were conjugated with 4 µg of anti-N- or anti-C-epitope gal-3 antibodies. A total of 1600 µg serum protein (20 µ L serum diluted in 580 µ L PBST) was incubated overnight at 4°C with antibody-conjugated beads on a rotator. The following day, supernatants were collected, and 225 µ L aliquots were incubated overnight at 4°C with or without 34 mM lactose or 100 µM GA. Lactose/GA-treated and untreated supernatants (225 µ L each) were mixed 1:1 with cardiomyocyte culture medium and incubated for 10 hours. After incubation, cells were fixed and surface area analyzed by capturing five random images per condition, quantifying approximately 200 cells. Equal volumes of supernatants were analyzed by Western blot to confirm N- and C-epitope enrichment, with reciprocal enrichment expected (e.g., N-epitope pull-down enriching C-epitope in the supernatant).

### 2.7. Surface Visualization of N- and C-Epitope of Galectin-3

To visualize only surface-bound N- and C-epitope pools of gal-3, fixed cells were surface incubated with the respective antibodies for 45 minutes without permeabilization. For simultaneous visualization of both epitopes on the same tissue section, primary antibodies (N-epitope-FITC conjugated, C-epitope Alexa 594 conjugated) were applied sequentially, and signals were developed. For visualization of surface-specific signals from secreted, surface-bound fluorescently tagged EGFP-Gal-3 protein, anti-GFP antibody was used, followed by incubation with Alexa 594-conjugated secondary antibody on un-permeabilized fixed cells. Imaging was performed using 561nm laser for capturing signal from Alexa 594 emission.

### 2.8. Proximity Ligation assay

PLA was performed on cardiomyocytes and cardiac tissues as per the manufacturer’s instructions. To compare the phosphorylation status of protein products expressing WT-Gal-3-GFP and S6-Mut-Gal-3-GFP in cardiomyocytes under various treatments, PLA was conducted using anti-GFP and anti-pan-phospho serine (pSer) antibodies as per the manufacturer’s protocols.

### 2.9. Image analysis

Image analyses were performed as previously described [28]. For single-cell analyses, between 50 and 200 cells were quantified from five randomly selected fields per condition, across three independent experiments. Image analysis was conducted using Fiji software, and calibration bars for LUT-converted images were included in the corresponding figures.

### 2.10. Statistics

All experiments were performed in triplicate, and error bars represent the standard deviation (SD) calculated from three independent experiments. Statistical significance for comparisons was determined using t-test with Bonferroni’s correction, with significance levels indicated as *p ≤ 0.05, **p ≤ 0.01, and ***p ≤ 0.001. Data are presented as means ± SD from a minimum of three independent trials.

## 3. RESULTS

Please note that the uncropped original images of dot blots and Western blots, are available in the supplementary file1, section C. Supplementary figures-tables, detailed methodology and sources of reagents are available in supplementary file 1, section A and B respectively.

### 3.1. AR reverses left ventricle (LV) hypertrophic phenotype in rats

The Wistar rat model of biological aging and pressure-overload induced left ventricular cardiac hypertrophy (hereafter, BA and PO-CH respectively) with and without AR administration, has been extensively validated and previously published by our co-author [20, 21]. For this study, we utilized the same animal cohorts, and as a result, we have not provided previously established data on echocardiographic and molecular improvements in BA and PO-CH owing to AR multifarious effects. However, new data derived from the same cohorts **(Figure. 1 and Figs. S1–S2)** are presented to further validate the study model before undertaking a detailed analysis of Gal-3 C-epitope oligomers in cardiac hypertrophy and their attenuation by AR.

**Figure 1.**
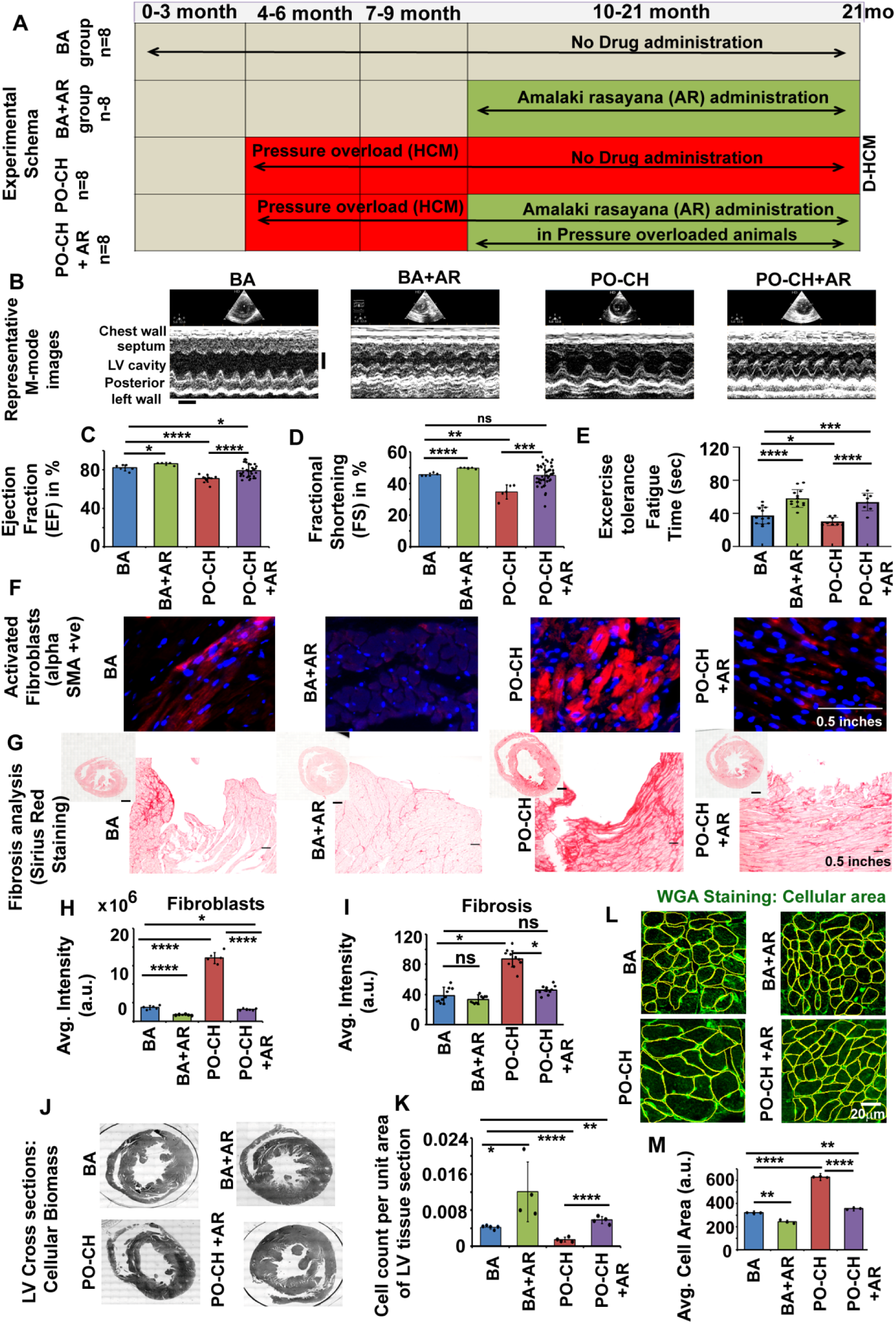
AR-based rat model for left ventricular hypertrophy. **(A)** Study design: Groups were BA (21-mo aged rats with mild CH), BA+AR (BA rats treated with AR from 9–21 months), PO-CH (TAC at 3 mo, CH by 9 mo), and PO-CH+AR (PO-CH treated with AR 9–21 mo). **(B–D)** Echocardiography (M-mode): AR-treated BA and PO-CH rats have improved LVEF, LVFS, and ventricular dimensions vs. untreated groups. Scale bar, x axis-0.4 cm; y axis 2.4 s. **(E)** Exercise tolerance (treadmill fatigue time): AR-treated rats perform better than untreated. **(F, H)** IHC for αSMA⁺ fibroblasts: AR-treated animals have significantly fewer activated fibroblasts, characterized by enlarged-swollen phenotypes. **(G, I)** Sirius Red staining: AR-treated hearts show reduced fibrosis. **(J, K)** LV wall representative cross-sections (21 mo): untreated BA/PO-CH show dilated hypertrophy; AR-treated hearts maintain ventricular wall thickness with less cell loss than untreated cohorts. **(L–M)** WGA staining for visualization of cardiomyocyte surface at the study end-point: AR-treated rats have smaller cardiomyocyte cross-sectional area (less hypertrophy) in all respective myocardial layers. Data are mean ± SD (n=4–8 rats/group); t-test: *p < 0.05, **p < 0.01, ***p < 0.001, ****p < 0.0001.

AR treatment (500 mg/kg for 12 months; **Figure 1A**-experimental schema) significantly improved left ventricular ejection fraction (EF) and fractional shortening (FS), which were notably higher than in untreated BA and PO-CH rats **(Figure 1B-D)**. Treadmill tests also showed AR-treated rats ran significantly longer than untreated counterparts **(Figure 1E)**. Histological analysis revealed fewer activated fibroblasts **(Figure 1F, H) and** macrophages **(Fig. S1A, C)**, with reduced collagen deposition (fibrosis, **Figure 1G, I**) and calcification (stiffness, **Fig. S1B, D**) in AR-treated hearts compared to untreated BA and PO-CH LV tissue sections.

At study completion, AR-treated rats exhibited significantly better morphology, cellular biomass (**Figure 1J,K**, cell density per unit LV cross-sectional area) and reduced cardiomyocyte cross-sectional area (reduced hypertrophy) in PO-CH compared to untreated BA as baseline control **(Figure 1L,M)**. AR-treated LV exhibited protein profiles akin to recovery from CH, with downregulation of hypertrophy-associated proteins (ANP, GATA4, MYH7, MEF2A, CAMK2D, CnA, NFATC2) and upregulation of those linked to normal LV function (cTNT, ACTN2, TMOD1, OBSCN, SERC2A) [**Fig. S2**, for detailed functions of the examined proteins, refer to **supplementary Table S1**]. These animal model re-analyses confirm AR’s efficacy in reversing maladaptive cardiac remodelling in both aging and pressure-overload CH models.

### 3.2. Galectin-3 C-epitope oligomers show extracellular accumulation in hypertrophied BA and PO-CH sera and tissues and are reduced by AR

Untreated hypertrophic rats (both BA and PO-CH models) exhibited elevated serum levels of total Gal-3 (i.e. full length, N- and C-epitopes) in dot blots **(Figure 2A–B)**. However, N- and C-epitope-specific Western blot analysis of PO-CH samples revealed a preferential enrichment of C-terminal oligomers **(Figure 2C–D)**. Further Western blot analysis of serum, across experimental groups, confirmed a significant increase in Gal-3 C-epitope enriched oligomers in both PO-CH and BA groups, which were reduced following Amalaki rasayana (AR) treatment **(Figure 2E–F)**. Quantitative densitometry showed that serum Gal-3 C-epitope oligomer levels were approximately twofold higher in untreated PO-CH compared to BA animals (p < 0.01), with AR-treated values returning close to BA baseline.

**Figure 2.**
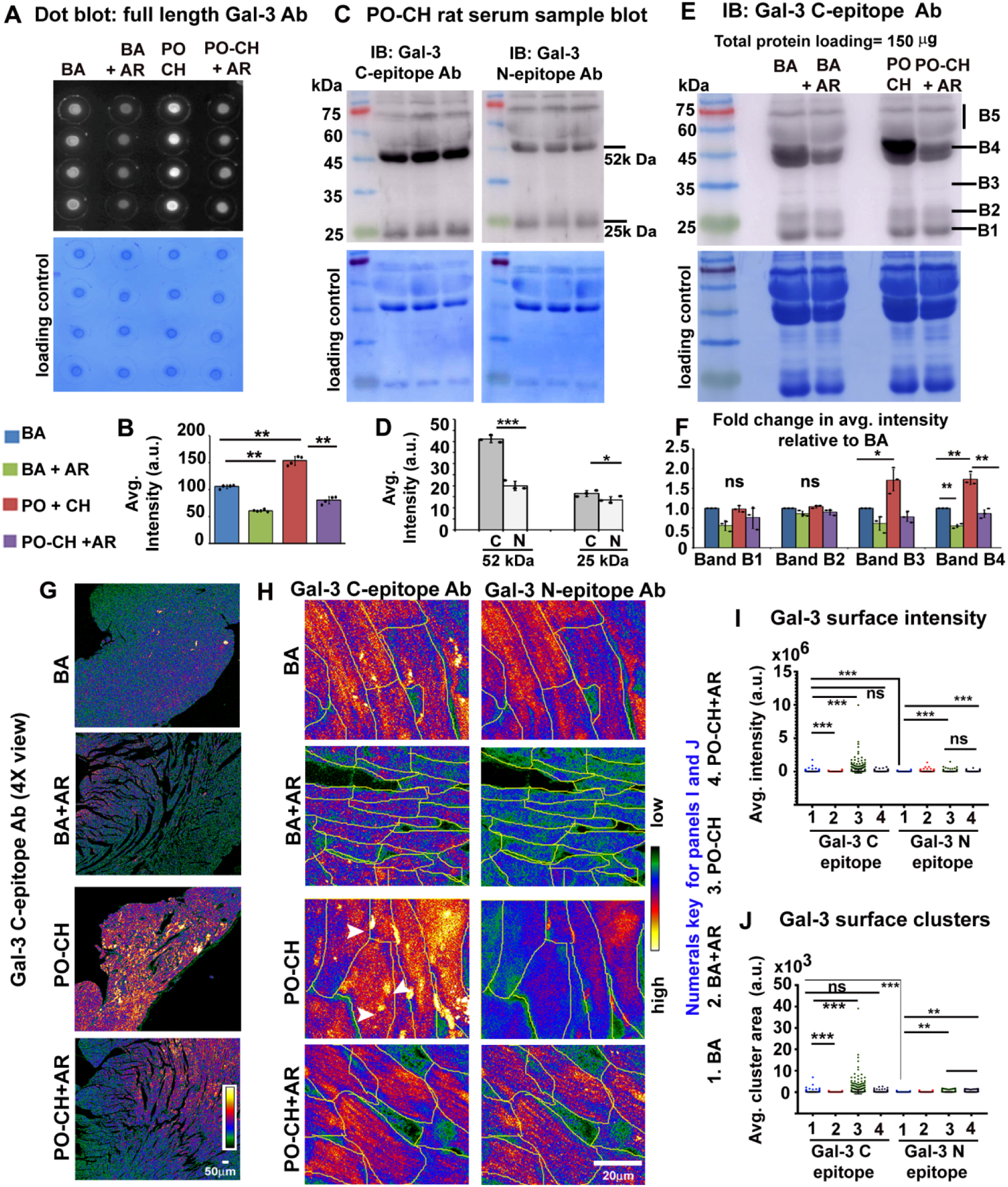
AR treatment reduces excess Gal-3 C-epitope oligomers in CH. **(A, B)** Dot blot assay (anti-Gal-3 aa 1–250) of serum from BA, BA+AR, PO-CH, and PO-CH+AR rats (n=4 per group; 6 µg serum/spot). KO-validated antibodies were used for signal detection. **(C, D)** Western blots with epitope-specific antibodies (60 µg PO-CH serum) show enrichment of Gal-3 in full length (26kDa),Gal-3 C-epitope homo-dimers (21-25 kDa, as monomer is 10.5 kDa) and oligomers (52 kDa and above). Note: C-epitope is present in C–C homo-oligomers and full-length forms and N-epitope is present in N–N homo-oligomers and full-length forms. Antibodies validated with blocking peptides were used for signal detection **(E, F)** Westerns confirm AR-treated BA and PO-CH rats have altered levels of oligomeric Gal-3 C-epitope. **(G)** Representative images (4×) of Gal-3 C-epitope clusters on cardiomyocyte membranes in left ventricular tissue: BA vs. PO-CH (PO-CH shows prominent clustering, see white arrowheads). **(H–J)** Quantification of surface Gal-3 N- vs. C-epitope in BA and PO-CH hearts (with/without AR); untreated PO-CH shows significantly more C-epitope clustering. Tissue sections are un-permeabilized to reveal surface staining. Data are mean ± SD (n=4 rats/group); t-test: *p ≤ 0.05, **p ≤ 0.01, ***p ≤ 0.001. Note for panel I,J: Conditions in which data values are tightly clustered around the mean appear as a straight line overlapping the mean.

Consistent with the serum profile, left ventricular (LV) tissue sections showed markedly higher Gal-3 C-epitope oligomer accumulation in PO-CH and moderately elevated levels in BA, distributed across multiple cardiac cell types and in the extracellular matrix **(Figure 2G)**. Please note that tissue sections were not detergent-permeabilized to ensure detection of only surface-localized epitopes. Cardiomyocytes from PO-CH tissue sections displayed prominent surface-associated C-epitope oligomers, which were notably stronger than those of N-epitope, indicating a predominant plasma membrane association of C-terminal Gal-3 oligomers during hypertrophy **(Figure 2H-J)**. Notably, the surface oligomer levels were substantially reduced in the corresponding AR-treated groups **(Figure 2J)**. In contrast, BA cohorts exhibited relatively fewer surface-bound C-epitope oligomers vs PO-CH, which were nearly absent following AR treatment. Restoration of Gal-3 C-epitope levels correlated with improved ventricular function, suggesting a potential causal association.

### 3.3 Galectin-3 C-epitope extracellular oligomers cause cardiomyocytes hypertrophy which is reversed by AR

Since Gal-3 C-epitope oligomers were abundantly localized on the surface of hypertrophied cardiomyocytes, we next investigated whether these extracellular oligomers could directly induce hypertrophy. For this, cardiomyocytes were incubated with conditioned media enriched in fluorescently-tagged wild-type or mutant Gal-3 C-epitope.

HEK-293 cells expressing EGFP wild-type Gal-3 C-epitope with intact CRD [carbohydrate recognition domain, hereafter, Gal-3-WT-CRD-EGFP] or a mutant CRD [R186S with reduced ability to bind β-galactosides, hereafter, Gal-3-MUT-CRD-EGFP] served as sources of surface high affinity glycan binding and low affinity glycan-binding C-epitopes, respectively [27]. To simulate stress, one set of cells was exposed to pressure-overload (PO) using a mechanical weight application method **(Figure 3A- experimental schema) [**23, 24], while another remained untreated (No PO). Conditioned media collected after 24 h were applied to cardiomyocyte cell line (rat left ventricular cardiomyoblasts cell line, H9C2, differentiated to cardiomyocytes) for 10 h before hypertrophy analysis.

**Figure 3.**
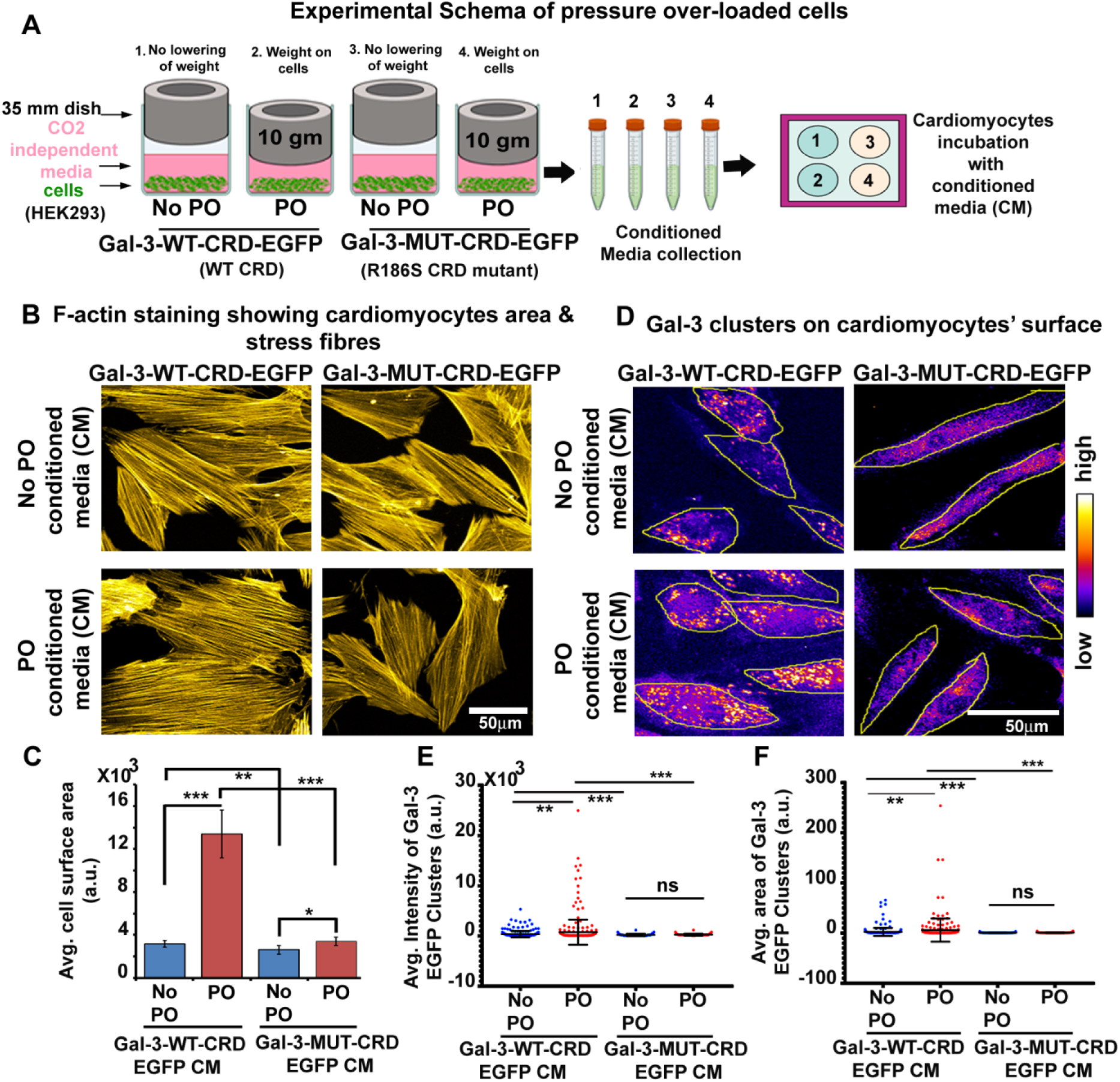
Predominance of Gal-3 C-epitope oligomers in cardiomyocyte hypertrophy. **(A)** Schematic of the *in vitro* pressure-overload (PO) model. **(B, C)** HEK293 cells transfected with Gal-3 C-epitope WT–CRD-EGFP or CRD-mutant (R186S) were cultured under No-PO and PO conditions. Conditioned media (CM) were collected and applied to fresh cardiomyocytes for 24 h; cells were fixed and stained with rhodamine-phalloidin (F-actin) to expose cell periphery for cell surface area analyses (hypertrophy measurement) **(D–F)** Anti-GFP antibody was incubated on the detergent un-permeabilized cell surface and signal was developed with Alexa 594 conjugated secondary antibody, to expose surface specific signal from Gal-3 WT and MUT CRD. Quantification of cardiomyocyte surface-bound Gal-3 C-epitope: surface binding-signal intensity and surface cluster area of WT-CRD-EGFP vs. CRD-MUT-EGFP under No-PO vs. PO. Data are mean ± SD; t-test: *p ≤ 0.05, **p ≤ 0.01, ***p ≤ 0.001. **(E-F**) Note: Conditions in which data values are tightly clustered around the mean appear as a straight line overlapping the mean.

Cardiomyocytes treated with Gal-3-MUT-CRD-EGFP condition media from PO and No-PO set-ups showed significantly reduced hypertrophic enlargement compared with those treated with Gal-3-WT-CRD-EGFP under the same conditions **(Figure 3B, C)**. Surface GFP signal imaging (via detection with anti-GFP primary ab and Alexa 594 conjugated secondary ab) confirmed markedly weaker binding of Gal-3-MUT-CRD-EGFP, corresponding to its diminished hypertrophic effect **(Figure 3D, E)**. Notably, Gal-3-WT-CRD-EGFP from PO-conditioned media formed distinct micron-sized surface clusters on cardiomyocytes, whereas Gal-3-MUT-CRD-EGFP showed diffuse staining, underscoring the essential role of the C-epitope CRD in oligomerization and surface membrane clustering **(Figure 3D, F)**.

As detailed earlier, in our rodent models of BA and PO-CH, AR treatment markedly reduced activated macrophages and fibroblasts—the primary sources of extracellular Gal-3, while cardiomyocyte density was restored. Therefore, we wanted to assess whether AR negatively influences excess Gal-3 secretion, C-epitope formation, and its surface binding in cardiomyocytes to enable cardiac viability. In this pursuit, we exposed differentiated cardiomyocytes (H9C2) to 24 hours of pressure-overload (PO) stress, followed by 100 µg/mL AR treatment for an additional 24 hours. The effective dosage of AR was validated through MTT viability and cell count assays, as shown in **Figures 4A-B**. The AR mediated reduction in hypertrophy was assessed through bright-field images **(Figures 4C)**.

**Figure 4.**
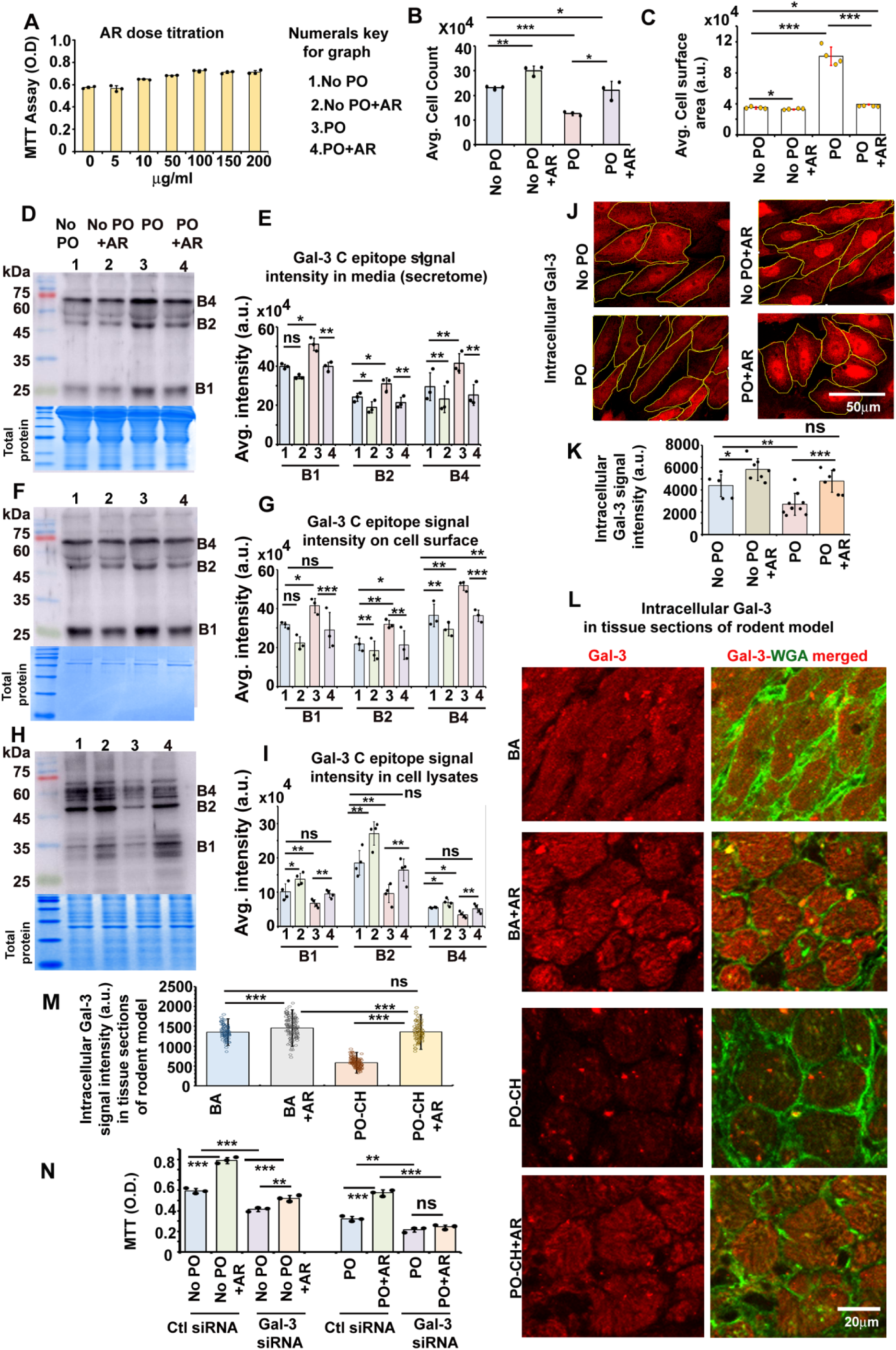
AR reduces extracellular Gal-3 C-epitope oligomers on cardiomyocytes *in vitro*. **(A)** MTT assay of cardiomyocyte (H9C2) viability with increasing AR doses (24 h); no viability gain above 100 µg/mL. **(B)** Cardiomyocytes under 24 h PO treated with 100 µg/mL AR for 24 h had higher cell counts than untreated controls. **(C)** Cardiomyocytes under 24 h PO treated with 100 µg/mL AR for 24 h had lower cell surface area (hypertrophy) than untreated controls. **(D–I)** Western blots of Gal-3 C-epitope dimers/oligomers in media (D, E), surface fractions (F,G), and cell lysates (H, I) under different experimental conditions (surface Gal-3 isolated by low-pH wash). **(J, K)** Confocal images analyses show that cardiomyocytes under 24 h PO treated with 100 µg/mL AR for 24 h had higher Gal-3 intracellular retention than untreated controls. **(L, M)** Confocal images analyses show that AR treated BA and PO-CH LV cardiomyocytes had higher Gal-3 intracellular retention than untreated controls. **(N)** MTT assay performed in cardiomyocytes under AR± and No PO/PO conditions with Gal-3 siRNA or scramble siRNA (labelled as Ctl siRNA) background*. Important note:* Complete siRNA-mediated ablation of Gal-3 is neither feasible nor physiologically relevant, given its essential intra- and extracellular functions,; t-test: *p ≤ 0.05, **p ≤ 0.01, ***p ≤ 0.001.

Correlative to enhanced viability and rescue from hypertrophy, Western blot analysis of culture media revealed a marked reduction in secreted Gal-3 C-epitope oligomers in AR-treated cells under both No PO and PO conditions compared to untreated controls **(Figures 4D, E)**. Correspondingly, surface-bound C-epitope signal intensity was significantly decreased following AR treatment **(Figures 4F, G)**. Notably, these observations are akin to those observed in the *in vivo* rodent model LV tissue cross-sections (refer to Figure 2H-I). Western blot analysis showed increased intracellular Gal-3 levels in AR-treated groups **(Figures 4H, I)**, indicating that AR may inhibit excessive Gal-3 secretion by altering its intracellular trafficking or post-translational modification, thereby limiting its extracellular accumulation [29, 30]. Intracellular retention was further confirmed at an *in vitro* cellular level and in tissues sections from the *in vivo* rodent model **(Figures 4 J-M)**. Importantly, MTT assay results indicated that AR treatment under both No-PO and PO conditions, in a Gal-3–depleted background (60–65% downregulation by siRNA), significantly reduced cell viability, highlighting the critical role of intracellular Gal-3 in maintaining cardiomyocyte survival **(Figure 4N)**.

### 3.4 Gallic acid in AR binds Gal-3 CRD and promotes intracellular retention

Amalaki rasayana (AR) is predominantly enriched in a bioactive component gallic acid (GA) [19, 20, 21, 22]. Amongst eighteen components of AR, GA has been well documented for its cardioprotective role in rodent pressure-overload model. In earlier pharmacokinetic data, GA was also identified as a major circulating component in AR-treated rodent serum [21]. Therefore, through next sets of experiments, described in sections 3.4-3.6, we wanted to examine GA as a causal component of AR in negatively regulating C-epitope oligomer formation, surface binding and hypertrophy.

Evidences show that extracellular Gal-3 C-terminal epitope oligomers primarily form through protease-mediated cleavage of the secreted full-length Gal-3 protein. Notably, MMP-2, MMP-9 proteases known to be secreted by stressed cardiomyocytes, fibroblasts, macrophages are implicated in this cleavage process, but is inhibited by GA [14, 17, 31, 32].

In cultured cardiomyocytes, GA treatment (100 µM) reproduced AR’s effects—significantly reducing extracellular and surface-bound Gal-3 C-epitope oligomers while increasing intracellular Gal-3 levels **(Figure 5A–K)**. Preliminary dose–response experiments indicated that 100 µM GA produced maximal efficacy, but in Gal-3 depleted cardiomyocytes, the same concentration was less effective, indicating Gal-3 intracellular retention as one of the dominant anti-apoptotic function of both GA and AR **(Figure 5L)**.

**Figure 5.**
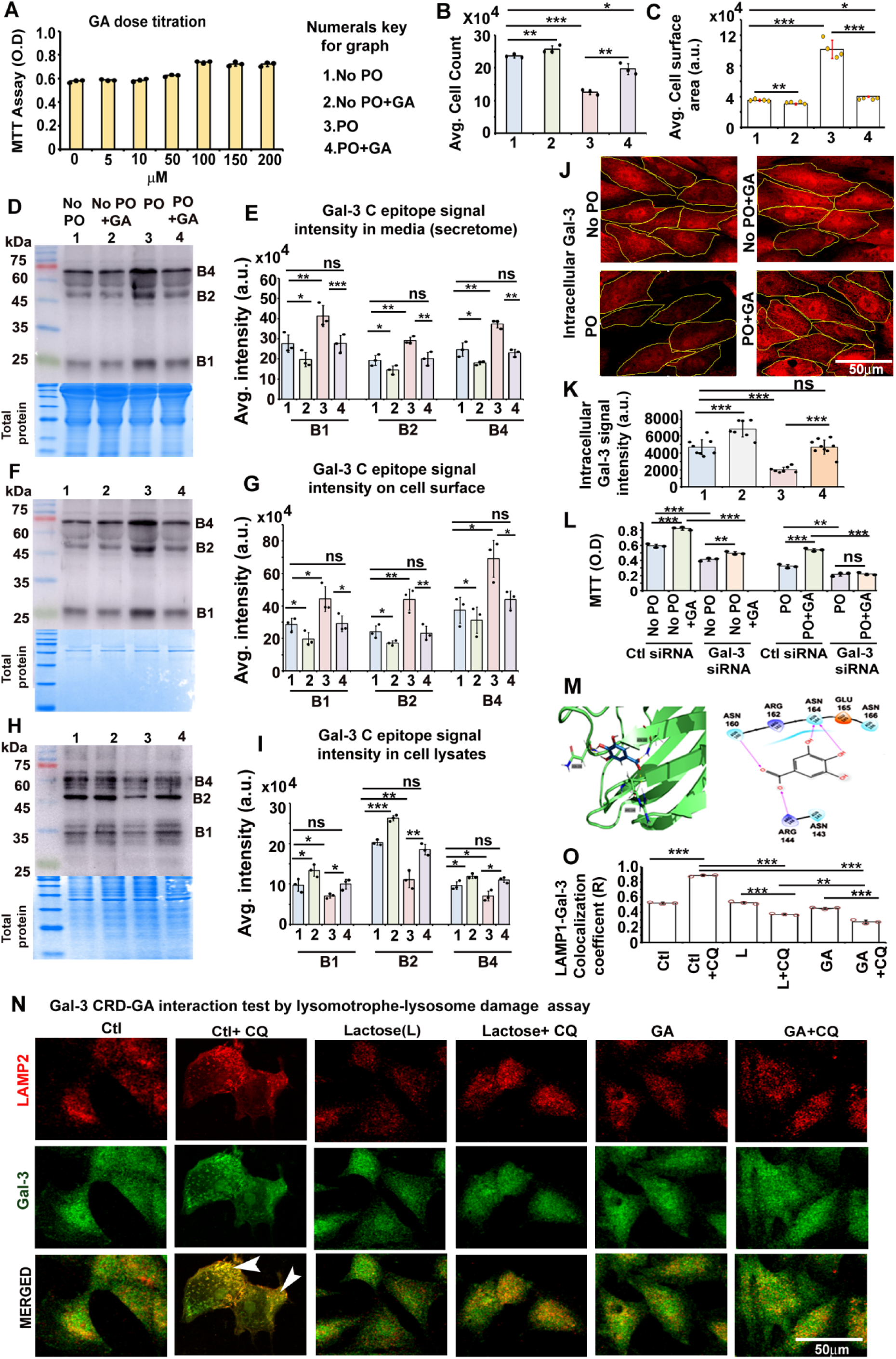
Gallic acid (GA) from AR reduces extracellular Gal-3 C-epitope oligomers. **(A)** MTT assay of cardiomyocytes with increasing GA concentrations (24 h): no viability increase above 100 µM. **(B)** Cardiomyocytes under 24 h PO treated with 100 µM GA for 24 h had higher cell counts than controls. **(C)** Cardiomyocytes under 24 h PO treated with 100 µM for 24 h had lower cell surface area (hypertrophy) than untreated controls. **(D–I)** Western blots of Gal-3 C-epitope in media (D, E), surface (F, G), and lysates (H, I) under different experimental conditions. **(J)** MTT assay performed in cardiomyocytes under GA± and No PO/PO conditions with Gal-3 siRNA or scramble siRNA (labelled as Ctl siRNA) background. **(K, L)** Molecular Docking: GA forms hydrogen bonds with Gal-3 CRD residues Arg144, Asn160, Asn164. **(M)** In the absence of GA pre-treatment, CQ mediated exposure of lysosomal inner membrane glycans leads to marked accumulation of Gal-3 (red) at LAMP1⁺ lysosomes (green) in cardiomyocytes (white arrowheads points to such colocalized areas that appear yellow), suggesting that GA interacts with the Gal-3 carbohydrate recognition domain (CRD) to block its glycan-binding activity. Data are mean ± SD; t-test: *p ≤ 0.05, **p ≤ 0.01, ***p ≤ 0.001.

A previous computational modelling study on a GA derivative-C-glycoside of 4-O-methylgallic acid binding to Gal-3 showed strong and exclusive involvement of CRD [33]. Our basic molecular docking analyses showed that GA forms hydrogen and electrostatic bonds with key Gal-3 C-epitope CRD residues (Arg144, Asn160, Asn164) **(Figure 5M)** further relevance of these residues is demonstrated in section 3.5). This binding conformation was associated with a Glide docking score of –6.191 and an MM-GBSA ΔG bind of –14.35 kcal/mol, indicating a stable interaction and energetically favorable interaction between GA and Gal-3. To experimentally verify GA binding to the CRD, we conducted a well known Gal-3 CRD-dependent intracellular function inhibition assay. Report show that upon chemically-induced lysosomal damage [via use of a lysomotrophe- chloroquione (CQ), LLOMe, GPN etc] [34, 35], Gal-3 rapidly accumulates around damaged lysosomes and binds to exposed glycans on their membranes via CRD, unless its CRD is blocked with an inhibitor. To examine the potential of GA as a Gal-3 CRD blocker, differentiated H9C2 cells were treated with 50 µM CQ for 24 hours and lysosomal-associated membrane protein (LAMP1)-full length-Gal-3 protein co-localization were probed. Conditions with undamaged lysosomes showed no Gal-3-LAMP1 accumulation **(Figure 5N-O**, panel ‘O’ indicates Mander’s colocalization coefficient). However, GA pre-treatment of cells, before CQ incubation, reduced Gal-3 binding to damaged lysosomes, indicating inhibition of Gal-3’s glycan binding activity. Although, this inhibition does not cause cell toxicity as other factors are known to overtake Gal-3 lysosomal repair functions. Collectively, these findings suggest that GA directly binds to the Gal-3 CRD, promoting its physiologically needed intracellular retention and thereby limiting Gal-3 secretion, extracellular Gal-3 C-epitope oligomer formation and accumulation.

### 3.5. GA and AR promote Gal-3 Ser6 phosphorylation *in vitro* and *in vivo* for intracellular retention

The next step was to elucidate the major mechanism by which AR and GA promote intracellular retention of Gal-3. High intracellular calcium is a well-established trigger for Gal-3 exocytosis [29]. Treatment of cardiomyocytes with gallic acid (GA) or Amalaki rasayana (AR) resulted in a reduction of surface-bound Gal-3 C-epitope oligomers comparable to that observed with BAPTA-AM, a calcium chelator **(Figure 6A–C)**. Calcium imaging further revealed that both AR and GA attenuate intracellular calcium levels **(Figure 6D, E)**. These findings suggest that GA and AR may inhibit Gal-3 secretion by lowering intracellular calcium. Consequently, AR and GA help sustain the intracellular Gal-3 pool, preserving its physiological roles while preventing the formation of pathological extracellular C-epitope oligomers. Supporting this notion, previous studies have shown that GA decreases cytosolic calcium by enhancing its reuptake into the endoplasmic reticulum through SERCA2 activation, while also inhibiting calcium-dependent kinases such as CaMK2A, which otherwise contribute to elevated intracellular calcium levels [36,37].

**Figure 6.**
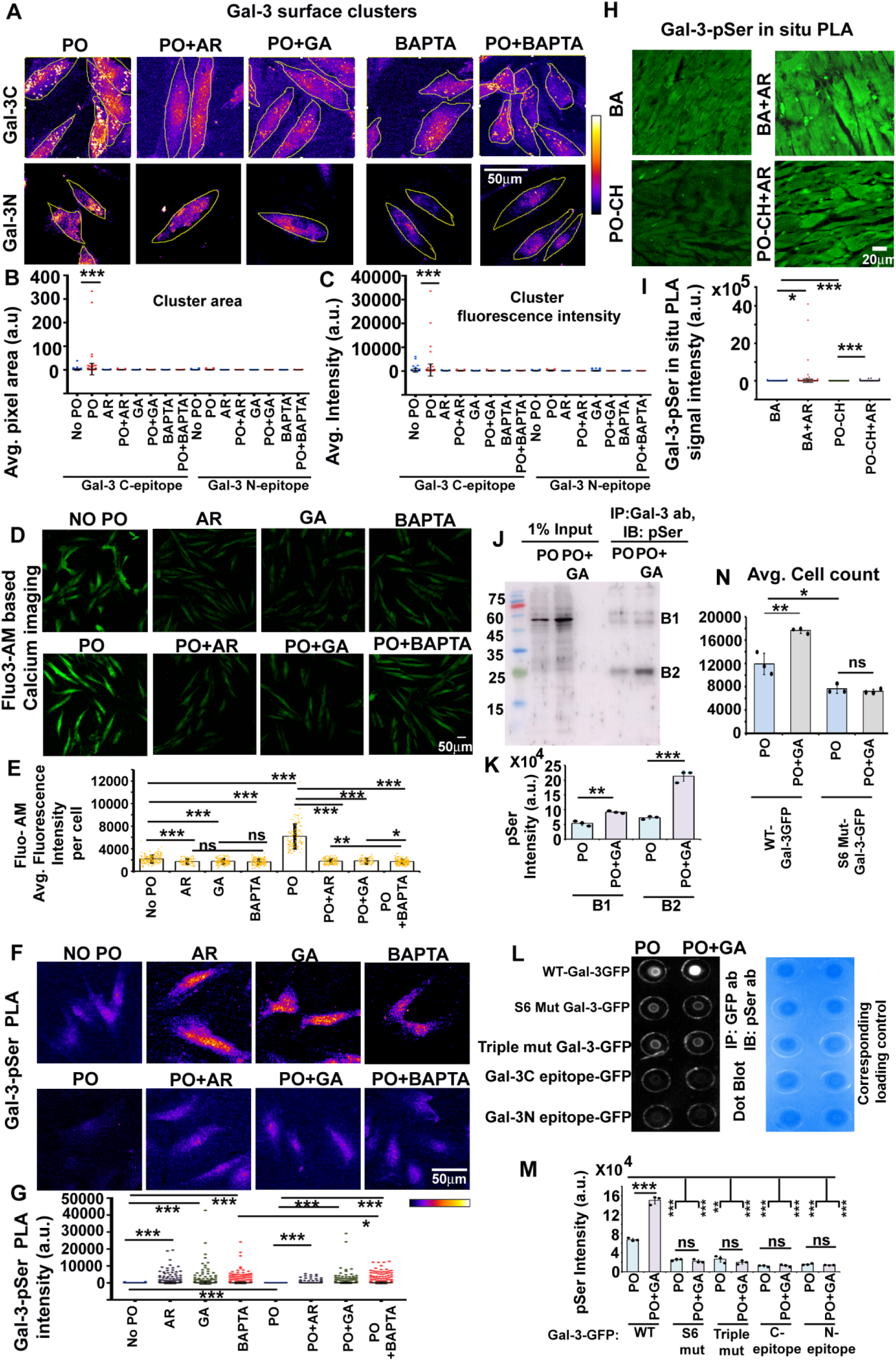
GA in AR enhances Gal-3 N-epitope phosphorylation and Gal-3 retention. **(A–C)** GA, AR, and BAPTA-AM treatments each reduce the surface cluster area and intensity of extracellular Gal-3 C-epitope on cardiomyocytes (No-PO and PO conditions). **(D, E)** GA, AR, and BAPTA-AM treatments each reduce the intracellular calcium in cardiomyocytes (No-PO and PO conditions). **(F, G)** Proximity ligation assay (PLA) shows AR/GA increase Gal-3 phosphorylation and intracellular retention; BAPTA-AM (Ca²⁺ chelator) has similar effects, implicating Ca²⁺ in Gal-3 secretion. **(H, I)** PLA quantification in rat hearts: untreated PO-CH tissues have significantly lower phosphorylated Gal-3 than AR-treated hearts. **(J, K)** IP (anti–Gal-3), IB: Anti-pan-phosphoserine Western: GA-treated cardiomyocytes show increased intracellular Gal-3 serine phosphorylation in PO condition. **(L, M)** IP of GFP-Gal-3 from PO ± GA cells: GA treatment enhances serine phosphorylation of WT-Gal-3. The S6 mutant shows a weak signal with no change under PO ± GA conditions, confirming Ser6 as a major phosphorylation site regulated by GA. Similarly, the triple-CRD mutant, which cannot bind GA, shows a weak, unchanged signal, indicating that GA–CRD interaction is required for phosphorylation. Lysates expressing only the N- or C-epitope show minimal signal, indicating a lack of serine phosphorylation and underscoring the dependency between GA binding and Ser6 phosphorylation. These findings suggest that GA binds to the C-epitope CRD, triggering a conformational shift in the N-epitope that facilitates phosphorylation at Ser6. **(N)** PO ± GA conditions expressing Ser6 mutant had lower cardiomyocyte cell count vs WT-GFP-Gal-3, indicating GA mediated Gal-3 phosphorylation is cytoprotective. Data are mean ± SD (n ≥ 4 rats or 3 independent experiments); t-test: *p ≤ 0.05, **p ≤ 0.01, ***p ≤ 0.001. Note: Conditions in which data values are tightly clustered around the mean appear as a straight line overlapping the mean.

Reports suggest that phosphorylation of the Ser6 residue in the N-terminal domain of Gal-3, catalyzed by casein kinase I (CKI), favors its cytoplasmic localization over extracellular release [38]. This phosphorylation is strongly associated with the intracellular anti-apoptotic activity of Gal-3 [39]. However, elevated cytoplasmic calcium, levels in CH activate calcineurin, which dephosphorylates and inactivates CKI [40], thereby can potentially reduce Gal-3 Ser6 phosphorylation. Since GA attenuates intracellular calcium elevation, it may indirectly inhibit calcineurin activation, thereby preserving CKI activity, supporting Gal-3 Ser6 phosphorylation. In addition, protein phosphatase 1 (PP1) is known to dephosphorylate Gal-3 Ser6, and GA/GA-derivatives has been reported to directly inhibit PP1 [38, 41]. Thus, GA can act as a positive modulator of Gal-3 Ser6 phosphorylation through dual regulation of calcineurin and PP1.

Because antibodies specific to phospho-Ser6-Gal-3 or pan-phospho-Ser-Gal-3 are currently unavailable, to obtain a preliminary evidence of GA mediated Gal-3 serine phosphorylation, we employed a proximity ligation assay (PLA) using Gal-3 and pan-phosphoserine antibodies. AR- or GA-treated cardiomyocytes showed markedly higher Gal-3 serine phosphorylation compared to controls. Similarly, treatment with the calcium chelator BAPTA-AM increased Gal-3 serine phosphorylation, supporting a calcium-dependent regulatory mechanism mediated by GA **(Figure 6F, G)**. In vivo, hypertrophic rat hearts treated with AR displayed approximately two- to three-fold higher Gal-3-phosphoserine PLA signal levels than untreated BA and PO-CH hearts, respectively **(Figure 6H, I)**.

Further complementary experiments were conducted to support observations obtained through the PLA assay. Immunoprecipitation of cardiomyocyte lysates from PO ± GA conditions using anti-Gal-3 antibodies followed by detection with anti-pan-phosphoserine revealed increased intracellular Gal-3 serine phosphorylation specifically under PO + GA conditions **(Figure 6J,K)**. Mechanistic validation using cardiomyocytes overexpressing Gal-3 with a mutated CRD (Arg144, Asn160, Asn164 triple mutant), lacking GA binding sites, demonstrated that an inhibition of GA-Gal-3 CRD binding failed to induce Gal-3 serine phosphorylation **(Figure 6L, M)**. Likewise, cells over-expressing Ser6Ala Gal-3 mutants showed no phosphorylation response even in presence of GA. These findings indicate that GA binding to the Gal-3 C-epitope CRD facilitates exposure of the N-epitope Ser6 site to CKI, promoting its phosphorylation and cytoplasmic retention. IP with lysates over-expressing C-epitope alone and N-epitope alone constructs and IB with phosphoserine ab, revealed very weak and similar signal profile in PO and PO+GA condition, again suggesting the C-epitope is not serine phosphorylated by itself but it is needed for N-epitope phosphorylation. Overall, mutational analyses demonstrated that GA-induced phosphorylation requires an intact Gal-3 CRD and the Ser6 site, as neither CRD mutants nor Ser6Ala variants responded to GA. These results also indicate that GA binding to the Gal-3 CRD promotes exposure and phosphorylation of Ser6 by CKI, facilitating cytoplasmic retention and reducing extracellular oligomer formation to alleviate, cardiomyocytes’ death. Indeed, cell counts were far reduced in PO±GA cardiomyocyte cultures expressing Ser6 Gal-3 mutant vs the PO+GA WT-Gal-3 condition, highlighting the cytoprotective role of GA via Ser6 Gal-3 phosphorylation **(Figure 6N)** .

### 3.3. Gal-3 C-epitope oligomers show accumulation on the surface of human hypertrophic myocardium

To validate our *in vitro* cell culture and *in vivo* rodent observations in a human context and gain deeper insights into the pathological effects of the Gal-3 C- epitope compared to the N-epitope, we conducted a series of experiments outlined in sections 3.3–3.6. Tissue samples from human cardiac hypertrophy (CH) revealed distinct surface staining patterns for the N-and C-terminal galectin-3 epitopes **(Figure 7A, B)**. In hypertrophic cardiomyocytes from patients with hypertrophic cardiomyopathy (HCM), there was a significant reduction in the co-localization of the N- and C-epitopes compared to the normal tissue sections. This indicates that C-C and N-N homo-interactions dominated over C-N or full-length galectin-3 interactions. Images exhibited prominent clusters of the Gal-3 C-epitope on the surface than the N-epitope. HCM cardiomyocytes show reduced intracellular and phosphorylated Gal-3 vs. normals **(Figure 7C–E)**. Thus, human myocardium mirrors the rodent findings: excess Gal-3 C-epitope accumulates on hypertrophic hearts and intracellular pool, required for physiological functions is reduced.

**Figure 7.**
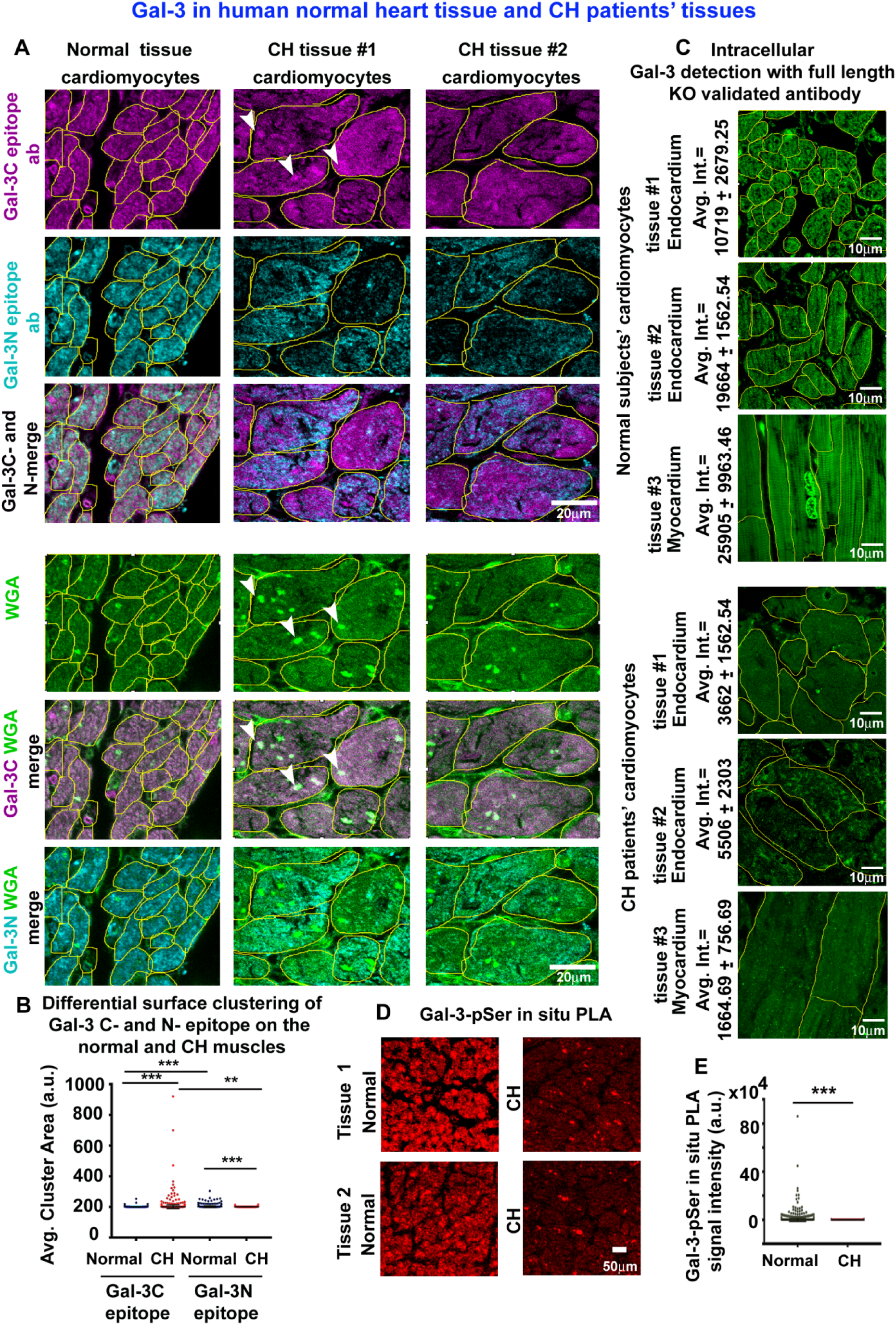
Elevated Gal-3 C-epitope on CH cardiomyocytes and reduction in intracellular retention. **(A, B)** Normal vs. CH human hearts stained for Gal-3 N-epitope (cyan) and C-epitope (magenta); WGA (green) marks surface membranes. Tissues are not permeabilized for surface specific staining. CH cells show reduced N–C co-localization and prominent C-epitope clusters (white arrowheads), indicating C-epitope oligomer dominance. **(C)** Detergent permeabilized human tissue reveals intracellular Gal-3 staining, which is significantly reduced CH cardiomyocytes **(D, E)** Gal-3-and phosphoserine PLA on human LV tissues: HCM cardiomyocytes show reduced phosphorylated Gal-3 vs. normals. Data are mean ± SD; t-test: *p ≤ 0.05, **p ≤ 0.01, ***p ≤ 0.001. Note: Conditions in which data values are tightly clustered around the mean appear as a straight line overlapping the mean.

### 3.6. Extracellular Gal-3C oligomers induce hypertrophic signalling

Next, we performed an analysis of Gal-3 C-epitope oligomers in human serum samples from three distinct groups: normal young individuals, normal elderly individuals, and survivors of cardiac hypertrophy that were under treatments (CH). Alongside this, we assessed the levels of Atrial Natriuretic Peptide (ANP), a secreted biomarker that reflects cardiomyocyte stress [42], to explore potential correlations between Gal-3 oligomers and cardiac stress **(Figure 8A, C).** CH survivors had slightly lower levels of C-epitope oligomers compared to normal age-matched controls, suggesting treatment reduced cardiac strain. Young donors with low ANP also exhibited lower C-epitope oligomer levels **(Figure 8B, D)**. Interestingly, total Gal-3 levels, measured by ELISA, did not correlate with the oligomer profile, as CH survivors had the highest total Gal-3 levels among all groups **(Figure 8E)**. This discrepancy highlights the limitations of full-length Gal-3 ELISAs, which may not detect C-epitope oligomers, thus obscuring Gal-3’s potential as a biomarker of treatment response in CH.

**Figure 8.**
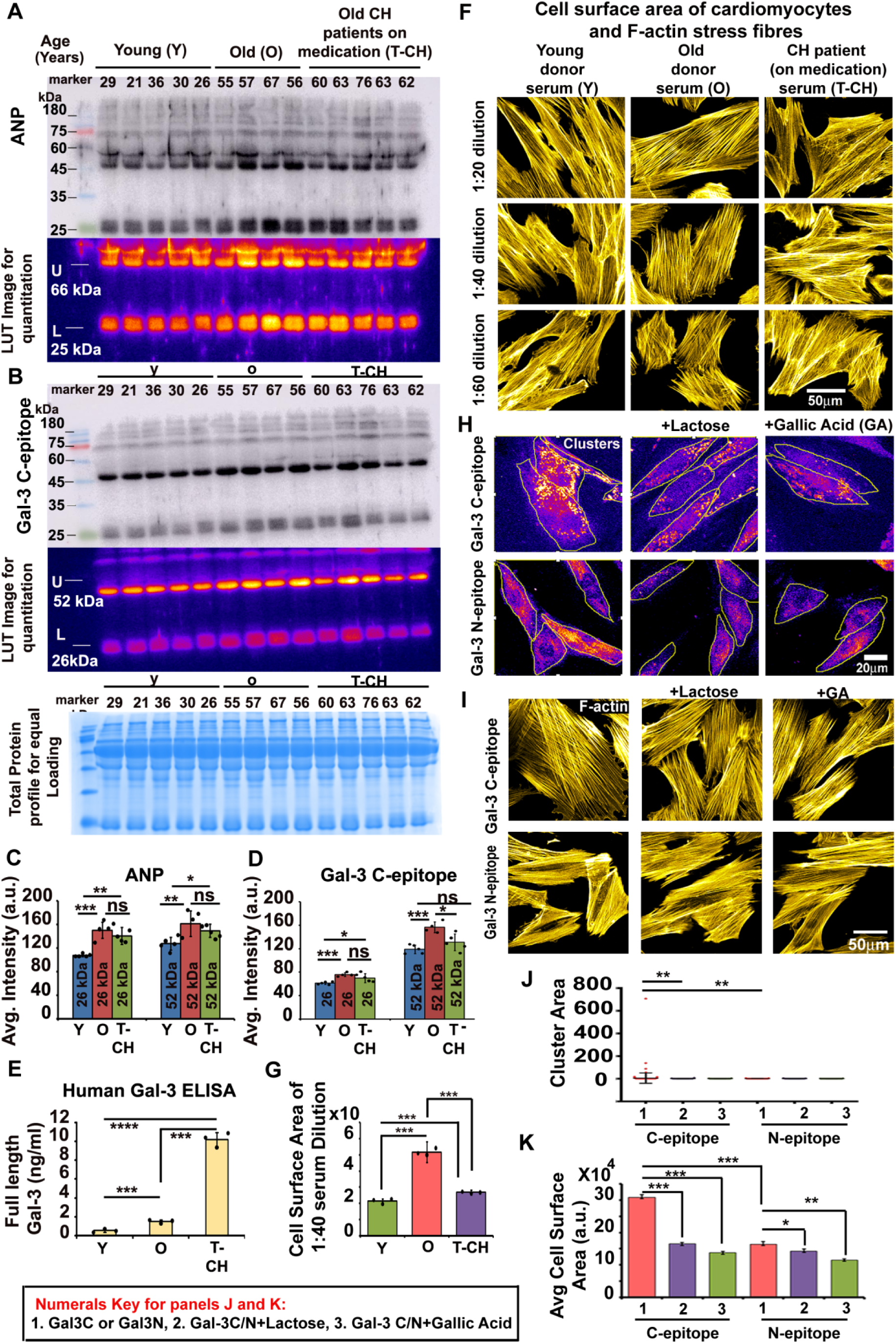
Dominance of Gal-3 C-epitope oligomers in human sera and CH cardiomyocytes vs. N-epitope. **(A-D)** ANP and Gal-3 C-epitope levels in serum from normal young, older, and clinically treated CH donors. **(E)** Total Gal-3 in the same sera measured by ELISA (full-length Gal-3 antibody). Data are mean ± SD; t-test: *p ≤ 0.05, **p ≤ 0.01, ***p ≤ 0.001, ****p ≤ 0.0001. **(F,G)** Rat cardiomyocytes treated with serum from young, old, and treated CH donors were stained with rhodamine-phalloidin. Older-donor sera induced larger cell area (hypertrophy) and more F-actin stress fibers. **(H, J)** Old-donor human sera were immunoprecipitated to isolate C- or N-epitope Gal-3 pools. Fractions were further incubated with lactose/GA or not. Immunofluorescence: Antibody mediated exposure of N- and C-epitope surface binding and clustering; C-epitope forms distinct membrane clusters (absent with lactose/GA); N-epitope is diffuse. Lactose and GA disrupts C-epitope clustering. Antibody was incubated on the un-permeabilized cells. Note: Conditions in which data values are tightly clustered around the mean appear as a straight line overlapping the mean. **(I, K)** Old-donor human sera were immunoprecipitated to isolate C- or N-epitope Gal-3 pools. Fractions were further incubated with lactose/GA or not. C-epitope–enriched serum induced significant hypertrophy (stress fibers, increased cell area), while lactose or GA blocked this effect; N-epitope pools (representing full length Gal-3 and N-N self associations) had less effect. Data are mean ± SD; t-test: *p ≤ 0.05, **p ≤ 0.01, ***p ≤ 0.001.

Although serum samples from drug-naïve CH patients were unavailable, our tissue analysis from drug naïve CH heart tissue (Figure 7) has already revealed significantly higher C-epitope oligomer accumulation on cardiomyocyte surfaces compared to controls. This suggests that reduced serum C-epitope oligomers in treated CH patients reflect therapeutic efficacy, supporting their use as biomarkers for treatment response.

To explore whether human Gal-3 C-epitope oligomers directly drive hypertrophy, we incubated cardiomyocytes with serum from older donors (high in Gal-3 C-epitope oligomers), treated CH patients (lower Gal-3 C-epitope oligomers) and young donors (lowest Gal-3 C-epitope oligomers). Cells exposed to serum from older donors hypertrophied more **(Figure 8F, G)**. Immunopurifying old donors’ sera to enrich either Gal-3 N-epitope or Gal-3 C-epitopes revealed that the Gal-3 C-epitope-enriched fraction induced significantly higher C-epitope binding, clustering and greater hypertrophy **(Figure 8H,J and Figure 8 I,K)**. Administration of Gal-3 C-epitope fraction pre-incubated with lactose (CRD inhibitor) or GA abolished this effect.

Mechanistically, Gal-3 C-epitope oligomers, but not Gal-3 N-epitope, activated the RhoA-ROCK1 stretch associated mechanotransduction pathway **(Figure 9A–C)** involved in extensive and deleterious increase in F-actin stress fibers (as shown in Figure 8I,K), and disrupted membrane tension buffering modules-the caveolae **(Figure 9D–E)** [43,44,45]. It also triggered cell stiffness associated nuclear mechanotransduction, with YAP1 translocation, increased nuclear envelop stiffness associated intermediate filament-LaminA, and elevated pγH2AX DNA damage markers **(Figure 9F–K)** [46, 47, 48]. Gal-3 C-epitope upregulated hypertrophic genes MEF2A and NFATC2 (**Figure 9L–O)** and increased membrane permeability, assessed through LDHA levels in media] **(Figure 9P-Q)**. GA or lactose treatment attenuated these effects. For detailed functional involvement of the examined signalling and structural proteins in CH pathology, refer to **supplementary Table S2**. Together, these data demonstrate that Gal-3 C-epitope oligomers bind to the cardiomyocyte surface, activating RhoA-ROCK1-mediated hypertrophic signaling, which can be blocked by inhibiting Gal-3 C-epitope glycan-binding. A graphical summary of these results highlights the role of excessive galectin-3 C-epitope oligomers as potential drug targets in cardiac hypertrophy **(Figure 9R** and **Figure 10)**.

**Figure 9.**
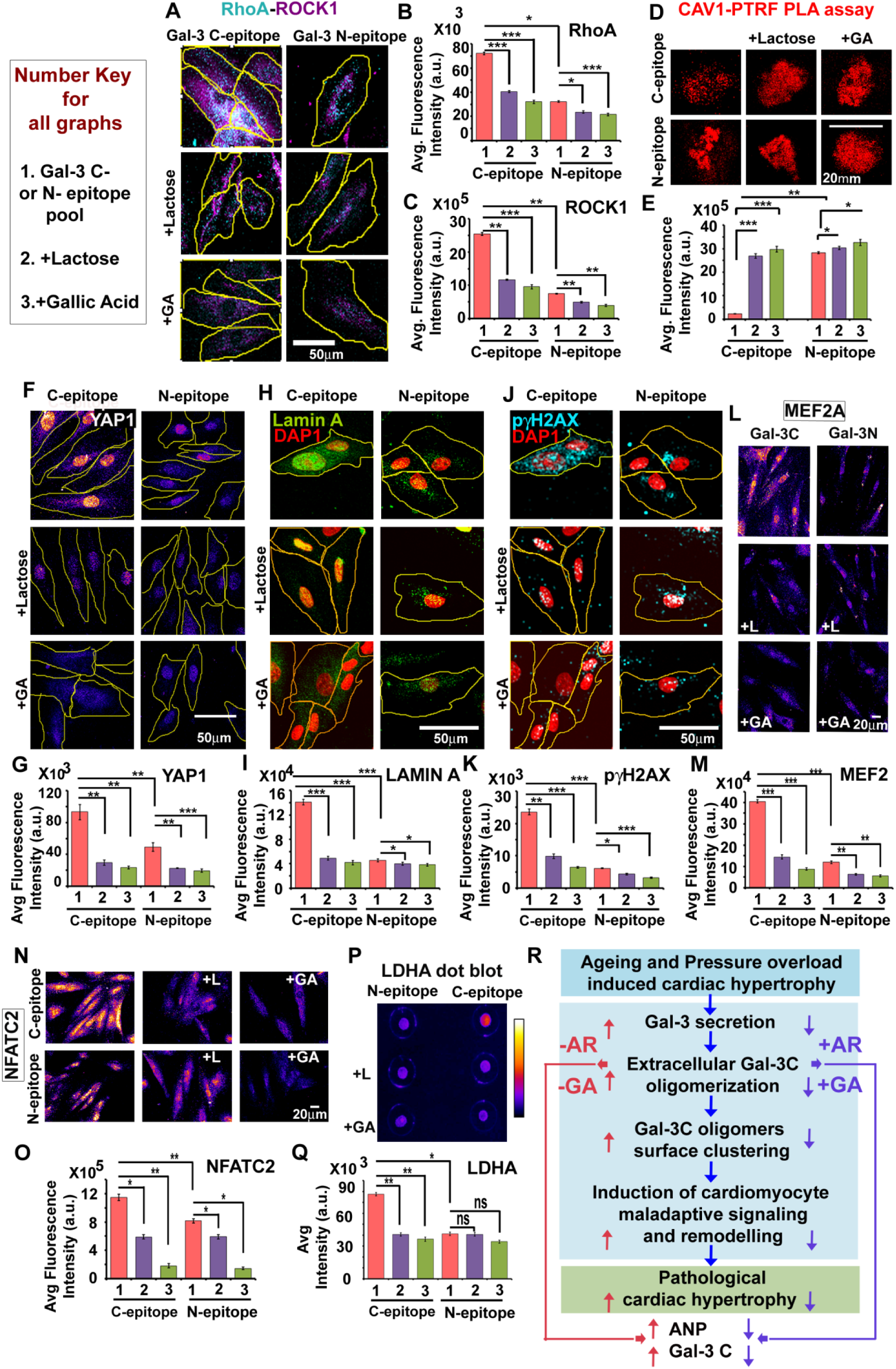
Human Gal-3 C-epitope oligomers drive cardiomyocytes’ hypertrophy and damage. Old-donor sera were used to prepare Gal-3 C- or N-epitope pools, applied to cardiomyocytes (1:40) with or without 34 mM lactose or 100 µM GA. **(A-C)** C-epitope activates RhoA–ROCK1 signaling, attenuated by lactose/GA. **(D, E)** C-epitope causes caveolae (membrane tension buffering assemblies) loss (reduced CAV1–PTRF PLA signal). **(F, G)** C-epitope increases nuclear YAP1 (indicative of surface stretch); lactose/GA reduce YAP1 nuclear signal. **(H, I)** C-epitope raises Lamin-A (nuclear envelope stiffening marker). **(J, K)** C-epitope elevates γH2AX (DNA damage marker). **(L–O)** C-epitope upregulates hypertrophy genes MEF2 and NFATC2 **(P, Q)** C-epitope increases membrane permeability (higher LDHA release in media). **(R)** Summary of findings: CH (aging or pressure-overload) elevates extracellular Gal-3 C-epitope oligomers, which cluster on cardiomyocyte membranes and activate maladaptive mechanotransduction (RhoA–ROCK1, stress fibers), leading to caveolae loss, nuclear damage, and cell death. AR/GA treatment blocks excess Gal-3 secretion and C-epitope oligomerization. This reduces membrane tension and mechanotransduction signaling, resulting in lower stress fibers, hypertrophy gene activation, cardiomyocyte enlargement, DNA damage, and cell death, thereby improving cardiac function. Levels of Gal-3 C-epitope and ANP decline with treatment and form a biomarker panel for therapeutic response.AR and GA emerge as effective inhibitors of Gal-3 C-epitope pathology in cardiac hypertrophy. Data are mean ± SD (3 experiments, ≥30–50 cells/experiment); t-test: *p ≤ 0.05, **p ≤ 0.01, ***p ≤ 0.001.

**Figure 10.**
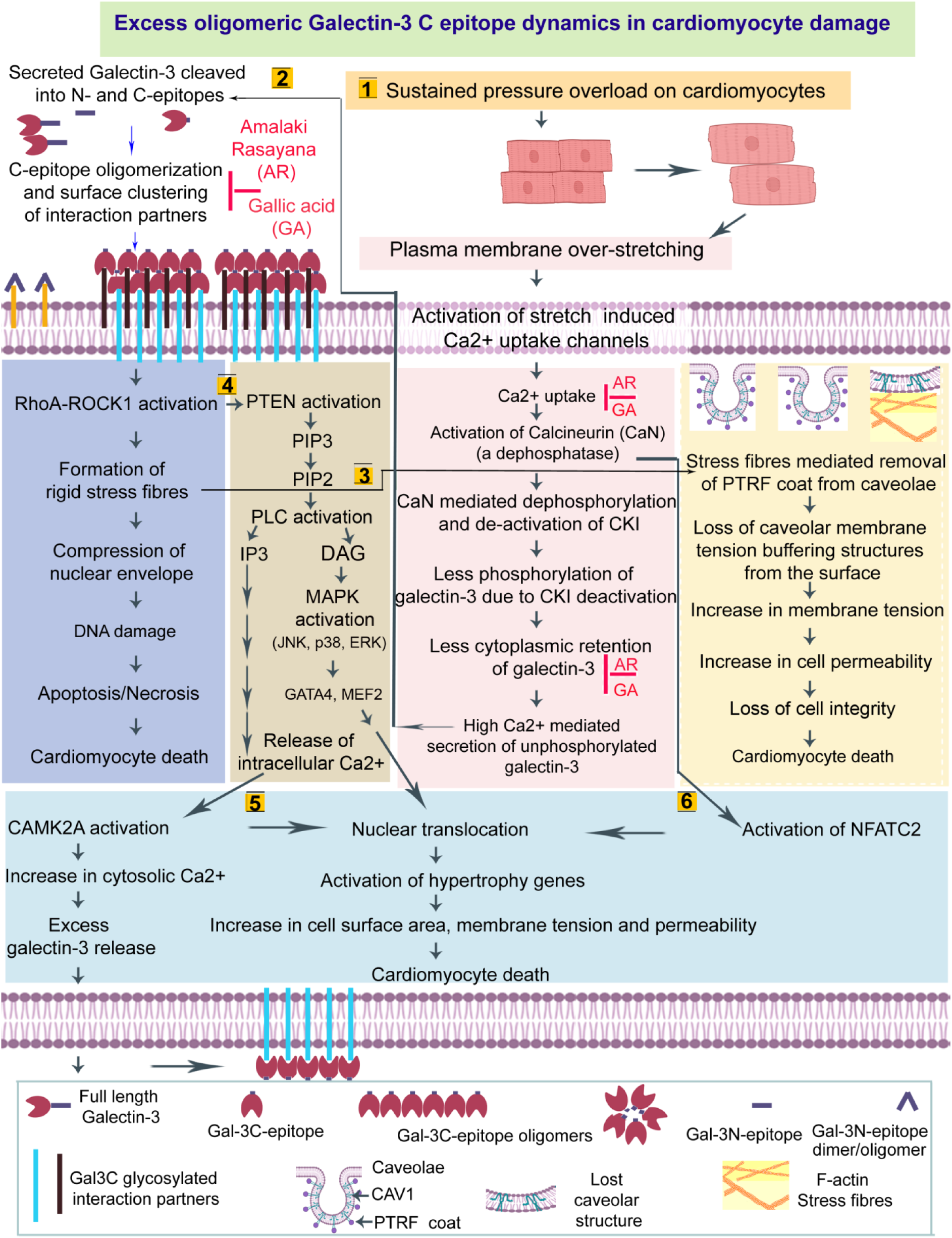
Proposed mechanisms of Gal-3 C-epitope oligomers–mediated CH and role of AR/GA therapy. Schematic model illustrating how extracellular Gal-3 C-epitope oligomerization drives cardiomyocyte hypertrophy and remodelling, and how AR/GA intervention mitigates these effects, integrating our findings with literature (citations: 43-50) 1. Pressure-overload stretches cardiomyocytes, raising intracellular Ca²⁺ that activates serine threonine dephosphatase-calcineurin. Calcineurin inhibits CKI (casein kinase I), reducing Gal-3 N-domain (Ser6) phosphorylation and increasing secretion of unphosphorylated Gal-3. 2. Extracellular Gal-3 is cleaved into N- and C-domains that homo-oligomerizes and bind on the cell surface; C-epitope oligomers form large clusters, causing membrane stress. This triggers RhoA–ROCK1 signaling, forming rigid F-actin fibers that compress the nucleus and damage chromatin. 3. Excess stress fibers dismantle caveolae (CAV-PTRF interaction loss), raising membrane tension and causing cell permeability and death. 4. RhoA–ROCK1 activates PTEN→PIP2 and PLC→DAG/MAPK, promoting hypertrophy gene transcription (MEF2, GATA4) and IP3-mediated Ca²⁺ release, further driving Gal-3 secretion. 5. High Ca²⁺ activates CAMK2A and NFATC2 pathways, driving hypertrophy gene expression and cell death. **Therapy:** AR/GA binds Gal-3’s CRD, inducing a conformation that promotes N-domain phosphorylation by CKI, increasing intracellular Gal-3 and reducing secretion. AR/GA also block Gal-3 surface clustering, preventing the RhoA–ROCK1–PTEN–PLC–Ca²⁺ cascade and its downstream damage. GA also reduces the deleterious levels of free calcium, deactivating calcineurin and PP1 dephosphatases induces Gal-3 dephosphorylation.

## 4. DISCUSSION

This study identifies extracellular galectin-3 C-terminal epitope (Gal-3 C-epitope) oligomers as key pathogenic drivers and drug-responsive biomarkers of cardiac hypertrophy (CH), with important therapeutic implications for the prevention of heart failure. We demonstrate that during pressure-overload induced left ventricular hypertrophy (PO-CH), galectin-3 undergoes enhanced secretion and proteolytic processing into distinct N- and C-terminal epitopes. Notably, Gal-3 C-epitope oligomers accumulate on cardiomyocyte surfaces, in the extracellular milieu, and in circulation, where their abundance correlates with adverse cardiomyocyte remodelling, cadiomyocyte loss and elevated cardiac stress markers, including atrial natriuretic peptide (ANP).

Although Amalaki rasayana (AR) is a multi-component phytomedicine, our findings reveal a clear and convergent mechanism by which AR and its major bioactive constituent, gallic acid (GA), modulate pathological Gal-3 biology. AR/GA enhanced galectin-3 phosphorylation, particularly at the Ser6 residue, promoting intracellular retention and limiting pathological secretion and extracellular oligomerization of the Gal-3 C-epitope. This, in turn, reduced Gal-3 C-epitope binding to cardiomyocyte surfaces, attenuated maladaptive hypertrophic signalling, prevented stress fibre formation, and preserved cardiomyocyte viability. Importantly, AR and GA improved cardiac structure and function in experimental models without detectable cytotoxicity, underscoring their cardioprotective potential.

Analysis of human cardiac tissues and sera further supports the translational relevance of these findings. In patients with cardiac hypertrophy, circulating Gal-3 C-epitope oligomer levels were altered relative to age-matched controls, despite elevated total Gal-3 concentrations. This divergence highlights a critical limitation of conventional total Gal-3 measurements and underscores the value of epitope-specific detection in distinguishing pathogenic from non-pathogenic Gal-3 pools. These data suggest that Gal-3 C-epitope oligomers may offer superior sensitivity for monitoring disease status and therapeutic response.

The dual role of Gal-3 C-epitope oligomers as both therapeutic targets and circulating biomarkers is particularly noteworthy. Changes in Gal-3 C-epitope oligomer levels closely paralleled regression of cardiac hypertrophy, positioning them as a promising tool for assessing treatment efficacy. Moreover, the capacity of AR/GA to suppress Gal-3 C-epitope oligomer formation supports their potential use as adjunctive therapies, complementing established neurohormonal interventions such as β-blockers and angiotensin-converting enzyme inhibitors [49]. The favourable safety profiles, dietary prevalence, and bioavailability of GA—delivered through AR—further facilitate translational potential for managing CH, left ventricular hypertrophy, and early-stage heart failure [50, 51, 52, 53]. While optimization of GA derivatives with enhanced Gal-3 C-epitope binding may be pursued, AR remains a clinically established formulation with demonstrated tissue bioavailability and multi-target engagement [22].

A key strength of this study lies in its integrated experimental design, combining mechanistic in vitro analyses, in vivo models of ageing- and pressure-overload induced hypertrophy, and validation using human samples. Together, these approaches establish a causal role for extracellular Gal-3 C-epitope oligomers in maladaptive hypertrophic mechanotransduction and identify AR as a rapidly translatable intervention. Limitations include the modest size of the human serum cohort and the absence of drug-naïve CH samples. While human specimens were analysed as independent biological validation rather than powered clinical endpoints, larger and longitudinal studies will be necessary to confirm the prognostic utility of Gal-3 C-epitope oligomers and to determine whether their reduction predicts improved clinical outcomes. Additionally, although a Ser6 phosphorylation-specific Gal-3 antibody is currently unavailable, complementary mutagenesis studies support the mechanistic conclusions. While GA emerged as the dominant active component, other constituents of AR—such as ellagic acid and vitamin C—may contribute synergistically to cardioprotection, warranting further biophysical and biochemical investigation.

Beyond cardiovascular disease, these findings open new avenues for targeting Gal-3 C-epitope oligomers in a broader spectrum of Gal-3–associated pathologies, including malignancies, non-alcoholic steatohepatitis, chronic kidney disease, and COVID-19 [54,55]. Future studies should assess the long-term effects of AR/GA therapy, explore synergy with standard-of-care treatments, and examine impacts on inflammatory and fibrotic remodelling.

In summary, this study demonstrates that Gal-3 C-epitope oligomers are central mediators of cardiac hypertrophy induced cardiomyocyte loss, and represent actionable therapeutic target and precision biomarker along with ANP. By selectively targeting the pathogenic Gal-3 C-epitope pool—rather than the total galectin-3 repertoire—Amalaki rasayana exemplifies a precision phytomedicine strategy with potential applicability across cardiovascular and systemic diseases.

## DECLARATIONS

### Supplementary Materials

Supplementary File 1 accompanies the main manuscript and includes supplementary figures, all uncropped, original Western and dot blot images used to support the study’s conclusions, as well as detailed information on the sources of the reagents and kits used.

### CRediT author statement

**Puja Laxmanrao Shinde:** Methodology, Formal analysis, Investigation, Visualization. **Vikas Kumar:** Methodology, Formal analysis. **Siddhartha Singh:** Methodology, Formal analysis, Investigation, Visualization.

**K.C. Sivakumar :** Methodology, Formal analysis, Investigation. **Abhirami P**: Methodology, Formal analysis. **Amit Mishra:** Methodology, Suggestions, Writing – review & editing **Rashmi Mishra:** Conceptualization, Methodology, Formal analysis, Resources, Investigation, Visualization, Supervision, Project administration, Funding acquisition, Writing – review & editing. All authors have read and approved the final manuscript.

## Funding

The work is supported by Indian Council of Medical Research (ICMR) extramural research grant (File no. 2020-0169/CMB/ADHOC-BMS) to Dr. Rashmi Mishra.

### Institutional review board statement/Compliance with Ethics Requirements

**Animal Ethics statement:** All animal experiments were carried out with the approval of the Institutional animal ethics committee (IAEC) in Rajiv Gandhi Centre for Biotechnology (RGCB) under the protocol no. IAEC/150/CCK/2012. Animal experiments were conducted by strictly following the rules and regulations of the Committee for the Purpose of Control and Supervision of Experiments on Animals (CPCSEA), Government of India. The animal studies strictly follow all ARRIVE guidelines.

**Human Ethics Statement:** Human samples were sourced from Innovative Research ((https://www.innov-research.com/, USA) following approval from the Institutional Human Ethics Committee of RGCB (File no. IHEC/12/2023_Ex/07, approval date: 20/03/2024) and in accordance with ICMR guidelines, Government of India. The Institutional Ethics Committee has granted a waiver of consent, as the samples were not collected by the investigator and do not contain any subject identifiers. In other words, the subjects were not in contact with the investigator.

### Informed Consent Statement

Not applicable

### Data availability statement

All data sets generated and analyzed to support the conclusions of this study are included in the main manuscript and its supplementary files, including un-cropped, original western and dot blot images.

## Acknowledgments

PLS and SS acknowledge research fellowships from the Department of Biotechnology (DBT) and University Grant Commission (UGC), respectively.

## Conflicts of Interest

The authors have declared no conflict of interest

## Declaration of generative AI use

During the preparation of this work the author(s) used ChatGPT and Grammarly in order to improve language and readability. After using this tool/service, the author(s) reviewed and edited the content as needed and take(s) full responsibility for the content of the published article.

## ABBREVIATIONS

ACTN2: Alpha-Actinin-2 protein
ALB: Albumin
ANP: Atrial Natriuretic Peptide
ANXA2: Annexin A2
Asn: Asparagine
Arg: Arginine
AR: Amalaki Rasayana
αSMA: Alpha smooth muscle actin
BA: Biological aged rats
BAPTA: 1,2(Bis-2-amino-5-methylphenoxy)ethane-N,N,N’,N’-tetraacetic acid tetrakis-acetoxymethyl) ester)
CAMK2A: Calcium/Calmodulin-dependent protein kinase II
CAMK2D: Calcium/Calmodulin dependent protein kinase II delta
CaN: Calcineurin
CAV1: Caveolin-1
CD44: Cluster of Differentiation 44
CD68: Cluster of Differentiation 68
CH: Cardiac hypertrophy
CRD: Carbohydrate Recognition Domain
cTNT: Cardiac troponin T
D-HCM: Dilated Hypertrophic cardiomyopathy
EGFP: Enhanced Green Fluorescent Protein
EGFR: Epidermal Growth Factor Receptor
ELISA: Enzyme-linked immunosorbent assay
FDA: Food and Drug Administration
GA: Gallic Acid
GAL-1: Galectin-1
Gal3: Galectin-3
Gal-3N: Cleaved fragment of Galectin-3 N-terminal (AA1 to AA113)
Gal-3C: Cleaved fragment of Galectin-3 C-terminal (AA114 to AA250)
GAL3BP: Galectin-3-Binding Protein
GAL-9: Galectin-9
GATA4: GATA Binding Protein 4
HCM: Hypertrophic cardiomyopathy
HEK293: Human Embryonic Kidney 293 cells,
HMGB1: High Mobility Group Box 1
IP: Immunoprecipitation
LDHA: Lactate dehydrogenase A
LVEF: Left Ventricular Ejection Fraction
LVFS: Left Ventricular Fractional Shortening
LVH: Left Ventricular Hypertrophy
MYH7: beta-Myosin Heavy Chain protein
MEF2A: Myocyte-specific Enhancer Factor 2A
mut R186S: Mutant EGFP tagged Galectin 3 with mutation of carbohdrate recognition binding domain (CRD)
NASH: Non alcoholic Steatohepatitis
NFATC2: Nuclear Factor of Activated T-cells-cytoplasmic-calcineurin-dependent 2
OBSCN: Obscurin
PLA: Proximity Ligation Assay (Duolink)
PO-CH: Pressure Overloaded rats with left ventricular cardiac hypertrophy
PPIA: Peptidylprolyl isomerase A (Cyclophilin A
PPID: Peptidylprolyl isomerase D
RhoA: Ras homolog family member A
ROCK1: Rho-associated coiled-coil-containing protein kinase 1
TMOD1: Tropomodulin-1
WGA: Wheat Germ Agglutinin
WT: Wild Type

## Notes

### Competing Interest Statement

The authors have declared no competing interest.

