## Supplementary Information File for "Targeting Galectin-3 C-epitope oligomers associated maladaptive mechanotransductive signaling in pressure-overload induced left ventricular cardiac hypertrophy"

#### Supplementary file 1

### Targeting Galectin-3 C-epitope oligomers associated maladaptive mechanotransductive signaling in pressure-overload induced left ventricular cardiac hypertrophy

***Authors:***

Puja Laxmanrao Shinde[a,c], Vikas Kumar [b,c,h,f], Siddhartha Singh[a, d,h], K.C.Sivakumar[e] , Abhirami P[a, d], Amit Mishra[g,i] and Rashmi Mishra,[a,c,d,i]*

This file has the following three sections:

1) Section A: Supplementary Figures and Tables
2) Section B: Supplementary Methods and Materials, Details and Sources of purchased reagents
3) Section C: Uncropped Dot and Western blot Images

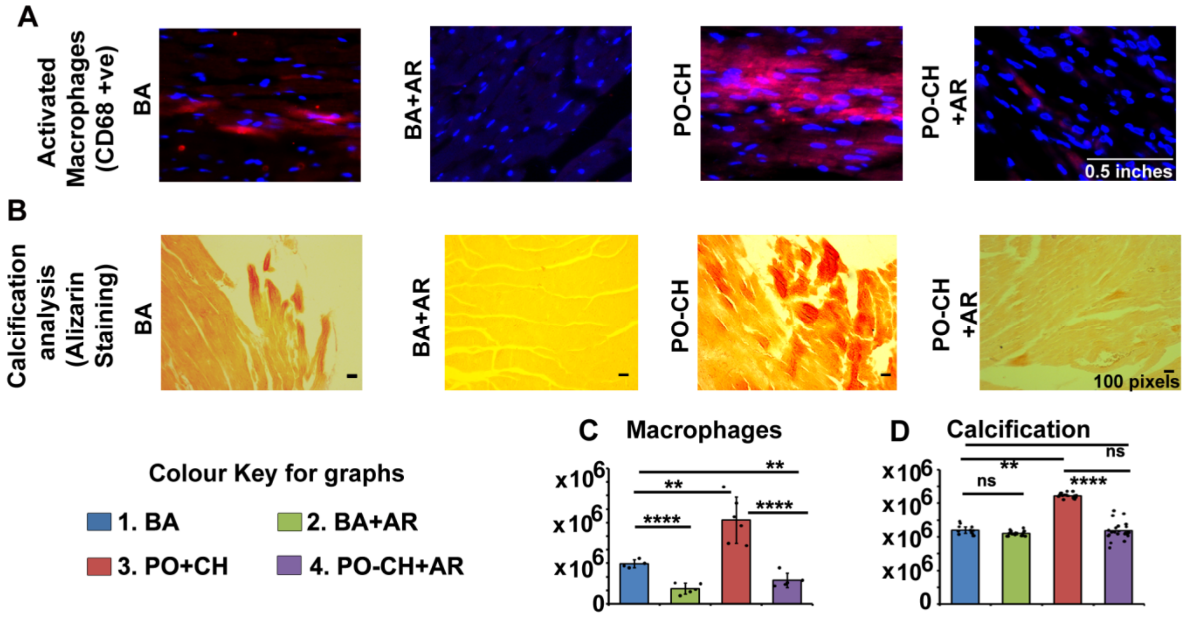

**Fig.S1. AR therapy in left ventricular hypertrophy reduces activated macrophage count and calcification: (A-C)** IHC for CD68+ Macrophages: AR-treated animals exhibit a significant reduction in activated macrophages, characterized by elongated, swollen phenotypic changes, compared to controls.

**(B-D)** Alizarin Red Staining: AR-treated hearts show a marked decrease in calcification, as evidenced by reduced calcium deposition.

Data are presented as mean ± SD (n = 4-8 rats/group). Statistical significance was determined by t-test: *$p < 0.05$, **$p < 0.01$, ***$p < 0.001$, ****$p < 0.0001$.

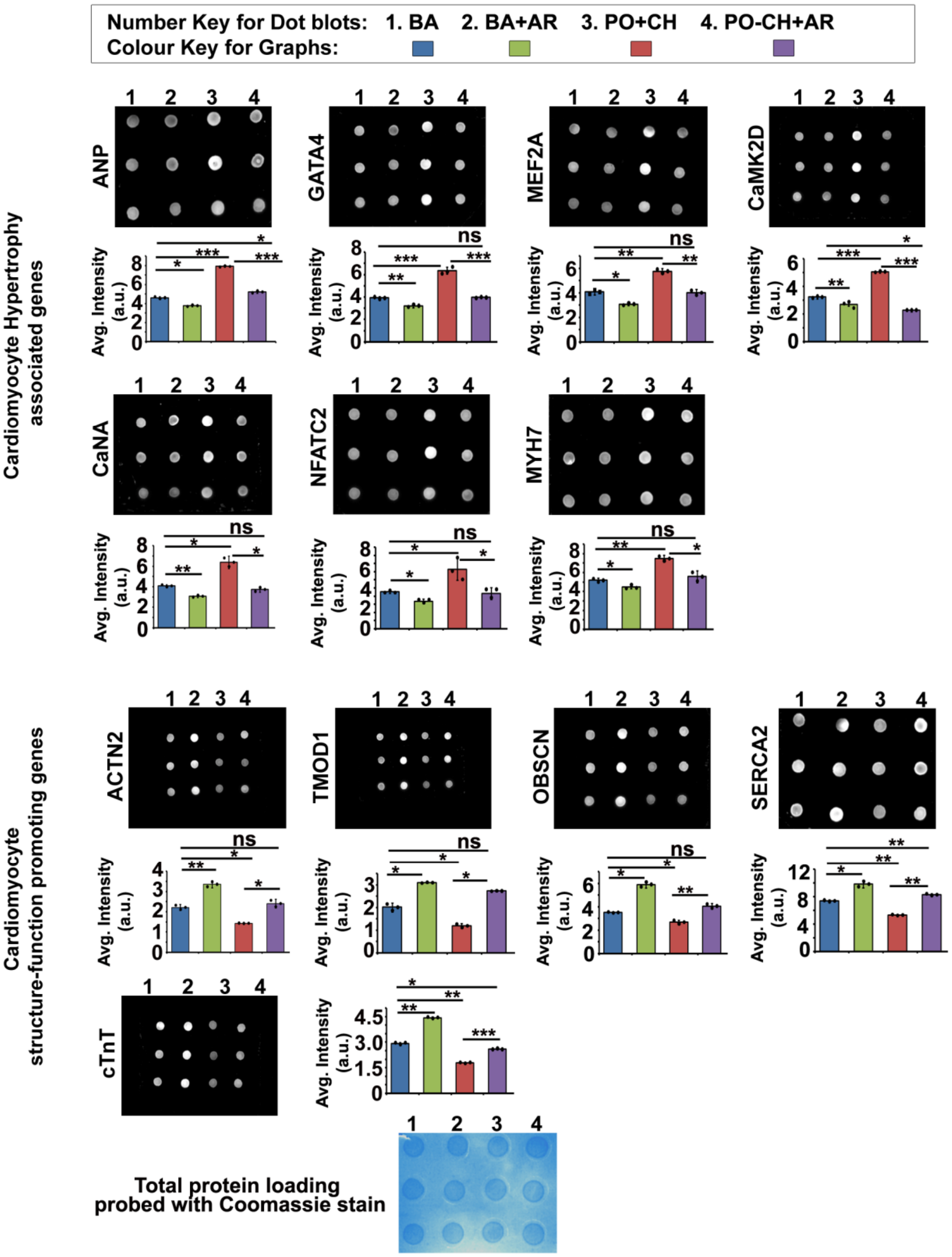

**Fig. S2. AR therapy in left ventricular hypertrophy reprograms cardiomyocyte-associated gene expression.**

AR-treated rats exhibit reduced expression of hypertrophy-associated proteins (ANP, GATA4, MYH7, MEF2A, CAMK2D, CaNA, NFATC2) and increased expression of adult cardiomyocyte structural and functional proteins (cTNT, ACTN2, TMOD1, OBSCN, SERCA2A), compared with untreated animals. KO-validated antibodies were used for signal detection. Data are presented as mean ± SD (n = 4–8 rats/group). Statistical significance was assessed by t-test: *$p < 0.05$, **$p < 0.01$, ***$p < 0.001$, ****$p < 0.0001$.

| TABLE S1 : Functions of genes probed in Fig. S1 | | |
|---|---|---|
| Gene Name | Function | References |
| **ANP** | • Atrial natriuretic peptide (ANP) is normally secreted at a basal level from atrial myocytes in the adult heart under physiological conditions. During prenatal development, ANP is produced by both atrial and ventricular myocytes, but ventricular expression declines after birth as the heart matures. In mature hearts, ANP expression becomes restricted mainly to the atria.<br>• Under pathological stress, such as ventricular stretch or pressure overload, ventricular myocytes re-activate ANP synthesis. Mechanical stretch of ventricular myocytes is a primary trigger for increased ANP production. Hormones, growth factors, and inflammatory cytokines further stimulate the production and release of ANP during cardiac hypertrophy.<br>• GATA4, a cardiac-enriched transcription factor, regulates ANP gene expression during Heart morphogenesis, prenatal cardiac development, and Maintenance of cardiac phenotype in adulthood.The activation of GATA4 during cardiac stress contributes to increased ANP expression, thereby linking it to the regulation of genes involved in hypertrophy.<br>• ANP acts as both a biomarker and a modulator of cardiac hypertrophy. It re-expresses as part of the fetal gene program, reflecting cardiac stress and compensatory remodelling. | • Kessler-Icekson G, Barhum Y, Schaper J, Schaper W, Kaganovsky E, Brand T. ANP expression in the hypertensive heart. *Exp Clin Cardiol*. 2002;7(2-3):80-84.<br><br>• Nandi SS, Mishra PK. Harnessing fetal and adult genetic reprograming for therapy of heart disease. *J Nat Sci*. 2015;1(4):e71.<br><br>• Bisping E, Ikeda S, Kong SW, et al. Gata4 is required for maintenance of postnatal cardiac function and protection from pressure overload-induced heart failure. *Proc Natl Acad Sci U S A*. 2006;103(39):14471-14476. doi:10.1073/pnas.0602543103 |
| **GATA4** | GATA4 is a zinc finger-containing transcription factor. Several stimuli that induce cardiac hypertrophy have been shown to enhance GATA4 translocational activity through phosphorylation, including pressure overload, phenylephrine, endothelin-1, and angiotensin II. These stimulations lead to phosphorylation of GATA4 at serine 105. GATA4 Phosphorylation occurs via ERK1/2 (Extracellular signal-regulated kinase 1/2) and p38 MAPK (Mitogen-activated protein kinase), resulting in enhanced DNA binding and transcriptional potency. Phosphorylated GATA4 binds to the promoters of genes such as ANP, BNP, and β-MHC, increasing their expression and facilitating adaptive | • Oka T, Maillet M, Watt AJ, et al. Cardiac-specific deletion of Gata4 reveals its requirement for hypertrophy, compensation, and myocyte viability. *Circ Res*. 2006;98(6):837-845. doi:10.1161/01.RES.0000215985.18538.c4 |

| | | |
|---|---|---|
| | remodelling, and consequently the hypertrophic growth. | |
| **Calcineurin A (CnA; calcium-activated phosphatase)** | • CnA is a (serine/threonine) $Ca^{2+}$/calmodulin-dependent phosphatase.<br>• Main target: NFAT transcription factors (NFATC2, NFATC3).<br>• Hypertrophic stress increases intracellular $Ca^{2+}$ and $Ca^{2+}$/calmodulin binding unlocks its phosphatase activity.<br>• CaNdephosporylates NFATC2 leading to its activation, translocation into the nucleus, and increase transcription of fetal cardiac markers (pro-hypertrophy markers -ANP, BNP, MYH7 etc).<br>• Studies showed that CnA activation is sufficient to induce cardiomyocyte hypertrophy. | • Molkentin JD, Lu JR, Antos CL, et al. A calcineurin-dependent transcriptional pathway for cardiac hypertrophy. *Cell.* 1998;93(2):215-228. doi:10.1016/s0092-8674(00)81573-1<br><br>• Molkentin JD. Calcineurin-NFAT signaling regulates the cardiac hypertrophic response in coordination with the MAPKs. *Cardiovasc Res.* 2004;63(3):467-475. doi:10.1016/j.cardiores.2004.01.021<br><br>• Molkentin JD. Calcineurin and beyond: cardiac hypertrophic signaling. *Circ Res.* 2000;87(9):731-738. doi:10.1161/01.res.87.9.731 |
| **NFATC2 (Nuclear Factor of Activated T cells, cytoplasmic 2)** | • NFATC2 is a transcription factor that plays a critical role in the cardiac hypertrophic remodelling through Calcineurin–NFAT signalling pathway.<br>• Pressure overload leads to mechanical stress on cardiomyocytes which activates calcium influx leading to stimulation of Calcineurin, a calcium/calmodulin-dependent phosphatase. Calcineurin dephosphorylates NFATC2, enabling its translocation from the cytoplasm to the nucleus. Inside the nucleus NFATC2 binds to hypertrophic gene promoters, often in cooperation with some other transcription partners such as GATA4, MEF2 to upregulate fetal cardiac genes like ANP (NPPA) and BNP (NPPB), MYH7. Upregulation of genes like ANP, BNP, further leads to hypertrophic remodelling. | • Bourajjaj M, Armand AS, da Costa Martins PA, et al. NFATc2 is a necessary mediator of calcineurin-dependent cardiac hypertrophy and heart failure. *J Biol Chem.* 2008;283(32):22295-22303. doi:10.1074/jbc.M801296200 |
| **CaMK2D** | • CaMK2D is one of the most important | • Anderson ME, Brown JH, |

| | | |
|---|---|---|
| **(CaMKIIδ) (Calcium/Calmodulin-dependent Protein Kinase IIδ)** | CaMKII isoforms in the heart, predominantly expressed in cardiomyocytes.<br>• Sustained increase in intracellular $Ca^{2+}$ (due to pressure overload, neurohormonal activation) activates CaMK2D.<br>• Activated CaMKIIδ enters the nucleus and phosphorylates HDAC4 (Histone Deacetylase 4).<br>• Phosphorylated HDAC4 gets exported out of the nucleus leaving MEF2 free and active (HDAC4 normally suppresses MEF2 by binding to it).<br>• OthersideCaMKIIδ also phosphorylatets RyR2 making it leaky for $Ca^{2+}$ which further leads to activation of pathways involved in progression of pathological cardiac hypertrophy (CaMK2 → MEF, Calcineurin → NFAT → ANP,BNP).<br>• Also CaMKIIδ → activates NFκB → inflammation → hypertrophy → fibrosis → heart failure progression | Bers DM. CaMKII in myocardial hypertrophy and heart failure. *J Mol Cell Cardiol*. 2011;51(4):468-473. doi:10.1016/j.yjmcc.2011.01.012<br><br>• Backs J, Song K, Bezprozvannaya S, Chang S, Olson EN. CaM kinase II selectively signals to histone deacetylase 4 during cardiomyocyte hypertrophy. *J Clin Invest*. 2006;116(7):1853-1864. doi:10.1172/JCI27438 |
| **MEF2A (myocyte enhancer factor-2A)** | • It is a transcription factor belonging to the MEF2 family.<br>• In heart, MEF2A controls growth, differentiation, and stress responses.<br>• It activates during cardiac hypertrophy through CaMKII and MAPK signalling and drives expression of fetal and structural genes, contributing to maladaptive hypertrophic growth. | • Passier R, Zeng H, Frey N, et al. CaM kinase signaling induces cardiac hypertrophy and activates the MEF2 transcription factor in vivo. *J Clin Invest*. 2000;105(10):1395-1406. doi:10.1172/JCI8551 |
| **MYH7 (β-myosin heavy chain)** | • **MYH7 (β-MHC)** is a key fetal gene reactivated during pathological cardiac hypertrophy. Its upregulation reduces contraction but conserves energy, reflecting an adaptive-to-maladaptive transition in stressed myocardium.<br>• Pressure overload → Mechanical stress → integrin/Fak → RhoA → activation of MEF2/GATA4 → ↑ MYH7.<br>• Calcineurin–NFATc pathway → NFATc translocate to nucleus → bind MYH7 promoter → ↑ transcription<br>• CaMKII → MEF2 activation → MEF2A activate fetal genes including MYH7.<br>• MAPK pathway (ERK1/2, p38) → Activates GATA4 and AP-1 → increase MYH7 expression.<br>• Thyroid hormone (T3) → T3 suppresses | • Krenz M, Robbins J. Impact of beta-myosin heavy chain expression on cardiac function during stress. *J Am Coll Cardiol*. 2004;44(12):2390-2397. doi:10.1016/j.jacc.2004.09.044<br><br>• Nadruz W Jr, Corat MA, Marin TM, Guimarães Pereira GA, Franchini KG. Focal adhesion kinase mediates MEF2 and c-Jun activation by stretch: role in the activation of the cardiac hypertrophic genetic program. *Cardiovasc Res*. |

| | | |
|---|---|---|
| | MYH7 → Low T3 (hypothyroidism, HF) → increased MYH7. | 2005;68(1):87-97. doi:10.1016/j.cardiores.2005.05.011<br>• Janssen R, Zuidwijk MJ, Kuster DW, Muller A, Simonides WS. Thyroid Hormone-Regulated Cardiac microRNAs are Predicted to Suppress Pathological Hypertrophic Signaling. *Front Endocrinol (Lausanne).* 2014;5:171. Published 2014 Oct 20. doi:10.3389/fendo.2014.00171 |
| **ACTN2 (α-actinin-2)** | • ACTN2 is a major Z-disc actin-crosslinking protein in cardiomyocytes.<br>• It stabilizes sarcomeres, anchors actin filaments, and acts as a mechanosensor. | • Ribeiro Ede A Jr, Pinotsis N, Ghisleni A, et al. The structure and regulation of human muscle α-actinin. *Cell.* 2014;159(6):1447-1460. doi:10.1016/j.cell.2014.10.056 |
| **TMOD1 (tropomodulin-1)** | • TMOD1 is an actin filament pointed-end capping protein.<br>• It regulates thin filament length in cardiomyocytes.<br>• In normal hearts, localized at the pointed end of actin filaments near the M-line and Prevents actin depolymerisation. Works with tropomyosin to stabilize the filament and keeps sarcomere function intact.<br>• TMOD downregulated or mislocalized during pathological conditions.<br>• Hypertrophic growth requires addition of new sarcomeres so reduced TMOD1 capping allows more actin polymerization → filament elongation, → sarcomere addition. This helps the cell grow in length/size (remodelling).<br>• Also in cardiac hypertrophy, TMOD1 levels mostly decrease. So reduced TMOD1 weakens thin filament stability, promoting pathological sarcomere remodelling leading to increase in mechanical stress signalling (Calcineurin–NFAT, MEF2). | • Gokhin DS, Fowler VM. Tropomodulin capping of actin filaments in striated muscle development and physiology. *J Biomed Biotechnol.* 2011;2011:103069. doi:10.1155/2011/103069<br>• Vasilescu C, Colpan M, Ojala TH, et al. Recessive TMOD1 mutation causes childhood cardiomyopathy. *Commun Biol.* 2024;7(1):7. Published 2024 Jan 2. doi:10.1038/s42003-023-05670-9 |
| **OBSCN** | • Obscurin is a giant sarcomeric protein, | • Young P, Ehler E, Gautel |

| | | |
|---|---|---|
| **(obscurin)** | localized at - M-line, Z-disc and Sarcoplasmic reticulum (SR) junctions. Acts as a scaffold, linking – Sarcomeres, Cytoskeleton, Sarcoplasmic reticulum membranes.<br>• OBSCN loss activates hypertrophic pathways indirectly.<br>• Loss of obscurin → unstable M-line and Z-disc → higher cytoskeletal tension → stimulation of pathways: Calcineurin–NFAT, RhoA–ROCK, MAPK (ERK1/2, JNK).<br>• SR disorganization (Misalignment of SR and T-tubules) → abnormal $Ca^{2+}$ handling (Slower $Ca^{2+}$ reuptake and irregular $Ca^{2+}$ sparks) → Activation of: CaMKII, NFAT, MEF2 → Hypertrophic genes expression. | M. Obscurin, a giant sarcomeric Rho guanine nucleotide exchange factor protein involved in sarcomere assembly. *J Cell Biol.* 2001;154(1):123-136. doi:10.1083/jcb.200102110 |
| **SERCA2 (ATP2A2 / SERCA2a)** | • SERCA2a is the cardiac isoform of the Sarcoplasmic Reticulum $Ca^{2+}$-ATPase encoded by the ATP2A2 gene.<br>• Pumps cytosolic $Ca^{2+}$ back into the SR during diastole.<br>• Downregulation of SERCA2a is a well-established hallmark of pathological hypertrophy and heart failure.<br>• Low level of SERCA2a → Slower $Ca^{2+}$ reuptake → Reduced SR $Ca^{2+}$ content → Weaker systolic $Ca^{2+}$ release → Diastolic $Ca^{2+}$ overload → Higher cytosolic $Ca^{2+}$ → activation of pathways → Calcineurin–NFAT, CaMKII–HDAC4–MEF2; → Hypertrophic gene remodelling. | • Lipskaia L, Chemaly ER, Hadri L, Lompre AM, Hajjar RJ. Sarcoplasmic reticulum Ca(2+) ATPase as a therapeutic target for heart failure. *Expert Opin Biol Ther.* 2010;10(1):29-41. doi:10.1517/14712590903321462 |
| **cTnT (cardiac troponin T; TNNT2)** | • cTnT (TNNT2) is a key component of the heart's contractile apparatus, which binds to tropomyosin to form a complex that regulates $Ca^{2+}$-dependent muscle contraction.<br>• In normal condition It regulates $Ca^{2+}$-dependent contraction, Controls interaction between actin and myosin and Essential for sarcomere contractility.<br>• In cardiac hypertrophy, cTnT functional alterations in 1)Phosphorylation (by PKC,PKA, CAMKII), 2)Isoform switching (fetal cTnT), 3)Proteolysis (due to cardiac stress related proteases: Calpain, Caspases) leads to reduce contractile efficiency and contribute to diastolic and systolic dysfunction. | • Sumandea MP, Pyle WG, Kobayashi T, de Tombe PP, Solaro RJ. Identification of a functionally critical protein kinase C phosphorylation residue of cardiac troponin T. *J Biol Chem.* 2003;278(37):35135-35144. doi:10.1074/jbc.M306325200<br><br>• Anderson PA, Greig A, Mark TM, et al. Molecular basis of human cardiac troponin T isoforms |

| | | |
|---|---|---|
| | | expressed in the developing, adult, and failing heart. *Circ Res.* 1995;76(4):681-686. doi:10.1161/01.res.76.4.681<br><br>• Ke L, Qi XY, Dijkhuis AJ, et al. Calpain mediates cardiac troponin degradation and contractile dysfunction in atrial fibrillation. *J Mol Cell Cardiol.* 2008;45(5):685-693. doi:10.1016/j.yjmcc.2008.08.012 |

| TABLE S2: Functions of gene probed in Figure 9 | | |
|---|---|---|
| Gene Name | Functions | References |
| **RhoA-ROCK1** | • RhoA (Ras homolog family member A) is a small GTPase that plays a central role in cytoskeletal dynamics, mechanotransduction, and gene expression.<br>• In pathological cardiac hypertrophy, it acts as a mechanosensitive signalling molecule. It is responsible for converting mechanical stress into hypertrophic gene expression.<br>• It promotes hypertrophy via ROCK, MRTF-SRF, YAP, PKC, and actin dynamics.<br>• RhoA–ROCK1 (Rho-associated kinase) pathway - ROCK1 phosphorylates MLCP → ↑ myosin light chain phosphorylation, which further increases contractility and actin remodelling.<br>• MRTF-A/B and SRF (Serum Response Factor) pathway - Actin polymerisation releases MRTF free → moves to nucleus and activates SRF (Serum Response Factor) → Expression of fetal cardiac genes.<br>• RhoA →Actin polymerisation → 1) | • Miyamoto S. Untangling the role of RhoA in the heart: protective effect and mechanism. *Cell Death Dis.* 2024;15(8):579. Published 2024 Aug 9. doi:10.1038/s41419-024-06928-8<br><br>• Kilian LS, Voran J, Frank D, Rangrez AY. RhoA: a dubious molecule in cardiac pathophysiology. *J Biomed Sci.* 2021;28(1):33. Published 2021 Apr 28. doi:10.1186/s12929-021-00730-w |

| | | |
|---|---|---|
| | Sequesters AMOT 2) LATS1/2 inhibition → YAP activation → Hypertrophy and fibrosis gene transcription.<br>• RhoA–PKC interaction → PKC activation enhances hypertrophic pathways → Supports sustained hypertrophy growth. | |
| **CAV1** | • Caveolin is an important structural caveolar protein, which performs various functions- 1) Regulation of endothelial function, 2) Regulation cardiac nitric oxide biogenesis, 3) Cellular lipid homeostasis, 4) Oxidative stress, 5) Modulation of signal transduction pathways which mediates oxidative stress and inflammatory responses. All these results in maladaptive cardiac remodelling and progression of cardiac hypertrophy.<br>• Cav-1 interacts with TGF-ß Receptor and blocks the downstream pathway leaving Smad2/3 unphosphorylated which leads to degradation of TGF-β responsive protein. In Cav-1 depleted condition activation of TGFß/SMAD2 → Induction of excessive extracellular matrix deposition → Cardiac fibrosis → Pathological hypertrophic growth.<br>• Cav-1 depletion → Increase in pumping of Na+/K+-ATPase → Inhibition of cardiotonic steroids (CTS)-mediated growth induction → excess collagen production → Cardiac fibrosis driven pathological hypertrophic growth. | • An Z, Tian J, Zhao X, et al. Regulation of cardiovascular and cardiac functions by caveolins. *FEBS J*. 2024;291(17):3753-3761. doi:10.1111/febs.16798<br><br>• Tian J, Popal MS, Huang R, et al. Caveolin as a Novel Potential Therapeutic Target in Cardiac and Vascular Diseases: A Mini Review. *Aging Dis*. 2020;11(2):378-389. Published 2020 Mar 9. doi:10.14336/AD.2019.09603 |
| **PTRF** | • Polymerase I and transcript release factor (PTRF) is an essential structural protein required for the assembly of caveolae, the small invaginations on the plasma membrane that regulate mechanotransduction, lipid homeostasis, and signalling. It is a critical cardioprotective protein and its absence causes caveolae-related cardiomyopathy. | • Taniguchi T, Maruyama N, Ogata T, et al. PTRF/Cavin-1 Deficiency Causes Cardiac Dysfunction Accompanied by Cardiomyocyte Hypertrophy and Cardiac Fibrosis. *PLoS One*. 2016;11(9):e0162513. Published 2016 Sep 9. doi:10.1371/journal.pone.0162513 |

| | | |
|---|---|---|
| | • Deficiency of PTRF in the heart leads to → Loss of caveolae →<br>o Abnormal membrane tension and dysregulated signalling,<br>o Increased activation of: p38 MAPK, ERK1/2, AKT, Fibrotic pathways,<br>o Promote hypertrophy, fibrosis, electrocardiographic abnormality and<br>o Upregulation of molecular markers of hypertrophy (ANP, BNP, β-MHC). | |
| **LaminA** | • Lamin A is an intermediate-filament proteins that form the nuclear lamina, a mesh-like structure below the inner nuclear membrane.<br>• Lamin A provides → Nuclear stiffness, nuclear structural integrity, regulates gene expression, mechanotransduction, and chromatin organisation, and Resistance to mechanical load during hypertrophic stress.<br>• During pathological hypertrophy, → mechanical stress increases, → Lamin A expression increases.<br>• Lamin A dysfunction → nuclear deformation, fragility, and DNA damage.<br>• On stiffened extracellular matrix (ECM), contractile cells maintain high lamin-A levels because lamin-A phosphorylation is low and its degradation by MMP-2 is slow; which stabilised (low-turnover) lamin-A—along with strong actomyosin contractility—protects the nucleus by reducing nuclear envelope rupture and minimising DNA damage. | • Donnaloja F, Carnevali F, Jacchetti E, Raimondi MT. Lamin A/C Mechanotransduction in Laminopathies. Cells. 2020;9(5):1306. Published 2020 May 24. doi:10.3390/cells9051306<br><br>• Kervella M, Muchir A. Role of Nuclear Lamins in the Regulation of the Genome: Focus on CardioLaminopathy. *Subcell Biochem*. 2025;115:1-22. doi:10.1007/978-3-032-00537-3_1<br><br>• Cho S, Vashisth M, Abbas A, et al. Mechanosensing by the Lamina Protects against Nuclear Rupture, DNA Damage, and Cell-Cycle Arrest. *Dev Cell*. 2019;49(6):920-935.e5. doi:10.1016/j.devcel.2019.04.020 |
| **γ-H2AX** | • p-γ-H2AX (Phosphorylated form of histone H2AX protein at ser139), which is considered a sensitive marker for the DNA double-strand breaks (DSBs).<br>• In the presence of hypertrophic stress (Pressure overload, stretch, angiotensin II, etc.), the level of | • Tossetta G, Fantone S, Compagnucci P, et al. γ-H2AX: A useful tool to detect DNA damage in sudden cardiac death heart tissues, an experimental study. *Tissue Cell*. 2025;96:103042. doi:10.1016/j.tice.2025.103042 |

| | | |
|---|---|---|
| | ROS increases, leading to mitochondrial dysfunction and eventually nuclear DNA damage. When DNA DSBs occur, kinases like ATM and ATR phosphorylate H2AX at ser139.<br>• Higher γ-H2AX levels are reported in failing hearts, which confirms the correlation between increased DNA damage and hypertrophic severity.<br>• ATM/ATR–γ-H2AX pathway also activates hypertrophic signalling. | • Dai Z, Ko T, Fujita K, et al. Myocardial DNA Damage Predicts Heart Failure Outcome in Various Underlying Diseases. *JACC Heart Fail*. 2024;12(4):648-661. doi:10.1016/j.jchf.2023.09.027<br><br>• Nakada Y, Nhi Nguyen NU, Xiao F, et al. DNA Damage Response Mediates Pressure Overload-Induced Cardiomyocyte Hypertrophy. *Circulation*. 2019;139(9):1237-1239. doi:10.1161/CIRCULATIONAHA.118.034822 |
| **YAP** | • Yes-associated-protein (YAP) is a transcriptional co-activator that regulates cell proliferation, differentiation, stress response, and mechanical signalling in cardiomyocytes.<br>• The Hippo pathway (MST1/2 → LATS1/2) inhibits YAP through phosphorylation. In the absence of Hippo signalling, YAP becomes dephosphorylated, enters the nucleus, and binds TEAD to activate hypertrophy-associated gene expression.<br>• YAP is activated by mechanical stress, ECM stiffness, and GPCR signalling. GPCR ligands (LPA, S1P, thrombin) activate YAP via Gα12/13–RhoA signalling.<br>• RhoA →Actin polymerisation → 1) Sequesters AMOT 2) LATS1/2 inhibition → YAP activation/ Nuclear translocation → Hypertrophy and fibrosis associated genes activation such as CTGF, CYR61.<br>• Phospholipase D-generated phosphatidic acid also contributes to RhoA-mediated LATS1/2 inhibition. | • Wang J, Liu S, Heallen T, Martin JF. The Hippo pathway in the heart: pivotal roles in development, disease, and regeneration. *Nat Rev Cardiol*. 2018;15(11):672-684. doi:10.1038/s41569-018-0063-3<br><br>• Han H, Qi R, Zhou JJ, et al. Regulation of the Hippo Pathway by Phosphatidic Acid-Mediated Lipid-Protein Interaction. *Mol Cell*. 2018;72(2):328-340.e8. doi:10.1016/j.molcel.2018.08.038<br><br>• Meng F, Xie B, Martin JF. Targeting the Hippo pathway in heart repair. *Cardiovasc Res*. 2022;118(11):2402-2414. doi:10.1093/cvr/cvab291 |

#### Section B: Supplementary Methods and Materials:

##### 2.1. *Reagents, Cell Lines, and Patient Samples*

Cell lines: Rat cardiomyoblasts-H9c2(2-1) and kidney cells-HEK-293, were procured from ATCC. All plasmid constructs were custom-synthesized by Zellebiotech.com. The pEGFP-hGal3 plasmid (#73080) was also obtained from Addgene (USA). Most reagents were purchased from Merck/Sigma-Aldrich. Galectin-3 ELISA kits—Rat ERLGALS3 and Human DGAL30—were sourced from Thermo Scientific and R&D Systems, respectively. Antibodies and dilutions used are as follows: CBP-112 AP, Rabbit anti-Galectin-3 (C-epitope): 1:1000; CBP-101 AP, Rabbit anti-Galectin-3 (N-epitope): 1:1000; Galectin-3 Mouse mAb (B2C10, sc-32790): 1:500; A13506, Galectin-3 KO validated ab 1:1000; NPPA Rabbit pAb (A1609): 1:5000; Albumin Rabbit pAb (A1363): 1:1000; Anti-Cyclophilin A (PPIA, AB58144): 1:5000. Anti-galectin-3 C- and N-epitope antibodies were validated for specificity with blocking peptides, P-CBP35.c and P-CBP35 (FabGennix International, USA) respectively, as per the manufacturer's protocol. Treatment-naive human heart tissue arrays were acquired from TissueArray.Com (USA). Human serum samples were procured from the Innovative research Inc. USA.

##### 2.2. *Standardised Preparation of Amalaki Rasayana by Arya Vaidya Sala, Kottakkal and its chemical fingerprinting, quantitation and quality control*

**A)** The preparation and other chemical characterizations of Amalaki rasayana (AR) has been described in detail in our previous publication [1,2]. Amalaki rasayana (AR) and the vehicle preparation consisting of ghee and honey (GH) were prepared by Arya Vaidya Sala, Kottakkal, Kerala, India, following the standard procedures prescribed in classical Ayurvedic texts [3,4]. Fresh green Amalaki fruits and their dried forms were procured from authenticated traders sourcing material from the Sathyamangalam region of Tamil Nadu (11.5048°N, 77.2384°E). The dried fruits were additionally procured from traders in Madhya Pradesh who source material from Chhattisgarh (21.2787°N, 81.8661°E) and Madhya Pradesh (22.9734°N, 78.6569°E), respectively.

Fresh gooseberries used for juice preparation were harvested during November and March, whereas fruits intended for powder preparation were collected in November. All experimental studies were performed using AR or GH prepared from a single production batch. The preparation protocol, described below, has also been reported previously [5].

**Preparation Procedure**

**Step 1:**

Dried gooseberry fruits (Phyllanthus emblica) were pulverized to 80-mesh particle size using a Tyco pulverizer.

**Step 2:**

Fresh gooseberry fruits were crushed using a juice extractor to obtain fresh juice.

**Step 3:**

The powdered material obtained in Step 1 and the fresh juice obtained in Step 2 were mixed in a 1:1 ratio and dried using a vacuum tray dryer for 24 h at 55 °C under 700 mmHg pressure. The dried mass was subsequently pulverized.

Steps 2 and 3 were repeated 20 times. According to Ayurvedic texts, a total of 21 cycles of this process—referred to as bhavana (trituration)—are required to enhance the intrinsic biological activity of the active ingredients present in the herb. This process involves repeated mixing of the powdered material with a liquid medium, drying, and re-pulverization. The procedure followed here conforms to the standard method recommended for the preparation of Amalaki rasayana [3–5].

**Step 4:**

The processed Amalaki powder obtained above was blended with honey (M/s Galaxy Honey, Kottakkal, Kerala) and ghee ("Milma," Malabar Milk Marketing Federation, Government of Kerala, Kozhikode, Kerala) in a ratio of 1:2:0.5 to obtain a thick, paste-like formulation designated as AR, as described previously [1,2,5]. The detailed composition of the ghee used is provided in Supplementary Table S8 of our publication, Kumar et al., Scientific Reports, 2017 [1].

The preparation method adheres to the classical Ayurvedic text Charaka Samhita. The addition of honey and ghee to the triturated Amalaki powder serves as a carrier or adjuvant (anupana), which improves acceptability, palatability, and bioavailability of the active constituents [4,5]. Ghee and honey mixed in a fixed proportion (2:0.5) were used consistently in the preparation. Both AR and GH formulations were subjected to comprehensive quality assurance and quality control testing.

**B)** ***Quality Assurance and Quality Control***

Quality assurance and quality control assays were conducted following standardized protocols described by (Dwivedi et al. (2012) [5]. Authentication of the Amalaki fruits and powder was performed by the Centre for Medicinal Plants Research, Kottakkal, and the

Quality Assurance Department of Arya Vaidya Sala, which is approved by the Government of India for testing and issuing quality control certificates for Ayurvedic raw materials and finished products (https://ayush.gov.in/#!/quality_standard).

The Ayurvedic Pharmacopoeia of India (API) confirmed that the macroscopic characteristics (shape and taste of fruits) and microscopic features (structure of pericarp, mesocarp, and vascular bundles) of Amalaki were consistent with the official pharmacopoeial standards (http://www.ayurveda.hu/api/API-Vol-1.pdf) [6]. The API provides the authoritative standards to be adopted for Ayurvedic preparations.

High-performance thin-layer chromatography (HPTLC) profiling of raw Amalaki fruits, Amalaki powder, samples obtained after the 1st, 10th, 15th, and 21st trituration cycles, and the final formulation demonstrated the presence of gallic acid (1.5%w/w; equal to 1.5 mg per 100mg of AR) and ellagic acid (0.40%w/w, equal to 0.4 mg per 100 mg of AR); when compared with authenticated standards. These results are shown in Figure S1A of our publication Kumar et al., 2017, Scientific reports; https://doi.org/10.1007/s11010-019-03637-1 [1].

The current manuscript is a continuation of our earlier work that comprehensively described the preparation, chemical fingerprinting, and characterization of Amalaki rasayana (AR). In our previous studies (Scientific Reports, 2017; https://doi.org/10.1038/s41598-017-09225-x and Molecular and Cellular Biochemistry, 2020; https://doi.org/10.1007/s11010-019-03637-1) [1, 2], all experimental procedures were conducted in accordance with the guidelines laid down by the Department of AYUSH, Government of India, which incorporate international regulatory standards, similar to those for the international medicines preparation and marketing guidelines.

**C)** ***Characterization of Components of Amalaki Rasayana (AR)***

Reference: Kumar et al., Scientific Reports, 2017 [1]

**I. Solubility Analysis**

To determine the solubility of AR and the vehicle control ghee + honey (GH), the following solvent systems were evaluated:

(i) acetonitrile:water (20:80 v/v and 70:30 v/v),

(ii) methanol:water (20:80 v/v and 70:30 v/v),

(iii) ethanol:water (20:80 v/v and 70:30 v/v), and

(iv) diethyl ether:water (20:80 v/v and 70:30 v/v).

For each condition, 10 mg of AR or GH was added to the respective solvent, and solubility was assessed based on the maximum extent of dissolution.

As summarized in Supplementary Table S1A of our published work Kumar et al., Scientific Reports, 2017 [1], AR exhibited good solubility in acetonitrile and ethanol, moderate solubility in methanol and diethyl ether, whereas GH showed good solubility in diethyl ether and moderate solubility in the remaining solvents. The RP-HPLC profile of AR dissolved in ethanol differed markedly from that of GH in the same solvent (see Supplementary Fig. S1b, d; Supplementary Tables S1A and S1C in Kumar et al., Scientific Reports, 2017 [1]). Overall, AR demonstrated good solubility in acetonitrile and ethanol and moderate solubility in methanol. A solvent system of 20% ethanol in water (20:80 v/v) was identified as optimal for solubilizing both AR and GH.

**II. RP-HPLC Profiling of AR and GH**

Following solubility assessment, RP-HPLC profiling was performed for both AR and GH. Analyses were conducted using a Waters HPLC system equipped with a Symmetry C18 column (150 mm × 3.9 mm i.d., 5 µm), a 2707 autosampler, and a 2489 UV/Visible detector. Data acquisition and processing were performed using Breeze software (Waters, USA).

Chromatographic separation was achieved at a flow rate of 1 mL/min using a mobile phase consisting of Solvent A [water:acetic acid (99.9:0.1 v/v)] and Solvent B (acetonitrile), following the gradient program described in Supplementary Table S1B in Kumar et al., Scientific Reports, 2017 [1]. The column temperature was maintained at 25 °C, the detection wavelength was set at 265 nm, and an injection volume of 25 µL was used. The elution gradient enabled separation of compounds from relatively more hydrophobic to less hydrophobic fractions. For further identification of AR constituents, LC–MS analysis was subsequently performed.

The RP-HPLC profile of AR revealed distinct peaks at retention times of 0.897, 2.030, 4.900, 8.109, and 9.465 min (see Supplementary Fig. S1b, d; Supplementary Table S1C in Kumar et al., Scientific Reports, 2017 [1]). When AR was dissolved in acetonitrile, additional peaks were observed at retention times of 2.155, 2.568, 5.482, 7.985, 18.026, and 18.210 min, which differed from the GH profile under identical conditions, suggesting differential composition (see Supplementary Fig. S1c, e; Supplementary Table S1C in Kumar et al., Scientific Reports, 2017). These distinct chromatographic signatures further justified the use of LC–MS analysis for component identification.

**III. HPTLC and LC–MS Analysis of AR**

**Step 1:**

One gram of AR was dissolved in 20 mL of solvent (20% ethanol in water) in a conical flask and stirred overnight at 4 °C.

**Step 2:**

An aliquot of 10 µL from the resulting solution was applied onto a TLC plate (silica gel 60 F254). Chromatographic separation was performed using a mobile phase consisting of toluene:ethyl acetate:formic acid:methanol in a ratio of 6:6:1.6:0.4.

**Step 3:**

Following TLC separation, three distinct fractions, based on polarity, designated as C1, C2, and C3, were resolved on the silica plate.

**Step 4:**

The separated bands were visualized under UV illumination, scraped from the plate, dissolved in 100% methanol, and subsequently lyophilized.

**Step 5:**

The lyophilized fractions were reconstituted in acetonitrile and subjected to LC–MS analysis using a Synapt-G2 Q-TOF LC-MS/MS system (Waters). MassLynx software was used for data acquisition, and PLGS v2.5.3 was used for raw data processing.

**Step 6:**

The acquired LC–MS data were uploaded to the XCMS online platform for metabolite identification.

In addition, HPTLC profiling of the finished formulation confirmed the presence of gallic acid and ellagic acid (see Supplementary Fig. S1a; Supplementary Tables S2A and S2B in Kumar et al., Scientific Reports, 2017 [1]).

Since only two constituents were chemically anchored using authenticated standards through HPTLC and LC–MS analyses—namely, gallic acid and ellagic acid—the remaining HPLC peaks represent unassigned fingerprint peaks. To further characterize these components, LC–MS analysis was performed on the lyophilized AR fractions. This analysis revealed enrichment of several putative bioactive metabolites, including anti-inflammatory arachidonate derivatives (eicosatetraenoic acid), norepinephrine sulfate, and vitamin-related metabolites, as identified using the XCMS online metabolomics platform (see Supplementary Fig. S1f; Table 1 in Kumar et al., Scientific Reports, 2017 [1])). In untargeted metabolomics, the use of XCMS software for LC-MS analysis often filters out common or extremely abundant primary metabolites to highlight "enriched" or "unique" components. Hence, Table

1 of Kumar et al., Scientific Reports, 2017 [1], we specifically lists components that were less abundant and were not identified through RP-HPLC.

**IV. *UPLC Analysis and Chemical Formula Confirmation of Gallic Acid and Ellagic Acid***

Reference: Kumar et al., Molecular and Cellular Biochemistry, 2020 [2]

Dried AR samples were reconstituted in LC–MS grade acetonitrile and methanol (J.T. Baker). Chromatographic separation was performed using an Accela ultra-high-performance liquid chromatography (UHPLC) system (Thermo Fisher Scientific, USA) equipped with an Accucore C18 column (150 × 2.1 mm, 2.6 µm particle size). The column oven temperature was maintained at 40 °C, and the autosampler temperature was set at 4 °C. A 5 µL sample was injected using a gradient mobile phase system consisting of solvent A (0.1% formic acid in water) and solvent B (acetonitrile). A linear gradient was applied from 5% B at 0 min to 95% B at 15 min at a flow rate of 350 µL/min.

Mass spectrometric acquisition was performed on a Q-Exactive Orbitrap mass spectrometer (Thermo Fisher Scientific, MA, USA) coupled with a heated electrospray ionization (HESI) source operated in both positive and negative electrospray ionization (ESI) modes. The capillary temperature was maintained at 300 °C and the probe heater temperature at 320 °C, with sheath gas set at 45 arbitrary units. The auxiliary gas was set at 5 and 12 arbitrary units for ESI+ and ESI– modes, respectively. The tube lens voltage was set at 50 V, and the mass scan range was set from 70 to 1000 m/z. The Orbitrap mass analyzer was operated at a resolution of 70,000. Tandem mass spectrometry (MS/$MS^2$) spectra were acquired using collision energies between 30 and 40 eV. Data acquisition and processing were performed using Thermo Scientific Xcalibur software (version 3.0).

For metabolite processing and identification, raw instrument files were converted to mzXML format using ProteoWizard (http://proteowizard.sourceforge.net/) and subsequently processed using XCMS and CAMERA for peak detection, alignment, annotation, and metabolite identification, as described previously. Putative metabolites identified by XCMS were further validated using accurate mass measurements and retention time matching against authenticated standards. Experimental MS/$MS^2$ spectra were also compared with reference compound spectra for confirmation.

**V. *LC–MS analysis*** (see Fig. 2 in Kumar et al., Molecular and Cellular Biochemistry, 2020 [2]) confirmed the presence of gallic acid (GA) and ellagic acid (EA) in the AR extract. Panels (a) and (b) showed comparative mass spectra of the GA standard and AR extract, respectively, while panels (c) and (d) show comparative mass spectra of the EA standard and AR extract [2]).

Notably, our 2020 study employed a targeted metabolomics approach using high-resolution LC–MS to validate the presence of key biomarkers in the AR extract.

Gallic Acid Identification:

Gallic acid was confirmed in the AR extract by matching both its exact molecular mass and retention time with the authenticated standard.

- Retention time (RT): approximately 4.46 min.
- Mass spectral data (m/z):

o The GA standard in negative ion mode (ESI−) exhibited a dominant peak at 169.0134 m/z.

o The AR extract showed a corresponding peak at 169.0133 m/z, confirming molecular identity.

#### VI. *Selection and Quantification of Analytical Markers for batch to batch authentication and reproducibility*

HPTLC profiling of the finished AR formulation revealed a major enrichment of gallic acid (1.5% w/w; 1.5 mg per 100 mg of AR) and ellagic acid (0.40% w/w; 0.4 mg per 100 mg of AR) (see Supplementary Fig. S1a; Supplementary Tables S2A and S2B in Kumar et al., Scientific Reports, 2017 [1]). Further confirmation by ultra-high-performance liquid chromatography (UHPLC) of dried AR extracts validated the presence of both compounds through retention time and mass spectral matching with authenticated standards, yielding characteristic m/z values of 169.0133 m/z for gallic acid and ~301 m/z for ellagic acid.

Quantitative analysis demonstrated that gallic acid is substantially more enriched than ellagic acid in both the finished AR formulation and the dried powder. Gallic acid has also been reported as a dominant phytochemical constituent in AR, representing up to >42% relative abundance in some analytical reports [7]. Gallic acid (GA) is known to play central roles in regulating myocardial bioenergetics, redox homeostasis, and antioxidant defense mechanisms.

Our previous pharmacokinetic and long-term AR feeding studies further established biological exposure relevance of GA. A daily oral dose of 500 mg/kg body weight of AR corresponds approximately to an intake of 7.5 mg gallic acid and 2.0 mg ellagic acid.

LC–MS analysis of serum samples from animals long-term fed AR for 12 months demonstrated sustained circulating levels of gallic acid, supporting its systemic bioavailability and translational relevance.

Multiple independent studies have demonstrated the cardioprotective efficacy of gallic acid in AR [8-9]. **Accordingly, gallic acid was selected as the primary active biomarker for mechanistic investigations assessing suppression of the deleterious effects of Gal-3 C-epitope in in vitro models. In addition, quantitative estimation of gallic acid provides a robust quality control metric for batch-to-batch consistency and therapeutic preservation in marketed AR formulations.**

Importantly, gallic acid was consistently confirmed across multiple orthogonal analytical platforms:

- HPTLC confirmed its presence and enrichment in the finished formulation;
- High-resolution LC–MS confirmed molecular identity through exact mass and retention time matching;
- Targeted metabolomics independently validated gallic acid as a dominant marker metabolite.

Based on this convergent analytical validation, gallic acid was selected as the analytical marker for standardization in accordance with marker-based quality control principles recommended for complex botanical formulations.

##### VII. *Quantitative Marker Selection and Scientific Rationale*

Marker compounds were selected based on the following criteria:

1. High abundance and reproducibility in the finished formulation and across batches.
2. Chemical stability under formulation and storage conditions.
3. Biological relevance, particularly involvement in cardioprotective, antioxidant, and anti-inflammatory pathways.
4. Detectability and confirmability across orthogonal analytical platforms (HPTLC, UHPLC, HR-LCMS, targeted metabolomics).

The selected markers were quantitatively determined. HPTLC profiling of the finished formulation demonstrated a major enrichment of gallic acid (1.5% w/w; 1.5 mg per 100 mg of AR) and ellagic acid (0.40% w/w; 0.4 mg per 100 mg of AR) see Supplementary Fig. S1a; Supplementary Tables S2A and S2B in kumar et al., 2017[1]. As shown in Supplementary Table S2A [1]), 100 mg of AR contains approximately 1.5 mg of gallic acid.

Based on the experimentally established dose titration used in our previous studies (500 mg/kg body weight of AR), the corresponding intake of marker compounds is approximately 7.5 mg of gallic acid.

**HR-LCMS Characterization and Validation of Gallic Acid as the Primary Marker**

Unlike our 2017 study, which focused primarily on identifying enriched or novel metabolites, the present investigation employed high-resolution liquid chromatography–mass spectrometry (HR-LCMS) to achieve a more rigorous chemical characterization of the AR extract itself. This targeted analytical approach confirmed the presence of gallic acid (m/z ~169) and ellagic acid (m/z ~301) by matching their exact molecular masses and retention times with authenticated reference standards. Hence, Gallic acid was unambiguously confirmed in the AR extract through exact mass accuracy and chromatographic retention matching with the reference standard. These results establish high analytical confidence for gallic acid as a chemically validated marker.

**2.3.** ***Animal experiments:*** To test our hypothesis, serum, left ventricular tissue sections, tissue protein lysates etc. were used from the same cohort of animals as described previously by Kumar *et. al.*, 2017 [1]). Please see supplementary methods for details. Briefly, three-month-old male Wistar rats (180–200 g) were housed under standard conditions (12 h light/dark cycle, controlled temperature and humidity) with ad libitum access to synthetic chow and water. Pathological left ventricular cardiac hypertrophy was induced by transverse aortic constriction [TAC] using a titanium clip (small, Horizon, Catalog #1204) to reduce the aortic diameter by ~60%. Anaesthesia was administered using 3% isoflurane in 100% oxygen. Aortic constriction was verified via trans-thoracic 2D colour Doppler imaging. Biologically age-matched sham-operated controls (hereafter BA) underwent identical surgery without clip placement. Left ventricular hypertrophy was monitored from the ninth month of age.

*Two experimental arms were included:*

(i). *Ageing/Age-matched/Sham control Group with and without AR administration:* Sixteen rats were divided into biologically age-matched, sham-operated control groups, hereafter labelled as 'BA' and 'BA+AR' groups. Amalaki rasayana (AR) was prepared and characterized as previously described [1,2]. The BA+AR group received Amalaki rasayana (AR) orally (500 mg/kg, 5 days/week) starting the 9th month, and continued until the 21st month, ensuring at least 12 months of therapeutic exposure. 21 months of age in rats is considered equivalent to about 50 years of age in humans. The dosage used is much less than the proven safe in the clinical trials on normal human subjects (approx. 1g/kg body weight) [7]. Dose selection was based on prior pharmacokinetic analysis [1,2]; . It demonstrated traditional use for adequate bioavailability with no potential adverse effects, which collectively support the feasibility of our therapeutic approach and justify the current dosage used in our *in vivo* efficacy model [1,2;7]. There was no significant sole effect of the vehicle used to prepare AR, as reported in our previous studies with Dr. Vikas Kumar as the lead author [1,2].

(ii). *LVH/PO-CH Group with and without AR administration:* Sixteen rats underwent TAC to generate pressure-overload (PO) at the 3rd month. From 9th month onwards, one cohort received AR (hereafter labelled as PO-CH+AR'), while the other remained untreated (labelled as 'PO-CH'). Treatment continued until 21 months of age, ensuring at least 12 months of therapeutic exposure.

**2.4.** ***Echocardiography***

Echocardiographic evaluations were conducted at multiple times before and during treatment, up to 21 months of age, in both BA and PO-CH groups (using Philips HD-7 series Ultrasound Systems with Mitsubishi P95 S12 probe). To stabilize the heart rate during echo data acquisition, 75mg/kg ketamine and 5mg/kg xylazine was administered intraperitoneally for the sedation, and the animals were continuously monitored via ECG for any adverse changes prior to the final experiments. The anesthesia was maintained throughout the procedure. M-mode echocardiography was recorded at the parasternal short-axis view, and septal thickness, ejection fraction (EF), and fractional shortening (FS) were calculated from systolic and diastolic measurements. Echocardiographic data were recorded over several minutes, during which multiple M-mode images were captured to obtain the optimal view of the left ventricle and to minimize statistical variability. We typically recorded 3 to 5 cardiac cycles. The representative recordings shown in Figure 1B correspond to brief image segments of a cardiac cycle of a few seconds' duration. Left ventricular dimensions and ventricular hypertrophy were measured to assess the impact of AR on cardiac changes associated with ageing and pressure-overload induced hypertrophy. Interventricular septal thickness in systole (IVSs) and in diastole (IVSd), left ventricular end-systolic diameter (LVESD), left ventricular end-diastolic diameter (LVEDD), left ventricular posterior wall thickness in systole (LVPWs) and in diastole (LVPWd) were measured. It enabled calculation of several cardinal cardiac parameters. Diastolic Volume=1.047x (LVEDD)$^3$] this represents volume of the LV at the end of Diastole. Systolic Volume [1.047x (LVESD)$^3$] this represents volume of the LV at the end of Systole. Stroke Volume (SV) was calculated by the difference between the diastole and systole volume and expresses the volume of blood that exits the LV with each heartbeat. FS was calculated by the formula [(LVEDD-LVESD)/LVEDDx100)] which expressed the percentage change in the diameter of the LV during systole compared with diastole EF was calculated by the formula [(stroke volume/diastolic volume) x100], which expressed the percentage of efficiency of the LV in pumping blood. As all these parameters are already published in the previous work of the co-author [1], and as we have used the same cohort of animals for our study, therefore we have not provided the details in this manuscript to avoid repetition and redundancy.

##### 2.5. *Exercise Tolerance*

One month before study completion, all rats underwent a standardized treadmill protocol (30 min/day, 5 days/week). Tolerance was assessed by recording fatigue time and distance covered at incremental speeds of 5, 10, and 15 m/min.

##### 2.6. *Tissue Collection*

At the end of the study, rats were euthanized, and cardiac tissues were collected. Samples were either fixed in 10% neutral-buffered formalin for histology or stored in RNAlater for downstream analyses, including Western blotting and qPCR. Formalin-fixed tissues were processed for paraffin embedding.

##### 2.7. *Assessment of Cardiac Fibrosis, Calcification and Hypertrophy*

Cardiac fibrosis was assessed in 5 µm heart sections stained with Picrosirius Red (#365548, Sigma). Slides were incubated in stain for 1 h, rinsed with distilled and acidified water, dehydrated in graded ethanol and xylene, and mounted with DPX. Fibrotic areas were quantified using Fiji software and expressed as a percentage of total tissue area. Cardiomyocyte hypertrophy was evaluated by measuring myocyte dimensions in Fiji software. Calcification was assessed using 1% Alizarin S (pH 4.2) staining. Sections were incubated for 30 min at room temperature, rinsed, and then briefly immersed in acetone and an acetone–xylene (1:1) mixture before clearing in xylene and mounting with DPX. Calcium deposits appeared red to orange and were quantified using image analysis software.

**2.8.** ***Immunohistochemistry***

Slides were pre-baked at 55 °C for 15 min, de-paraffinized in xylene (2 × 20 min), and treated with chloroform. Rehydration was performed through a graded ethanol series (100% to 70%, 5 min each). Sections were washed in PBS (pH 7.4) and permeabilized with 0.01% digitonin in PBS for 30 min at room temperature. Blocking was done using 3% BSA and 2% donkey serum in PBS for 1.5 h, followed by overnight incubation with primary antibodies at 4 °C. After PBS washes, secondary antibodies were applied for 1 h at room temperature. Nuclei were counterstained with DAPI (5 min), and slides were mounted with 70% glycerol. Imaging and analysis followed previously established protocols. For probing surface signals, the tissues were not permeabilized with a detergent.

**2.9.** ***Cell Culture and Conditions***

Cardiomyoblasts were cultured in low-glucose DMEM supplemented with 10% FBS and antibiotics (complete medium). For experiments, cells were seeded in 6-well plates, 8-well chamber slides, or 96-well plates at densities of 0.1, 0.01, or 0.001 million cells/well, respectively. After 24 h, differentiation into myocytes was induced by switching to serum-reduced medium (DMEM with 1% FBS and antibiotics) for another 24 h. Cells were then treated in complete medium containing 5% FBS.

**2.10.** ***MTT Assay for Dose Estimation of AR***

Cardiomyoblasts (5,000 cells/well) were seeded and differentiated in 96-well plates. Post-differentiation, cells were treated with varying concentrations of AR (0, 5, 10, 50, 150, 200 µg/mL) in complete medium for 24–48 h. After treatment, cells were washed with warm 1× HBSS, and MTT (0.5 mg/mL) was added to each well. Plates were incubated in the dark at 37 °C for 4 h. Formazan crystals were solubilized with 100 µL DMSO/well and gently rocked for 15 min. Absorbance was recorded at 570 nm using a Varioskan multimode plate reader (Thermo Scientific, USA). Cell viability was expressed as the OD (optical density) in treated versus untreated cells.

**2.11.** ***Lysomotrophe assay for evaluating Gal-3 CRD functional inhibition by Gallic Acid (GA):***

Cardiomyocytes were seeded on 8-well chamber slides and pre-treated with galectin-3 carbohydrate recognition domain (CRD) inhibitors: 38 µM lactose (positive control) or 100 µM gallic acid (GA, test compound) for 24 hours. Following pre-treatment, cells were exposed to the lysosomal disrupting agent chloroquine (CQ) at a concentration of 50 μM for an additional 24 hours, with the inhibitors maintained throughout the treatment period. Cells were then fixed in 1.5% paraformaldehyde (PFA) and subjected to immunostaining using antibodies against lysosomal-associated membrane protein 1 (LAMP1) and full-length Gal-3 to assess co-localization.

At least 100 cells across five regions of interest (ROIs) were imaged per experimental condition, and data were analyzed and plotted for interpretation. In conditions where lysosomal integrity was maintained, minimal to no co-localization of Gal-3 with LAMP1 was expected . Conversely, cells pre-treated with GA and then exposed to CQ were expected to show a reduced binding of Gal-3 to damaged lysosomes (thereby reduced Gal-3-LAMP1 co-localization), indicating a functional inhibition of Gal-3's glycan-binding activity.

**2.12.** ***Measurement of intracellular free/cytosolic calcium:*** Cells were cultured in 35 mm confocal dished with glass bottom. At the experimental endpoint, cardiomyocytes from each group were incubated with 2 μM of free calcium sensing Fluo-3 AM dye, mixed with Pluronic F-127, for 20 minutes at 37°C in the dark. After incubation, the cells were washed with PBS to remove any excess dye. Fluorescent images were then captured from random regions of interest (ROIs) across three independent experimental sets. The resulting data were subsequently plotted for analysis and interpretation.

**2.13.** ***Human Galectin-3 ELISA***

Serum samples were diluted 1:1 and processed as per the manufacturer's protocol. Absorbance was measured at 450 nm using a Varioskan multimode plate reader (Thermo Scientific, USA). Data represent mean ± SD from three independent experiments.

**2.14.** ***Immunocytochemistry***

Cells were fixed in 1.5% paraformaldehyde for 20 min, permeabilized with 0.25% saponin for 20 min, and blocked for 1 h in PBS containing 3% BSA and 2% normal donkey serum. Primary antibodies were applied overnight at 4 °C. After PBS washes, cells were incubated with Alexa Fluor–conjugated secondary antibodies (1:200; Jackson ImmunoResearch) for 1 h at room temperature in the dark. Slides were mounted with DAPI-containing medium (Invitrogen), and images were captured using an Olympus confocal microscope (60× oil objective, NA 1.29). Imaging and analysis followed established protocols. Duolink *in situ* proximity ligation assay was performed according to the manufacturer's instructions (Sigma,

USA). Some of the resulting images are converted into the pseudo-colour gradient for better visualization.

#### 2.15. *Western Blotting*

Cells were washed with 1× PBS and lysed in RIPA buffer (1% NP-40, 0.1% SDS, 0.5% sodium deoxycholate, 150 mM NaCl, 50 mM Tris-HCl, pH 7.4, protease inhibitors). Lysates were sonicated and centrifuged at 14,000 rpm for 15 min at 4 °C; supernatants were collected. Please note that no housekeeping genes are reliable loading controls in pathologies, as their levels are significantly changed; therefore, the BCA assay was used to normalize concentrations. Also, for Western blots of serum or secretome samples (unbound proteins in the media and cell surface bound proteins), conventional intracellular loading controls such as GAPDH or β-actin cannot be used, as these proteins are not consistently present in extracellular fluids. To ensure equal loading, total protein staining methods (e.g., Ponceau S and Amido Black) can be employed. In the current study, while we did not include such staining, as it impacts the background during chemiluminescence imaging via iBright (Thermo Fisher Scientific, USA) or ImageQuant LAS500 (GE Healthcare Lifesciences) machines. Rather, the protein is quantitated via BCA assay and concentrations were carefully loaded, wherever applicable. However, for an additional loading control, 4μg of each protein sample was spotted as dots on the nitrocellulose membranes and post-sample drying, dot blots were stained with 0.1% amido black solution for 5 min and destained for another 5 min. The images were captured in BioRad GelDoc and quantified in Fiji image processing software. S.D. was obtained from three independent experiments and t-test was used to obtain test of significance. For human serum blots, equal sample volumes are resorted. Normalized concentrations were further assessed via dot blotting the BCA normalized samples on the nitrocellulose membrane and staining it with Amido black protein stain. Dot blot images were captured, and densitometric analysis matched with the BCA-assisted sample normalization procedure. Equal amounts of protein were mixed with SDS-β-mercaptoethanol sample buffer, heated at 95 °C for 5 min, separated by SDS-PAGE, and transferred to PVDF membranes. Membranes were blocked overnight in TBST (0.1% Tween-20) with 3% BSA, then incubated with primary antibodies diluted in blocking buffer for 1 h at room temperature or overnight at 4 °C. After washes, membranes were probed with HRP-conjugated secondary antibodies and developed using Clarity ECL (Bio-Rad). Signals were detected with the LAS-500 imaging system. Densitometric analysis was performed in Fiji software, and individual values were graphically presented.

**2.16.** ***Plasmid and siRNA Transfections:***

Cells at 70% confluence were incubated in serum-free OPTIMEM media for 2 h before transfection. 100 pM of siRNA or high-purity plasmids (0.5 µg) were complexed with Lipofectamine LTX Plus reagent at room temperature for 15–30 min. The mixture was then diluted with 350 µL OPTIMEM and added dropwise to cells with gentle swirling. After 4–6 h incubation at 37 °C, 5% $CO_2$, the transfection medium was replaced with complete growth medium. Cells were cultured for 24–72 h for optimal plasmid expression or siRNA mediated translational inhibition.

**2.17.** ***Rhodamine Phalloidin,WGA, Pyronin Y and Amido black staining***

Fixed cells, prepared for immunocytochemistry if needed, were incubated with Rhodamine phalloidin (2 µg/mL) for 45 min to visualize F-actin stress fibers. For WGA staining, cells were treated with WGA (5 µg/mL) for 45 min, followed by brief washes. Cells were counterstained with Hoechst (5 µg/mL, 5 min) and mounted for imaging. Pyronin Y stains DNA and RNA through its positive charge that interacts with the negatively charged phosphate backbone of the nucleic acid. The cytoplasmic Pyronin Y signal can be extracted first merging the images of the Hoechst nuclear stain with fluorescent Pyronin signal and drawing ROI around the nuclei and the cell periphery, and finally subtracting the nuclear intensity with the whole cell intensity. Since RNA is cytoplasmically localised, RNA intensity is thus derived. Amido black stain (0.1%) was used to visualize total protein content of the tissue.

**2.18.** ***Galectin-3 Docking and Simulation Studies with AR Components***

The carbohydrate-binding site of human Galectin-3 (Gal-3, PDB ID: 3ZSK) and a homology-modelled structure of rat Gal-3 were prepared using the Schrödinger Protein Preparation Wizard (Schrödinger Release 2021-3: Glide, Schrödinger, LLC, New York, NY, 2021). The aim was to determine the binding conformations and energies of 18 compounds identified in the Amalaki rasayana (AR) extract. Receptor structures were energy-minimized with the OPLS_2005 force field, constraining heavy atoms to deviate no more than 0.3 Å from their original positions. For molecular docking, Glide [10,11,12] was used to generate receptor grids centered on the ligand-binding site defined by the human Gal-3 crystal structure, ensuring accurate representation of the binding pocket. These grids served as docking targets for the selected compounds. The LigPrep module generated 3D structures of the 18 compounds, optimized their geometry, and produced all possible stereoisomers and ionization states. The energy-minimized ligands were docked into the receptor grids using Glide in standard precision (SP) mode with pre-defined constraints. The docking protocol predicted the preferred conformation and orientation of each ligand within the receptor's binding site. GlideScore below -4.5 is generally considered good. The pose with the lowest GlideScore (GScore), i.e. was selected as the best-docked conformation and used for subsequent interaction analysis. A GlideScore estimates the binding affinity between a ligand and a protein receptor. It is based on an empirical scoring function that accounts for various physical interactions, such as hydrogen bonds, van der Waals forces, and hydrophobic effects. A more negative GlideScore generally indicates a stronger and more stable predicted binding.

#### Information on reagents used:

| Reagents/ Resources | Experiment and Dilutions Used | Sources | Cat. No. |
|---|---|---|---|
| **Primary Antibodies** | | | |
| Galectin-3 Mouse Monoclonal Antibody | (1:500); Immunoblot | Santa Cruz Biotechnology | (B2C10) sc32790 |
| Rabbit Galectin-3 Antibody C-Epitope | (1:1000); Immunoblot<br>(1:150); ICC/IHC<br>(1:25), 4µg; IP | FabGennix | CBP-112 AP |
| Rabbit Galectin-3 Antibody N-Epitope | (1:1000); Immunoblot<br>(1:150); ICC/IHC<br>(1:25), 4µg; IP | FabGennix | CBP-101 AP |
| NPPA(ANP) Rabbit pAb, | (1:5000); Immunoblot | ABclonal | A1609 |
| Albumin Rabbit pAb | (1:1000); Immunoblot | ABclonal | A1363 |
| Anti-Cyclophilin A Antibody (PPIA), | (1:5000); Immunoblot | Abcam | AB58144 |
| pSER-Ab Mouse | (1:1000); Immunoblot<br>(1:200); ICC/IHC | AbboMax | 500-020 |
| Rabbit Anti-GATA4 pAb, | | | ab84593 |
| NFATC4 Rabbit pAb | (1:1000); Immunoblot | ABclonal | A17511 |
| CaMKII delta Rabbit mAb | (1:1000); Immunoblot | ABclonal | A9196 |
| MEF2A Rabbit pAb | (1:1000); Immunoblot | ABclonal | A12059 |
| MYH7 Ab | (1:400); Immunoblot | ABclonal | A24664 |
| LDHA (C4B5) Rabbit mAb | (1:1000); Immunoblot | Cell Signaling Technology | 3582S |
| HMGB | (1:1000); Immunoblot | Abcam | ab227168 |
| ACTN2 Rabbit pAb | (1:10); Immunoblot | ABclonal | A3718 |
| TMOD1 Rabbit pAb | (1:1000); Immunoblot | ABclonal | A4160 |
| OBSCN Rabbit pAb | (1:000); Immunoblot | ABclonal | A18110 |
| SERCA2 | (1:2000); Immunoblot | Cell Signaling Technology | Ab 4388S |
| Anti-Galectin-3, Rat, Clone M3/38 | (1:25), 4µg; IP | Sigma-Aldrich | MABT51 |
| Anti-GFP Ab, Mouse, | (1:26.67), 3µg; IP | Roche | 1181446000 |
| Rabbit Galectin 3 Antibody FITC | (1:150); ICC/IHC | FabGennix | CBP35-FITC |
| Rabbit Galectin 3 Antibody FITC | (1:150); ICC/IHC | FabGennix | CBP35.c-FITC |
| α-Smooth Muscle Actin | (1:200); IHC | ABclonal | A7248 |

| | | | |
|---|---|---|---|
| (ACTA2) Rabbit pAb | | | |
| CD68 Rabbit pAb | (1:200); IHC | ABclonal | A13286 |
| Anti-PTRF antibody | (1:200); ICC | Abcam | ab48824 |
| Mouse Monoclonal Caveolin-1 antibody | (1:400); ICC | Abcam | ab17052 |
| Anti-gamma H2A.X (phospho S139) antibody | (1:300); ICC | Abcam | ab11174 |
| ROCK1 Rabbit mAb | (1:200); ICC | ABclonal | A11158 |
| Anti-RhoA Mouse mAb | (1:200); ICC | Cytoskeleton | ARH04 |
| Rabbit Anti-GATA4 pAb | (1:200); ICC | Abcam | ab84593 |
| NFATC2 Rabbit pAb | (1:500); ICC | ABclonal | A3107 |
| MEF2A Rabbit pAb | (1:500); ICC | ABclonal | A12059 |
| Lamin A Antibody (4A58) | (1:200); ICC | Santa Cruz Biotechnology | sc-71481 |
| Integrin β1 (P5D2) | (1:200); ICC | Santa Cruz Biotechnology | sc-13590 |
| β2-AR (H-73) rabbit pAb | (1:200); ICC/IHC | Santa Cruz Biotechnology | sc-9042 |
| Rabbit Galectin 3 Antibody BIOTIN | (1:150); ICC/IHC | FabGennix | CBP35.c-BIOTIN |
| Galectin 3 Antibody BIOTIN | (1:150); ICC/IHC | FabGennix | CBP35-BIOTIN |
| WGA FITC | (10μg/ml); IHC | | |
| Rhodamin Phalloidin | (2μg/ml); ICC | Invitrogen | R415 |
| Hoechst 33342 (bisBenzimide H 33342 trihydrochloride) | (5μg/ml); ICC/IHC | Sigma | 14533 |
| **Secondary Antibodies** | | | |
| Peroxidase conjugated affinitypure donkey anti Mouse IgG | (1:10,000); Immunoblot | Jackson ImmunoResearch | 715-035-150 |
| Peroxidase conjugated AffiniPure Donkey Anti-Rabbit IgG (H+L) | (1:10,000); Immunoblot | Jackson ImmunoResearch | 711-035-152 |
| Alexafluor 647 conjugated affinitypure donkey anti Rabbit IgG | (1:200); ICC/IHC | Jackson ImmunoResearch | 711-605-152 |
| Alexafluor 488 conjugated affinitypure donkey anti Rabbit IgG | (1:200); ICC/IHC | Jackson ImmunoResearch | 711-545-152 |
| Alexafluor 647 conjugated donkey anti Mouse IgG, | (1:200); ICC/IHC | Life Technologies | A31571 |
| Alexafluor 594 conjugated affinitypure donkey anti Rabbit IgG | (1:200); ICC/IHC | Jackson ImmunoResearch | 711-585-152 |
| DyLight™ 405 AffiniPure Donkey Anti-Mouse IgG (H+L) | (1:200); IHC | Jackson ImmunoResearch | 715-475-150 |
| Alexafluor 488 conjugated affinitypure donkey anti Mouse IgG | (1:200); ICC/IHC | Jackson ImmunoResearch | 715-545-150 |
| Rhodamine Red X Conjugated streptavidin | (1:200); ICC/IHC | Jackson ImmunoResearch | 016-290-084 |

| | | | |
|---|---|---|---|
| **Kits** | | | |
| Rat Gal-3 ELISA Kit | ELISA | Thermo | xERLGALS3 |
| Quantikine® ELISA Human Galectin-3 | ELISA | Quantikine | DGAL30 |
| Duolink PLA Kit: In Situ PLA Anti Rabbit Minus | PLA Assay | Sigma | DUO92005 |
| Duolink PLA Kit: In Situ PLA Anti Mouse Plus | PLA Assay | Sigma | DUO92001 |
| IP Kit – Dyna Beads - Protein A | 33.33µL (1mg); IP | Life Technologies | 10006D |
| Lipofectamine™ LTX Reagent with PLUS™ Reagent | Plasmid Transfection | Invitrogen | A12621 |
| **Other Reagents** | | | |
| Name | Working concentration & application | Manufacturer | Catalogue no. |
| Gallic Acid | 100µM/ml | Sigma | G7384-250G |
| Amalaki Rasayana | 100µg/ml | Arya Vaidya Sala, Kottakkal | |
| Beta-Lactose | 34mM | Sigma | L3750 |
| Pyronin Y | 0.1% IHC | Sigma | P9172 |
| Amido Black | 0.1% IHC | HIMEDIA | MB165 |
| Direct Red-80 (Sirius Red) | 0.1% IHC | Sigma | 36-554-8 |
| Alizarin S Red solution | 1% IHC | Merck | 2003999 |
| Thiazolyl Blue Tetrazolium Bromide (MTT) | 0.5mg/ml | Sigma | M2128 |
| 2x Laemmli Sample Buffer | 1x | Biorad | 1610737 |
| Bicinchoninic Acid (BCA) | Protein Estimation | Sigma | B9643 |
| Copper(II) sulphate | Protein Estimation | Sigma | C2284 |
| Clarity Western ECL Substrate | Immunoblot | Biorad | 170-5060 |
| Protein Marker (10-180kDa) | 1-3µL/well; Immunoblot | | Taurus-(TI-MB-19001) |
| DPX Mountant | 100%; IHC | Qualigens | Q18404 |
| Digitonin | 0.01%; IHC | Sigma | D141 |
| Saponin from Quilaja bark | 0.25%; ICC | Sigma | S7900 |
| Triton X-100 | 0.1%; ICC/IHC | Sigma | T9284-500ML |
| Glycerol | 70% ;ICC/IHC | Sigma | G5516 |
| Bovine serum Albumin (BSA) | 3%; Immunoblot<br>3%; ICC | Sigma | A7906 |
| Normal donkey serum | 2%; ICC/IHC | Jackson ImmunoResearc h | 017-000-121 |
| Paraformaldehyde (PFA) | 1.5% ;ICC | Merck | 30535-89-4 |
| BAPTA (1,2-Bis-2-amino-5-methylphenoxy)ethane-N,N,N′,N′-tetraacetic acid tetrakis-acetoxymethyl-ester) | 1µM | Sigma | 16609 |
| **Cell Culture Reagents** | | | |

| | | | |
|---|---|---|---|
| Fetal Bovine Serum EU Approved, Heat Inactivated | 1%, 5%, 10% | HIMEDIA | RM9955-500ML |
| Dulbecco's Modified Eagle Medium (DMEM) Low Glucose | Cell culture | Gibco | 31600-034 (10 x 1 L) |
| Dulbecco's Modified Eagle Medium (DMEM) High Glucose | Cell culture | Gibco | 12100046 (10 x 1 L) |
| CO2 Independent Medium (1X) | PO Treatment | Gibco | 18045-088 |
| Antibiotic-Antimycotic (100X) | 2% | Gibco | 15240062 (100 mL) |
| Trypsin – EDTA Solution 1X | 3ml | HIMEDIA | TCL033-500ML |

**Cell Lines**

| | |
|---|---|
| H9c2(2-1) | ATCC |
| HEK392 | ATCC |

**Plasmid Constructs**

| | | |
|---|---|---|
| WT hGal3 plasmid | GAL-3 WT GFP | Addgene and **Zellebiotech** |
| Gal-3 R186S plasmid | GAL-3 Mut | |

**Plasmid Synthesized with Zellebiotech**

| Construct | Sequence |
|---|---|
| 1. Gal-3 Mutant S6A Human galectin-3 S6A mutant, serine at position 6 is changed to arginine. | MADNFRLHDALSGSGNPNPQGWPGAWGNQPAGAGGYPGASYPGAYPGQAPPGAYPGQAPPGAYPGAPGAYPGAPAPGVYPGPPSGPGAYPSSGQPSATGAYPATGPYGAPAGPLIVPYNLPLPGGVVPRMLITILGTVKPNANRIALDFQRGNDVAFHFNPRFNENNRRVIVCNTKLDNNWGREERQSVFPFESGKPFKIQVLVEPDHFKVAVNDAHLLQYNHRVKKLNEISKLGISGDIDLTSASYTMI |
| 2. Gal-3 Triple Mutant Human galectin-3 triple mutant A144S, ASN160K and ASN 164 K; arginine mutated to serine and asparagine mutated to lysine | MADNFSLHDALSGSGNPNPQGWPGAWGNQPAGAGGYPGASYPGAYPGQAPPGAYPGQAPPGAYPGAPGAYPGAPAPGVYPGPPSGPGAYPSSGQPSATGAYPATGPYGAPAGPLIVPYNLPLPGGVVPRMLITILGTVKPNANSIALDFQRGNDVAFHFKPRFKENNRRVIVCNTKLDNNWGREERQSVFPFESGKPFKIQVLVEPDHFKVAVNDAHLLQYNHRVKKLNEISKLGISGDIDLTSASYTMI |
| 3. Gal-3-C-terminal Human Galectin-3 C-terminal domain cloned in mammalian expression vector (tagged with GFP. C-terminal domain is from 150-250 amino acids) | MQRGNDVAFHFNPRFNENNRRVIVCNTKLDNNWGREERQSVFPFESGKPFKIQVLVEPDHFKVAVNDAHLLQYNHRVKKLNEISKLGISGDIDLTSASYTMI |
| 4. Gal-3-N-terminal Human Galectin-3 N- | MADNFSLHDALSGSGNPNPQGWPGAWGNQPAGAGGYPGASYPGAYPGQAPPGAYPGQAPPGAYPGAPGAYPGAPAPG |

| | |
|---|---|
| terminal domain cloned in mammalian expression vector (tagged with GFP. C-terminal domain is from 1-150 amino acids) | VYPGPPSGPGAYPSSGQPSATGAYPATGPYGAPAGPLIVP YNLPLPGGVVPRMLITILGTVKPNANRIALDF |
| **Instruments & Software** | |
| Varioskan LUX Multimode Microplate Reader | |
| ImageQuant™ LAS 500 | |
| Olympus Confocal Microscope | |
| FiJi image processing software | |

##### Human Serum Sample Details

- **Samples Diagnosed with Hypertrophic Cardiomyopathy disease**

| Donor ID | Sample ID | Sex | Race | Age | Diagnosis | Medications |
|---|---|---|---|---|---|---|
| 2163402 | HMN512435 | Male | Caucasian | 62 | Type 2 Diabetes, Hypertension(HTN), Chronic Kidney Disease, Dyslipidemia, Benign Prostatic Hyperplasia (BPH), Coronary Artery Disease (CAD), **Hypertrophic Cardiomyopathy**, Hyperlipidemia; | Fenofibrate 160mg, Acyclovir 400mg, Finasteride 5mg, Atorvastatin Calcium 40mg, Simvastatin 40mg, Ranitidine 150mg, Bystolic 20mg, Benicar 12.5mg, Amlodipine Besylate 2.5mg, Proscar 5mg, Aspirin 81mg |
| 2166592 | HMN752093 | Female | Caucasian | 76 | Congestive Heart Failure (CHF), Hypertension (HTN),Hypercholesterolemia, Paroxysmal Atrial Fibrillation(AF), Obstructive Hypertrophic Cardiomyopathy, ParoxysmalAtrial Fibrillation (AF), Chronic Diastolic Heart Failure, Anemia(Iron Deficiency), Hypertension(HTN), Hypercholesterolemia, Diverticulitis of Large Intestine with Perforation and Abscess,Vitamin D Deficiency,Gastroesophageal Reflux Disease (GERD), | Amiodarone HCI 100mg,Atorvastatin Calcium 20mg, Pantoprazole Sodium 40mg, Warfarin Sodium 6mg,Losartan Potassium 100mg,Lasix 40mg, AmlodipineBesylate 5mg, Folic Acid 1mg, Ferrous Sulfate Iron 200mg,Vitamin D 50mcg,Acetaminophen 500mg,Vitamin B Complex, Nystatin 1,Multivitamin, Gabapentin |

| | | | | | | |
|---|---|---|---|---|---|---|
| | | | | | **Hypertrophic Cardiomyopathy**, Hyperlipidemia (HLD), Osteoarthritis (OA), Diverticulitis with Perforation; | |
| 2202569 | HMN502022 | Male | Caucasia n | 63 | Myocardial Infarction (MI),Hypertension (HTN), Hyperlipidemia (HLD), Type 2Diabetes, Chronic Kidney Disease (CKD), Dyslipidemia, atherosclerotic CardiovascularDisease, Benign localized hyperplasia of prostate withouturinary obstruction and other lower urinary tract symptoms,**Hypertrophic HMN cardiomyopathy**, Coronary Arthritis Disease; | Finasteride 5mg |
| 2198688 | HMN611152 | Male | Caucasia n | 63 | Chronic Kidney Disease, Essential hypertension, Hyperlipidemia, Type 2 diabetes mellitus, Dyslipidemia, Arteriosclerotic heart disease, Type 2 diabetes mellitus with diabetic nephropathy, without long-term current use of insulin, **Hypertrophic cardiomyopathy**, Genital herpes, Gastroesophageal reflux disease, Coronary artery disease; | Fenofibrate 160 MG, Acyclovir 800 MG, Finasteride 5 MG,Atorvastatin Calcium 40 MG, Simvastatin 40 MG, Bystolic 20MG, Benicar HCT 4012.5 MG,Sildenafil Citrate 20 MG, Systolic 10 MG, Amlodipine Besylate 2 .5 MG, Proscar SMG, Aspirin Adult Low Dose81 MG |
| 2201285 | HMN711025 | Male | Caucasia n | 63 | Chronic Kidney Disease, Hypertension, Hyperlipidemia, Type 2 Diabetes, Dyslipidemia, Arteriosclerotic Heart Disease, Benign localized hyperplasia of prostate without urinary obstruction and other | Ranitidine 150mg, Amlodipine Besylate 2.5mg, Proscar 5mg, Acyclovir 800mg, Aspirin81mg, Metformin 500mg, Losartan Potassium 50mg,Atorvastatin Calcium 40mg, |

| | | | | | | |
|---|---|---|---|---|---|---|
| | | | | | lower urinary tract symptoms, Arteriosclerotic heart disease, Nephropathy, **Hypertrophic Cardiomyopathy**, Gastroesophageal reflux disease, Genital Herpes, Coronary Artery Disease; | Finasteride 5mg, Sildenafil Citrate 20mg, Fenofibrate 160mg, Acyclovir 400mg,Simvastatin 40mg |

**Healthy Young and Old Human Donor Samples**

| Lot Number | Gender | Age | Race | Volume | DOM | EXP |
|---|---|---|---|---|---|---|
| 46545-46 | M | 26 | B | 10 ml | 11/28/2023 | 11/28/2026 |
| 46545-47 | M | 29 | H | 10 ml | 11/28/2023 | 11/28/2026 |
| 46545-48 | M | 30 | B | 10 ml | 11/28/2023 | 11/28/2026 |
| 46545-49 | M | 36 | B | 10 ml | 11/28/2023 | 11/28/2026 |
| 46545-40 | M | 21 | B | 10 ml | 11/28/2023 | 11/28/2026 |
| HMN1132955 | M | 57 | B | 10 ml | 06/14/2023 | 06/14/2026 |
| HMN1132957 | M | 67 | H | 10 ml | 06/14/2023 | 06/14/2026 |
| HMN1204367 | M | 56 | N | 10ml | 12/13/2023 | 12/13/2026 |
| HMN1204371 | M | 60 | C | 10ml | 12/13/2023 | 12/13/2026 |
| HMN1204388 | M | 55 | B | 10ml | 12/13/2023 | 12/13/2026 |

#### Section C: Uncropped Dot and Western blot Images

##### I. Uncut dot and western blots for figures are as follows:

#### For Figure 2

A. **Dot Blot Galectin-3 full length, Fig.2A**

Uncut Dot Blot Image

| | BA | BA+ AR | PO CH | PO-CH AR |
|---|---|---|---|---|

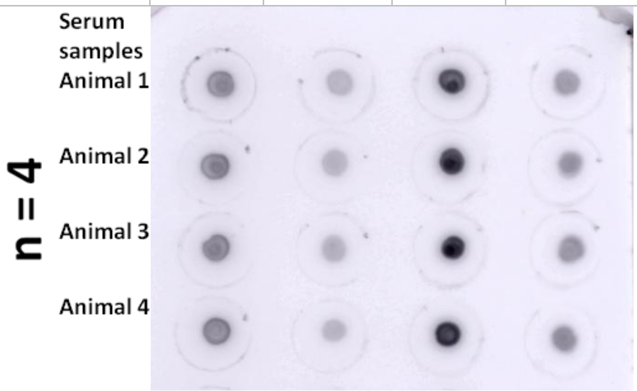

Representative Dot Blot Image (Inverted)

| | BA | BA+ AR | PO CH | PO-CH AR |
|---|---|---|---|---|

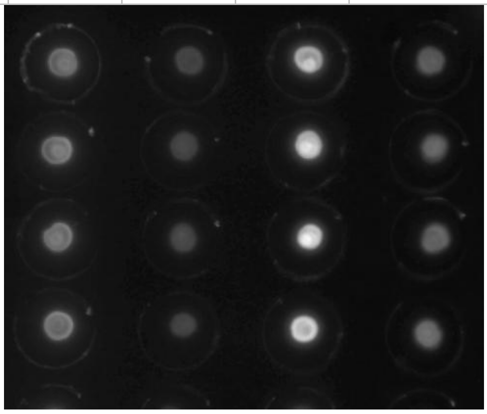

**B.Western blot with Gal-3 N-epitope-specific antibody, Fig.2B**

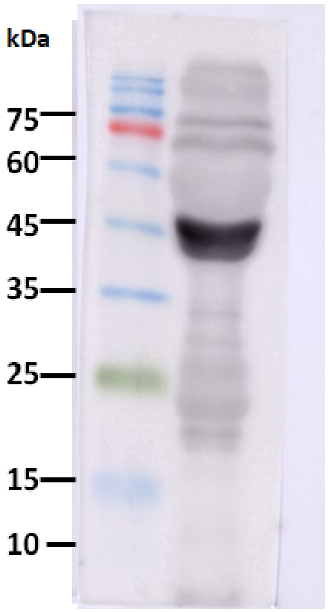

Representative blot shown in Figure2B

Three independent serum samples probed with Gal 3 N-specific antibody

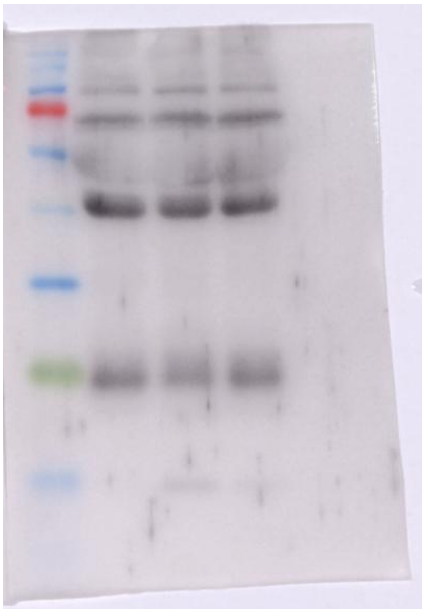

**C. Western blot with Gal-3 C-epitope-specific, Fig.2B**

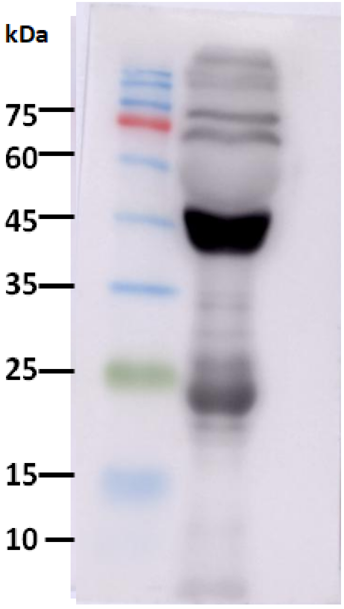

Representative blot shown in Figure 2B

Three independent serum samples probed with Gal 3 C-specific antibody

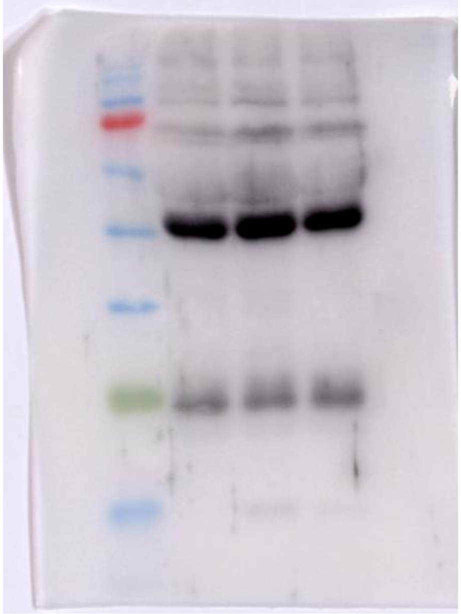

For fig 2B, Original uncut and uncropped dot blot images with three replicates

**A. Western blot with Gal-3 C-epitope antibody, Fig.2 C**

Blot 1

Representative blot shown in Figure 2C

| Marker | | BA | BA+ AR | PO-CH | PO-CH + AR |
|---|---|---|---|---|---|

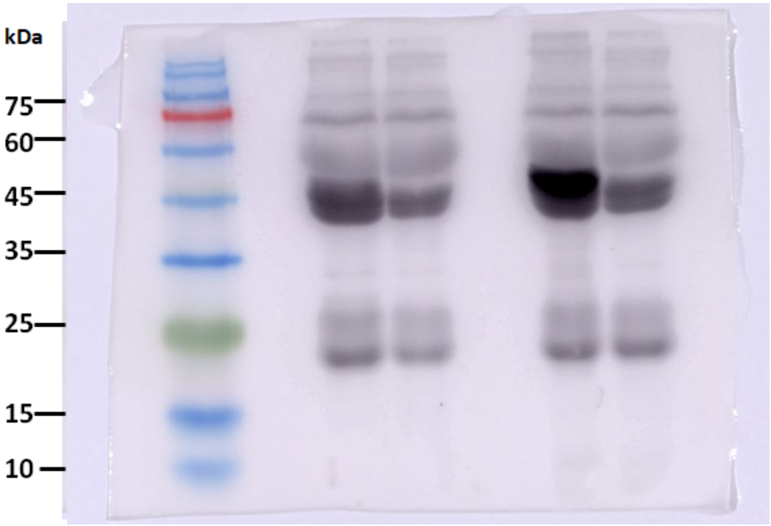

Blot 2

| Marker | | BA | BA+ AR | PO-CH | PO-CH + AR |
|---|---|---|---|---|---|

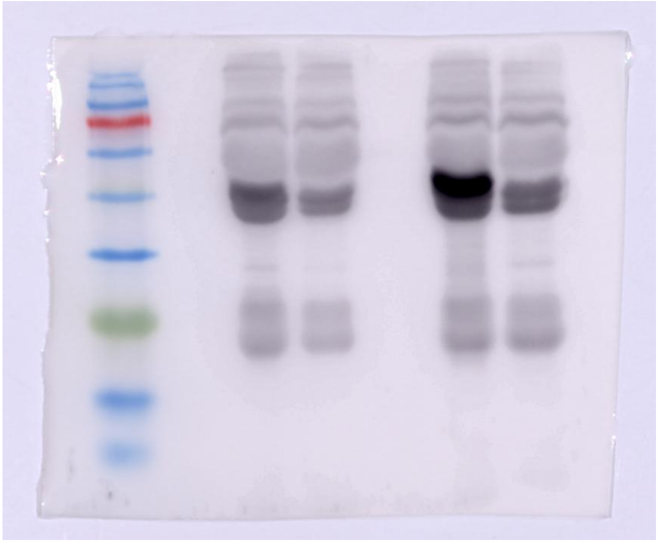

Blot 3

| BA | BA+ AR | PO-CH | PO-CH + AR |
| --- | --- | --- | --- |

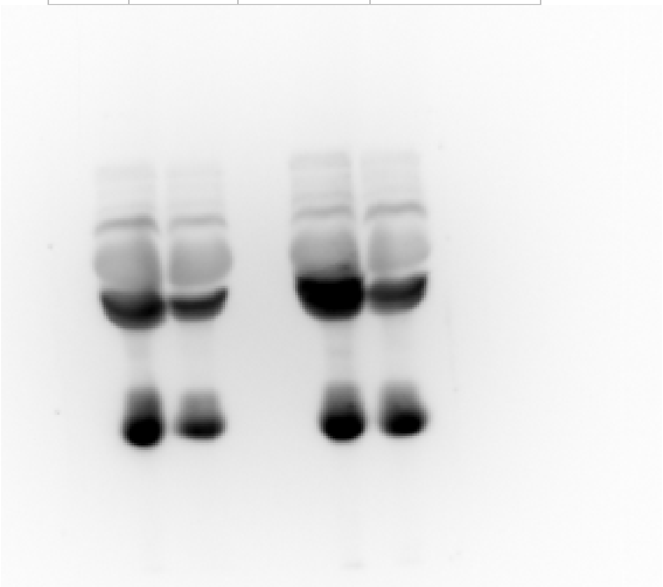

Blot 4

| Marker | BA | BA+ AR | PO - CH | PO-CH + AR |
| --- | --- | --- | --- | --- |

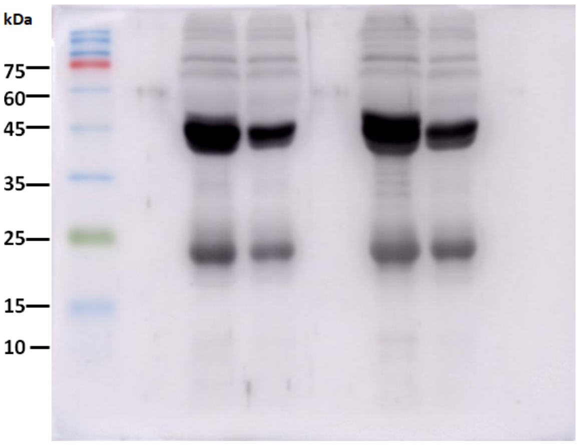

Blot 5

| Marker | BA | BA+ AR | PO - CH | PO-CH + AR |
|---|---|---|---|---|

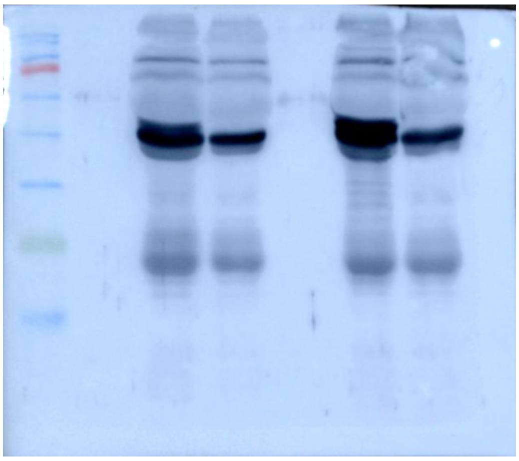

Blot 6

| Marker | BA | BA+ AR | PO - CH | PO-CH + AR |
|---|---|---|---|---|

.

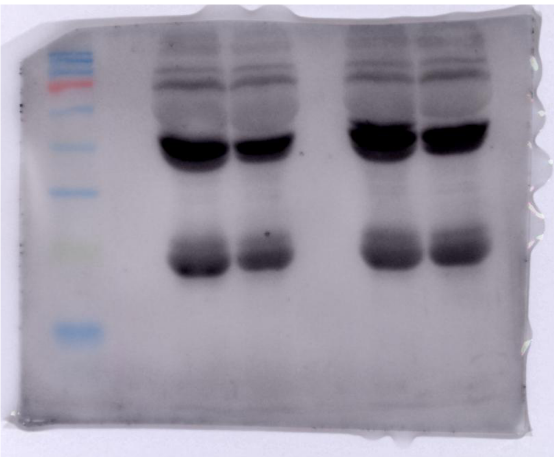

For Fig. 2C, Original Uncut and uncropped dot blot images with three replicates

**(A) Dot blot Original Images for Figure 2A along with the represented (inverted images). (B-C) Uncut Western Blot images shown in Figure 2. The original images are captured in LAS500 and shown in the inverted mode for better visualization**

### <u>For Figure 4</u>

#### A. Western blot Images for Fig. 4D

Blot 1

Representative

| M | NO PO | NOP O+A R | PO | PO +AR |
|---|---|---|---|---|

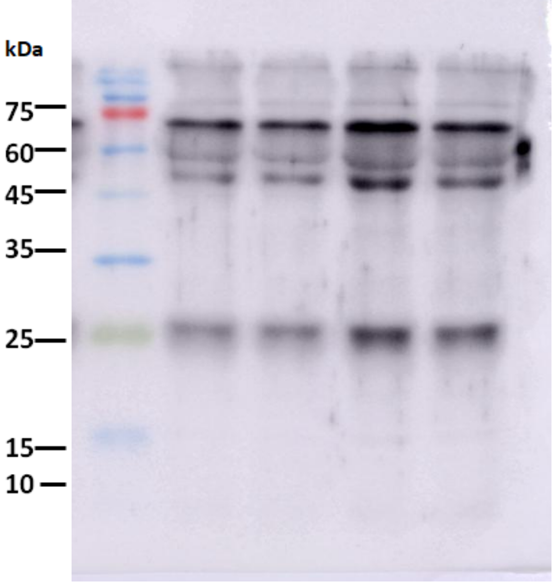

Blot 2

| M | NO PO | NO PO +A R | PO | PO +AR |
|---|---|---|---|---|

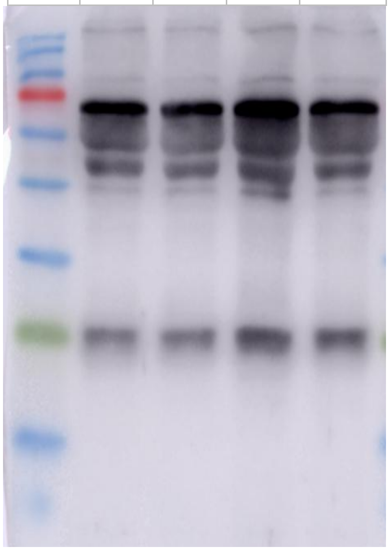

Blot 3

| NO PO | NO PO+ AR | PO | PO +AR |
|---|---|---|---|

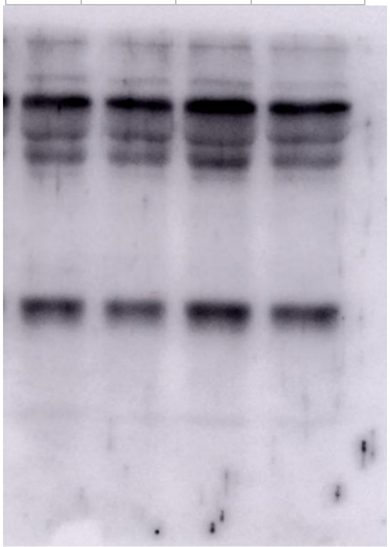

#### B. Western blot Images for Fig. 4F

Blot1 Representative

Blot2

| M | NO PO | NO PO +A R | PO | PO +A R | M | NO PO | NO PO +A R | PO | PO +A R |
|---|---|---|---|---|---|---|---|---|---|

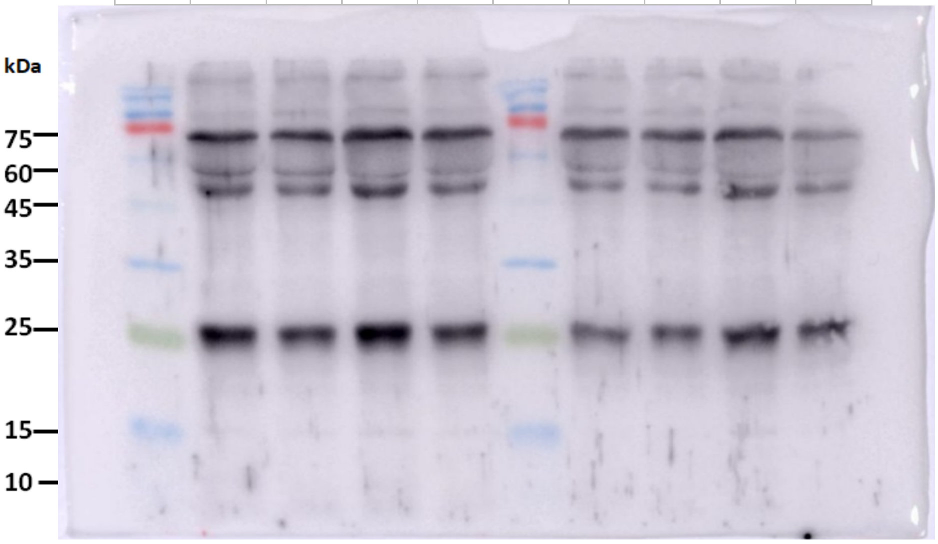

Blot 3 Blot 4

| M | NO PO | NOP O+ AR | PO | PO + AR | N O P O | NOP O+A R | PO | PO +AR |
|---|---|---|---|---|---|---|---|---|

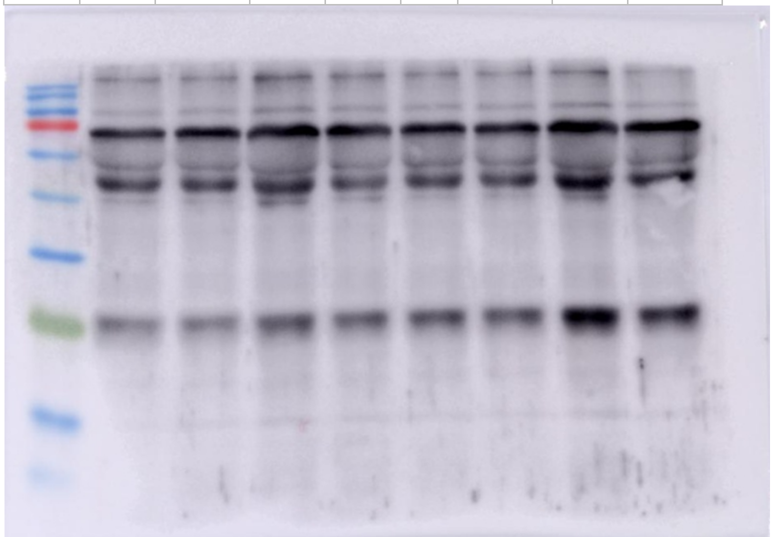

#### C. Western blot Images for Fig. 4H

Blot1 Blot2

Representative

| M | N O P O | NO PO +A R | P O | PO +AR | M | NO PO | NO PO +A R | PO | PO +A R |
|---|---|---|---|---|---|---|---|---|---|

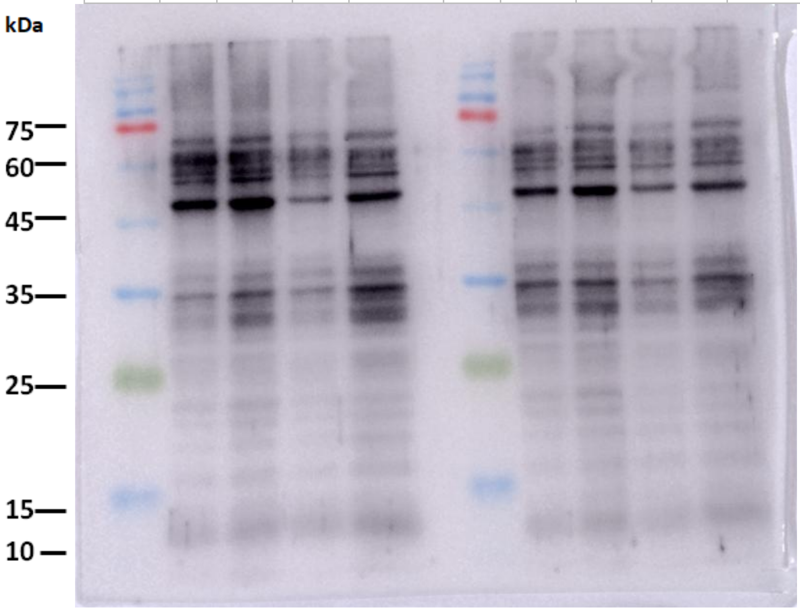

Blot 3 Blot 4

| M | NO PO | NO PO +AR | PO | PO +AR | M | NO PO | NO PO +AR | PO | PO +AR |
|---|---|---|---|---|---|---|---|---|---|

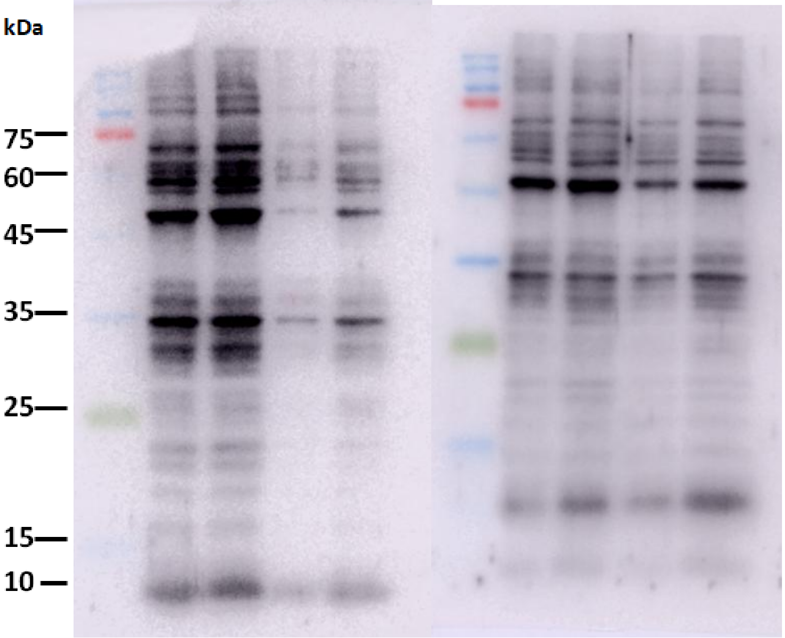

**Western blot uncut Original Images . 3-4 independent samples are probed in each group. The original images are captured in LAS500 and shown in the inverted mode for better visualization.**

### For Figure 5

**A. Western blot Images for Fig.5D**

Blot 1

Representative

| M | NO PO | NO PO +A R | PO | PO +AR |
|---|---|---|---|---|

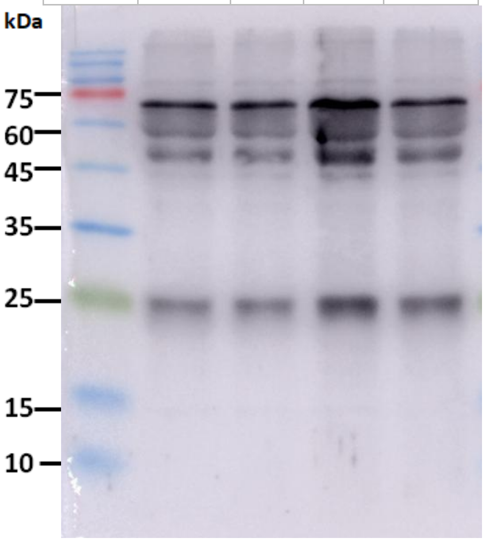

Blot 2

| M | NO PO | NO PO +A R | PO | PO +A R |
|---|---|---|---|---|

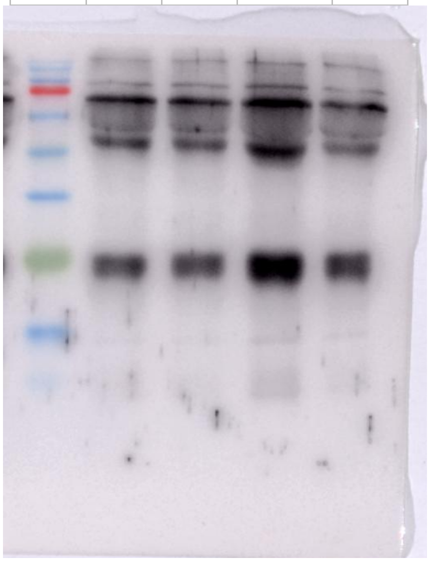

Blot 3

| M | NO<br>PO | NO<br>PO<br>+A<br>R | PO | PO<br>+A<br>R |
|---|---|---|---|---|

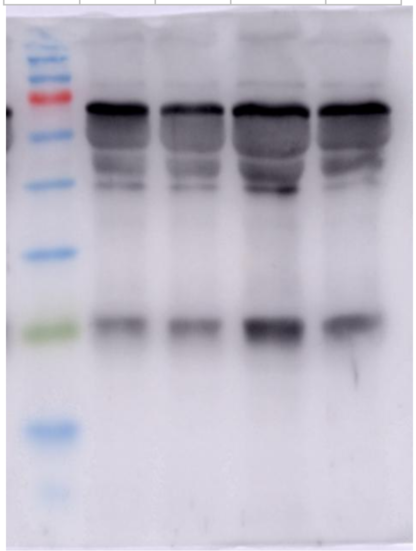

**B.Western blot Images for Fig. 5F**

Blot 1 | Blot 2

| M | NO<br>PO | NO<br>PO<br>+G<br>A | PO | P<br>O<br>+<br>G<br>A | NO<br>PO | NO<br>PO<br>+G<br>A | PO | P<br>O<br>+<br>G<br>A | M |
|---|---|---|---|---|---|---|---|---|---|

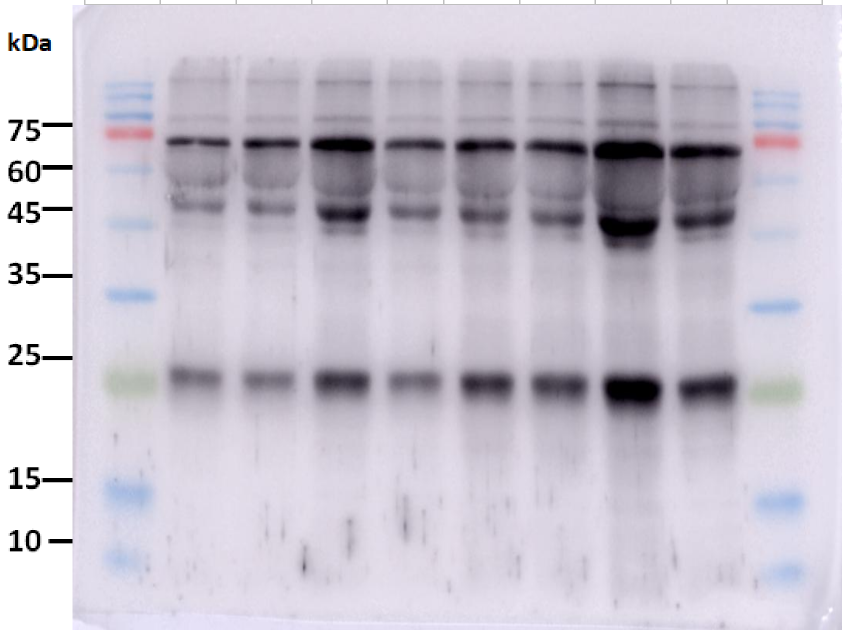

Blot 3

| M | NO PO | NOPO +GA | PO | PO +GA |
| --- | --- | --- | --- | --- |

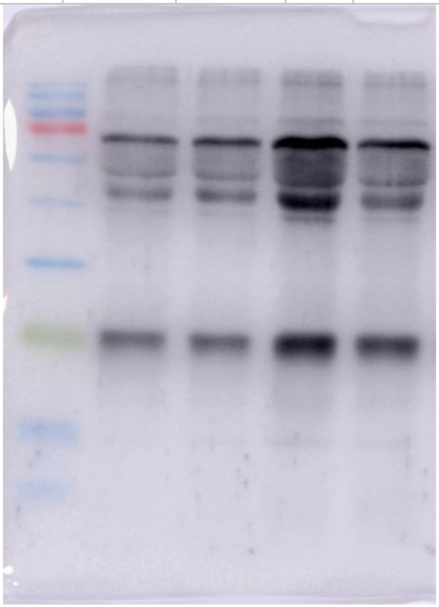

**C. Western blot Images for Fig.5H**

Blot 1

Representative

| M | NO PO | NOPO +GA | PO | PO +GA |
| --- | --- | --- | --- | --- |

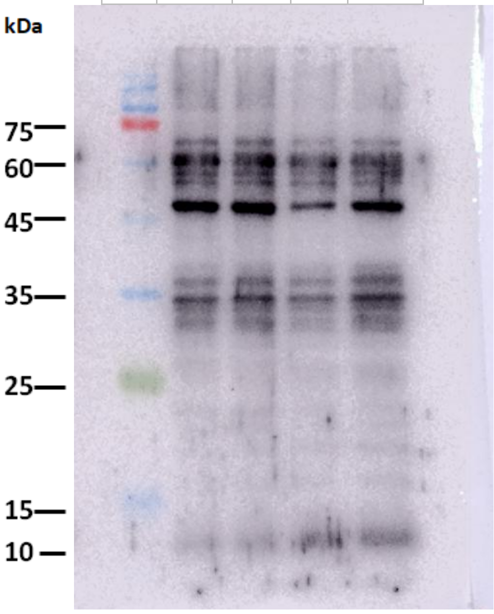

Blot 2 Blot 3

| M | NOPO | NOPO+GA | PO | PO +AR | M | NOPO | NOPO+GA | PO | PO+GA |
|---|---|---|---|---|---|---|---|---|---|

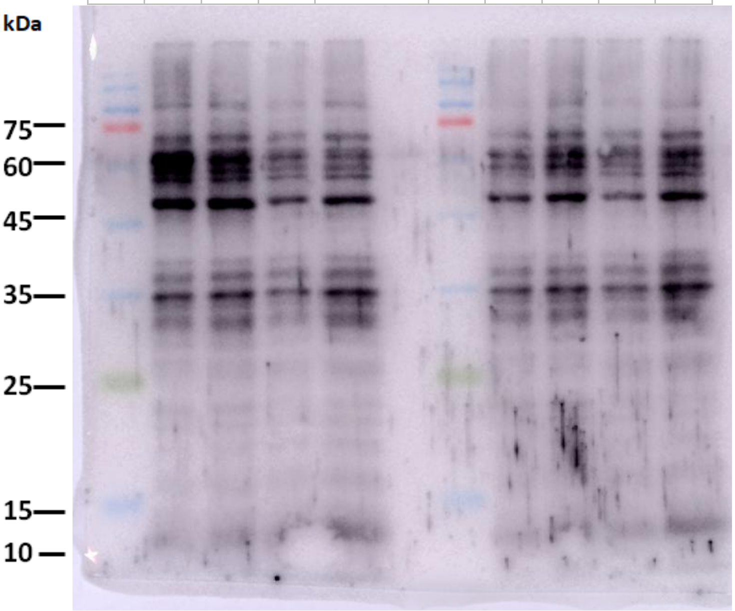

**Western blot uncut Original Images . Three independent samples are probed in each group. The original images are captured in LAS500 and shown in the inverted mode for better visualization.**

### For Figure 6

#### A. Western blot Images for Fig.6 J

Blot 1
Representative

Blot 2

| M | P<br>O | PO+<br>GA | P<br>O | PO<br>+GA | PO<br>+G<br>A | P<br>O | PO+<br>GA | P<br>O | M |
|---|---|---|---|---|---|---|---|---|---|

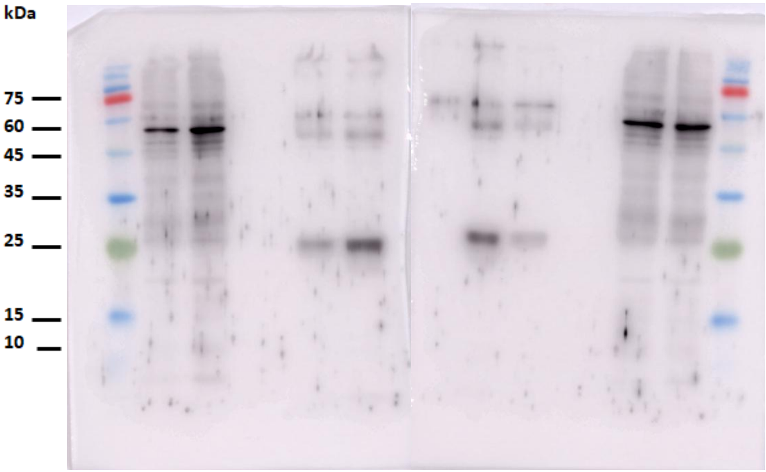

Blot 3

| M | PO | P<br>O<br>+<br>G<br>A | P<br>O | PO<br>+G<br>A |
|---|---|---|---|---|

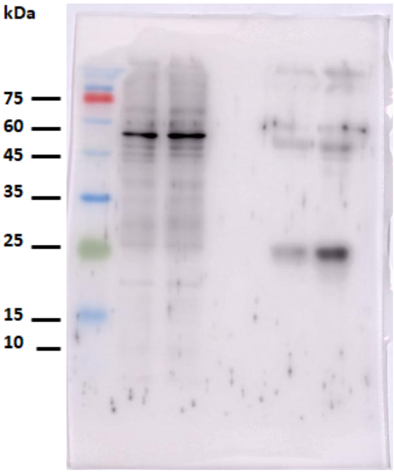

B. **Dot Blots for Fig.6L**

Blot 1

Representative

Original Inverted

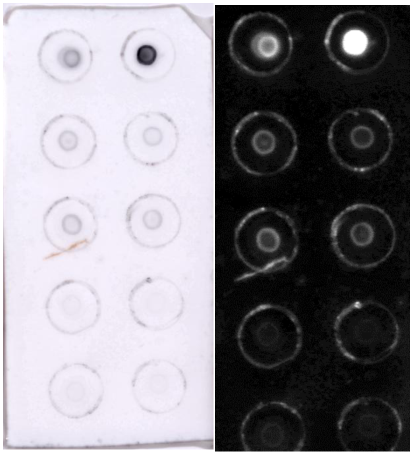

Blot 2

Original Inverted

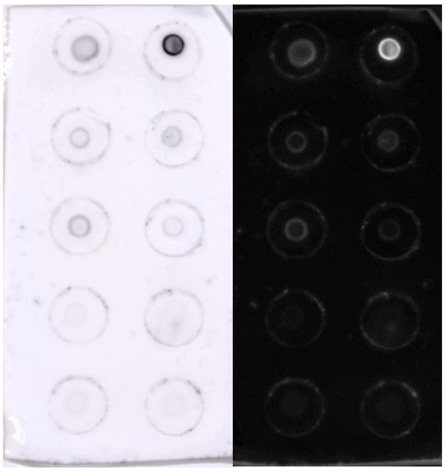

Blot 3

Original Inverted

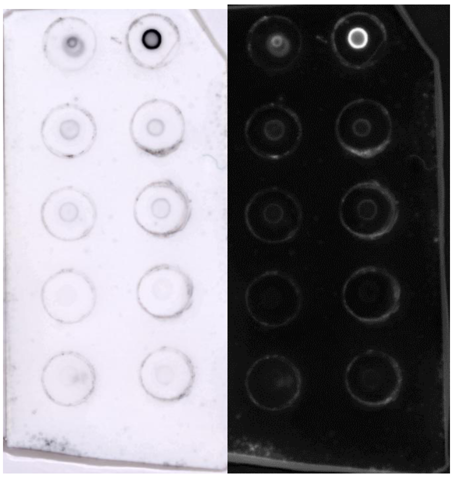

**(A) Uncut Western Blot images shown in Figure 6 J.(B)Dot blot Original Images for Figure 6L along with the represented (inverted images). The original images are captured in LAS500 and shown in the inverted mode for better visualization**

### <u>For Figure 8</u>

#### A. Western blot Images for Fig. 8A (ANP)

All independent biological samples are shown in the same blot

| | Young Donor Serum | | | | | Old Donor Serum | | | | Patients Serum | | | | |
|---|---|---|---|---|---|---|---|---|---|---|---|---|---|---|
| M | Y 1 | Y 2 | Y 3 | Y 4 | Y 5 | O 1 | O 2 | O 3 | O 4 | P 1 | P 2 | P 3 | P 4 | P 5 |

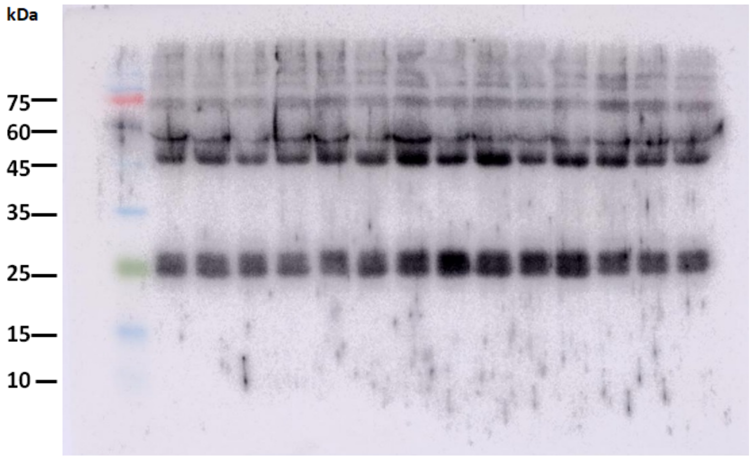

#### B. Western blot Images for Fig. 8A (Gal-3)

All independent biological samples are shown in the same blot

| | Young Donor Serum | | | | | Old Donor Serum | | | | Patients Serum | | | | |
|---|---|---|---|---|---|---|---|---|---|---|---|---|---|---|
| M | Y 1 | Y 2 | Y 3 | Y 4 | Y 5 | O 1 | O 2 | O 3 | O 4 | P 1 | P 2 | P 3 | P 4 | P 5 |

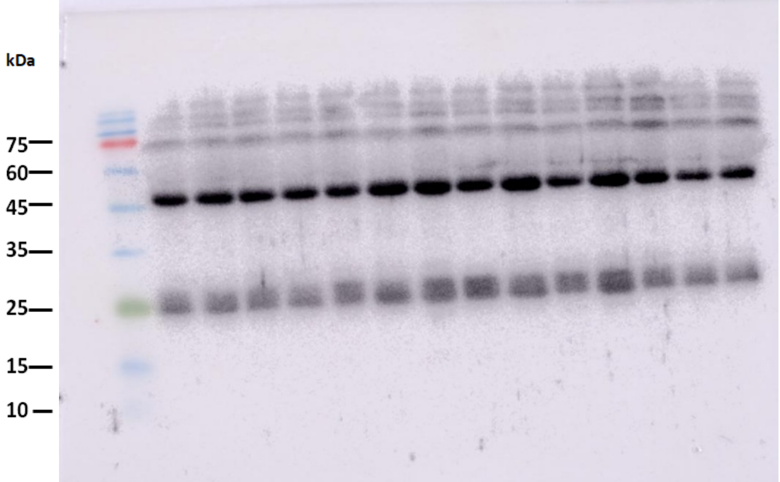

### For Figure 9

#### A. Dot Blots for Fig.9O

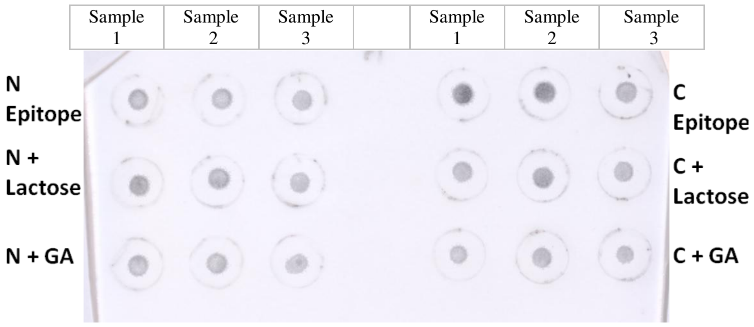

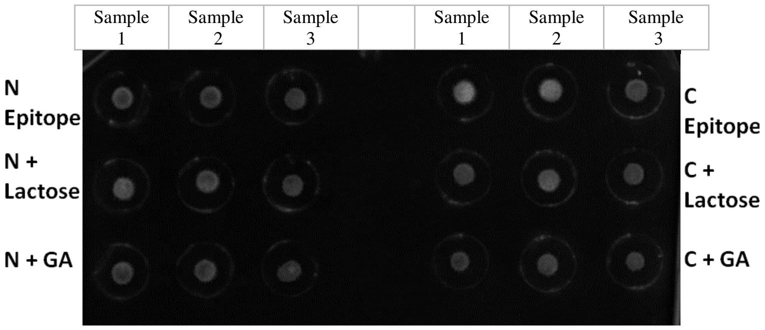

**Dot blot Original Images for Figure 9O along with the represented (inverted images) .Three independent samples are probed in each group. The original images are captured in LAS500 and shown in the inverted mode for better visualization.**

### For Fig. S2

#### A. Dot Blot ANP

Uncut Dot Blot Image

| | BA | BA+ AR | PO CH | PO-CH AR |
|---|---|---|---|---|

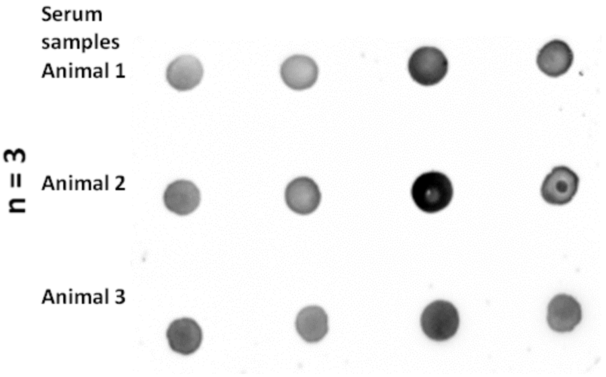

Representative Dot Blot Image (Inverted)

| | BA | BA+ AR | PO CH | PO-CH AR |
|---|---|---|---|---|

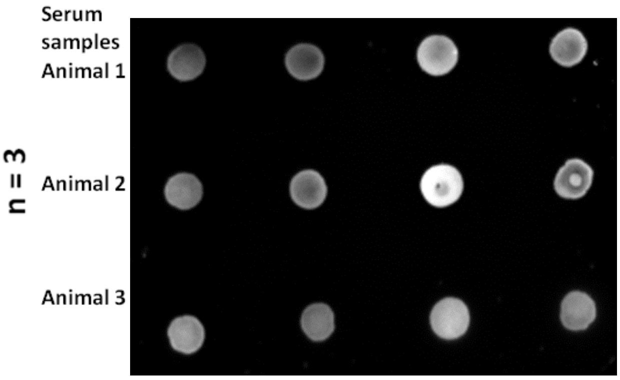

**B. Dot Blot GATA4**

Uncut Dot Blot Image

| | BA | BA+ AR | PO CH | PO-CH AR |
|---|---|---|---|---|

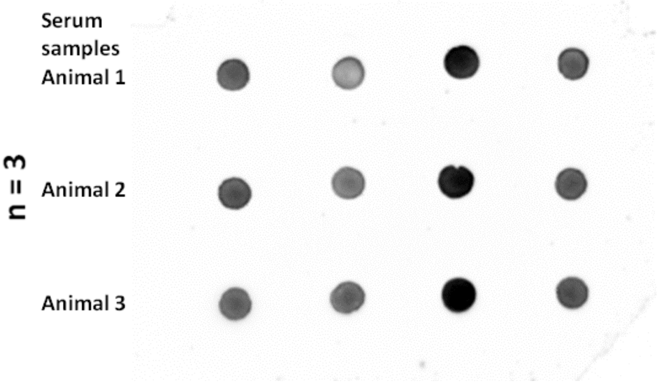

Representative Dot Blot Image (Inverted)

| | BA | BA+ AR | PO CH | PO-CH AR |
|---|---|---|---|---|

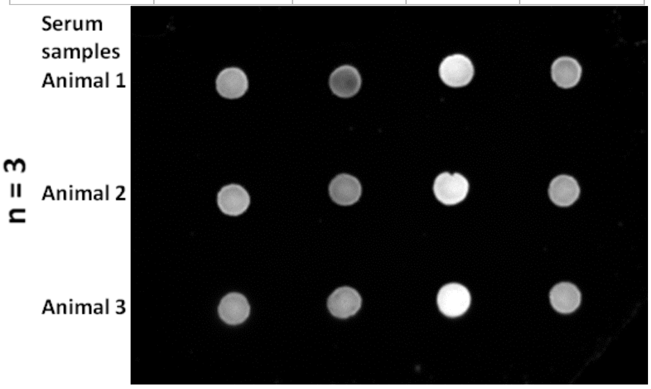

#### C. Dot Blot MYH7

Uncut Dot Blot Image

| | BA | BA+ AR | PO CH | PO-CH AR |
|---|---|---|---|---|

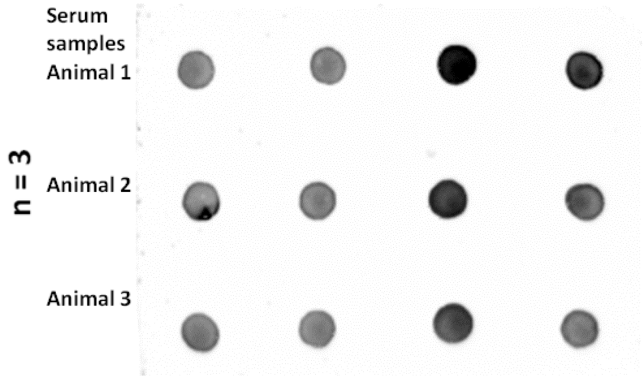

Representative Dot Blot Image (Inverted)

| | BA | BA+ AR | PO CH | PO-CH AR |
|---|---|---|---|---|

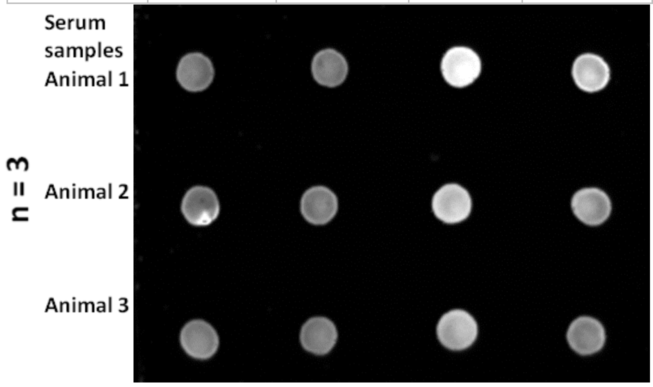

**D. Dot Blot MEF2**

Uncut Dot Blot Image

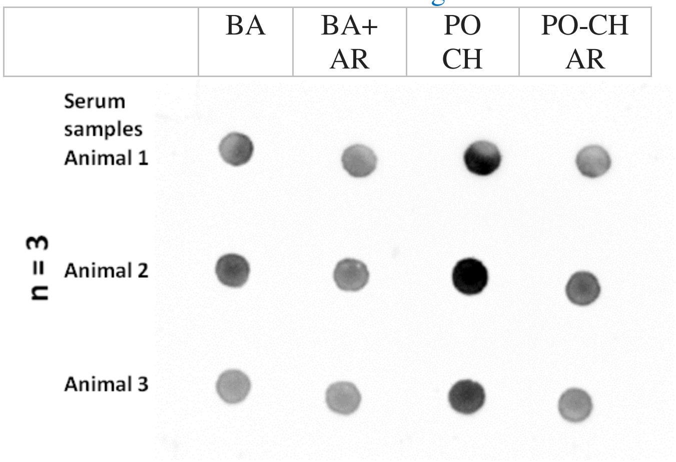

Representative Dot Blot Image (Inverted)

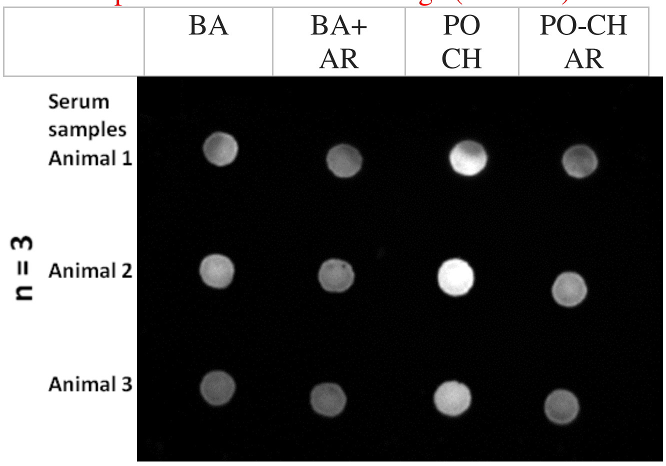

\

E. **Dot Blot cTnT**

Uncut Dot Blot Image

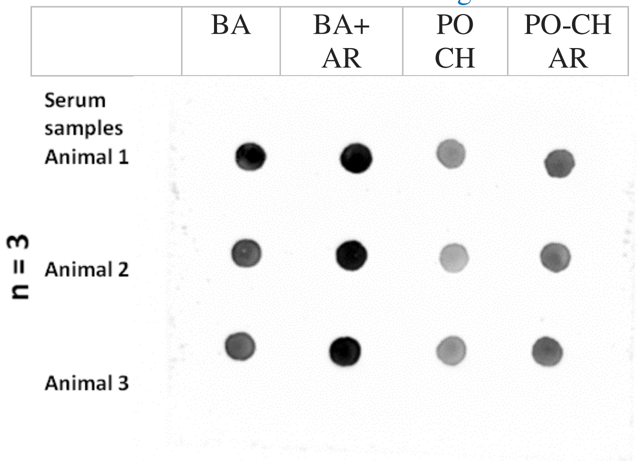

Representative Dot Blot Image (Inverted)

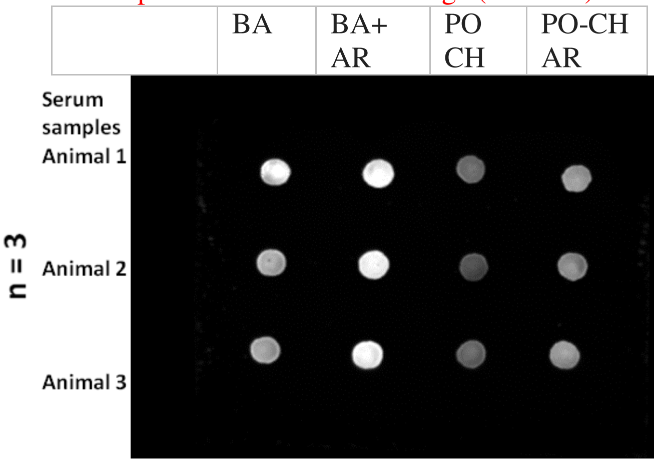

#### F. Dot Blot ACTN2

Uncut Dot Blot Image

| | BA | BA+ AR | PO CH | PO-CH AR |
|---|---|---|---|---|

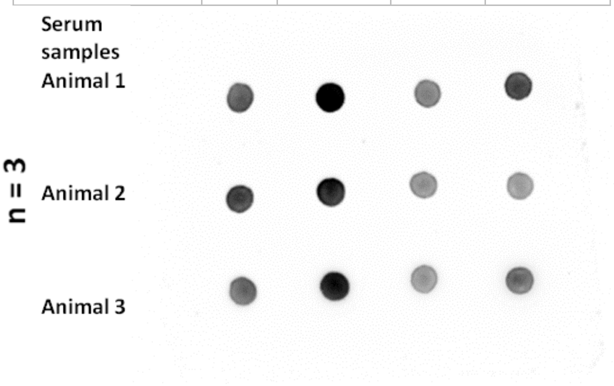

Representative Dot Blot Image (Inverted)

| | BA | BA+ AR | PO CH | PO-CH AR |
|---|---|---|---|---|

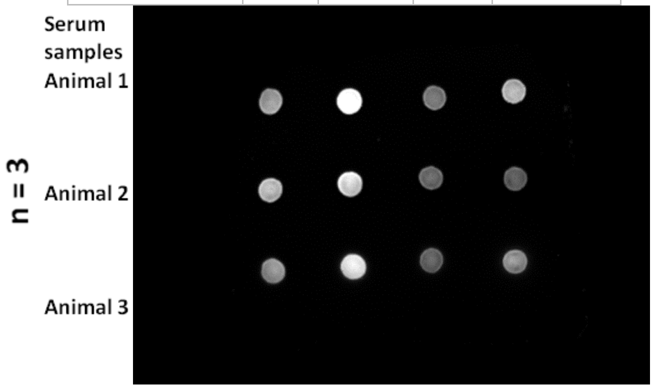

#### G. Dot Blot TMOD1

Uncut Dot Blot Image

| | BA | BA+ AR | PO CH | PO-CH AR |
|---|---|---|---|---|

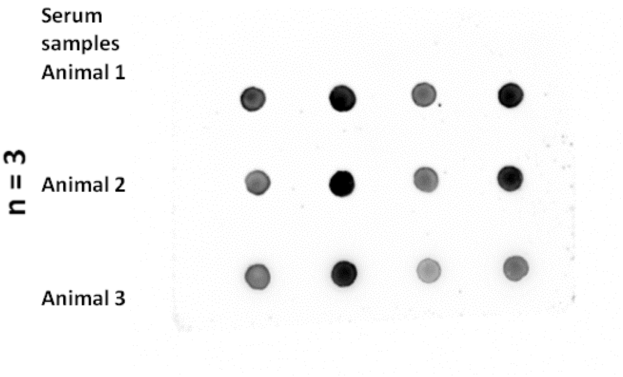

Representative Dot Blot Image (Inverted)

| | BA | BA+ AR | PO CH | PO-CH AR |
|---|---|---|---|---|

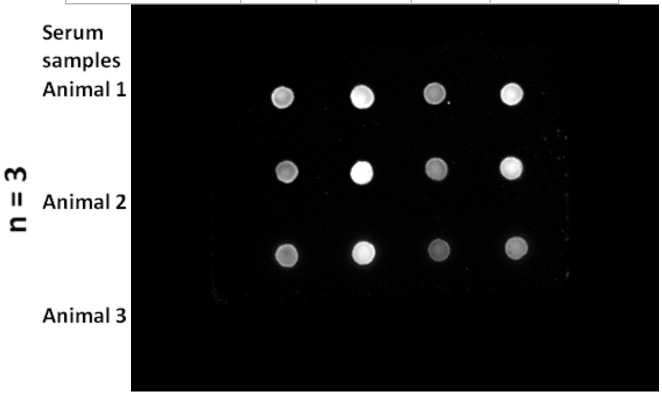

#### H. Dot Blot OBSCN

Uncut Dot Blot Image

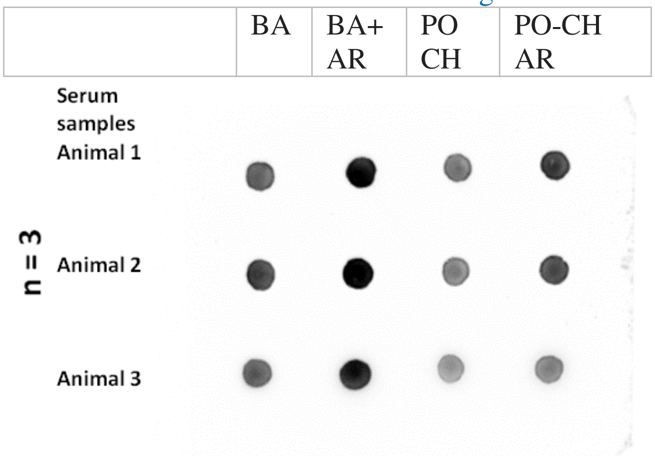

Representative Dot Blot Image (Inverted)

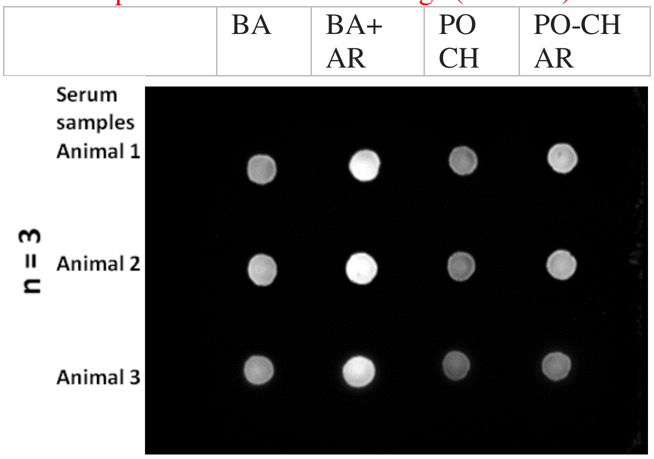

#### I. Dot Blot SERCA2A

Uncut Dot Blot Image

| | BA | BA+ AR | PO CH | PO-CH AR |
|---|---|---|---|---|

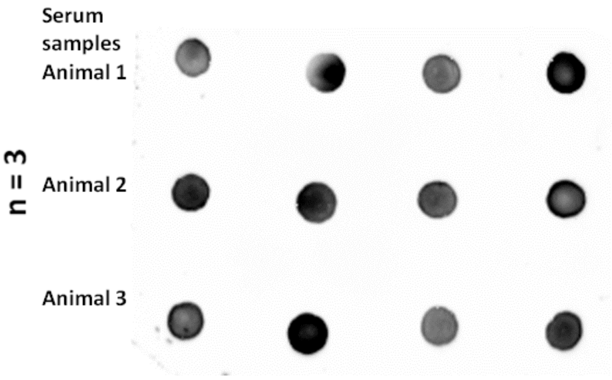

Representative Dot Blot Image (Inverted)

| | BA | BA+ AR | PO CH | PO-CH AR |
|---|---|---|---|---|

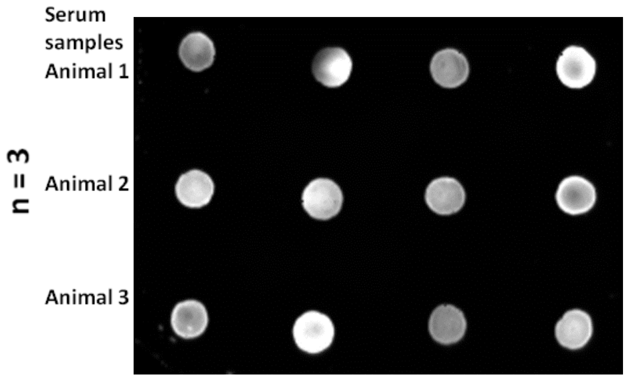

J. **Dot Blot CAMK2D**

Uncut Dot Blot Image

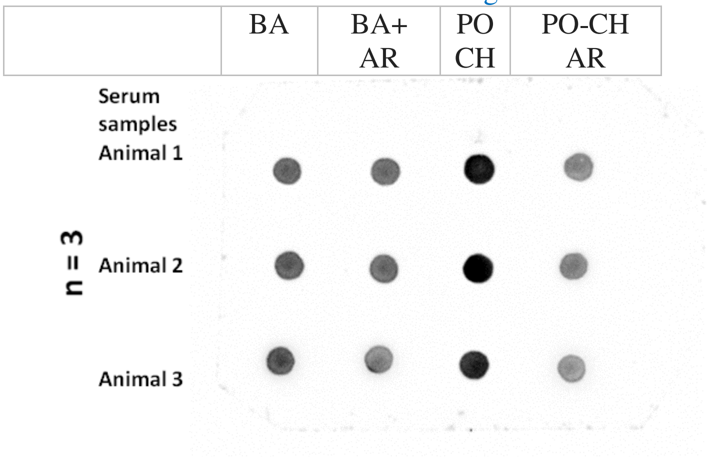

Representative Dot Blot Image (Inverted)

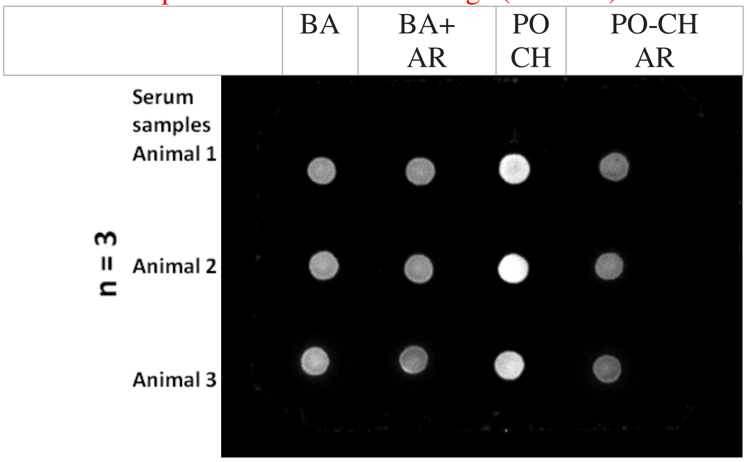

**K. Dot Blot CaNA, Fig.2K**

Uncut Dot Blot Image

| | BA | BA+ AR | PO CH | PO-CH AR |
|---|---|---|---|---|

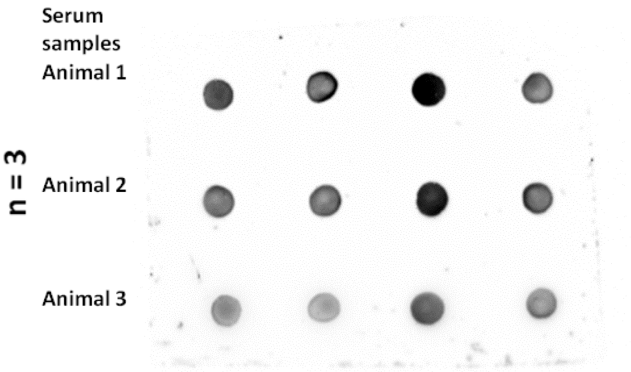

Representative Dot Blot Image (Inverted)

| | BA | BA+ AR | PO CH | PO-CH AR |
|---|---|---|---|---|

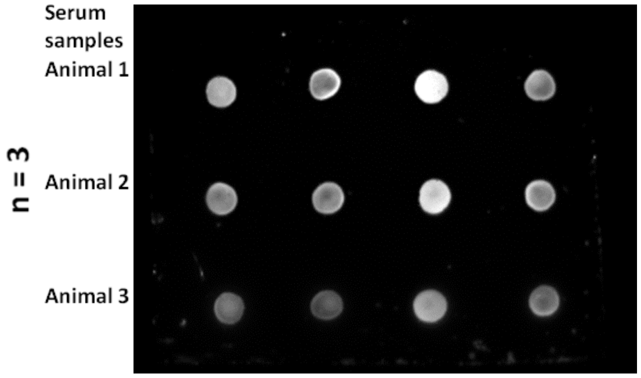

**L. Dot Blot NFATC2**

Uncut Dot Blot Image

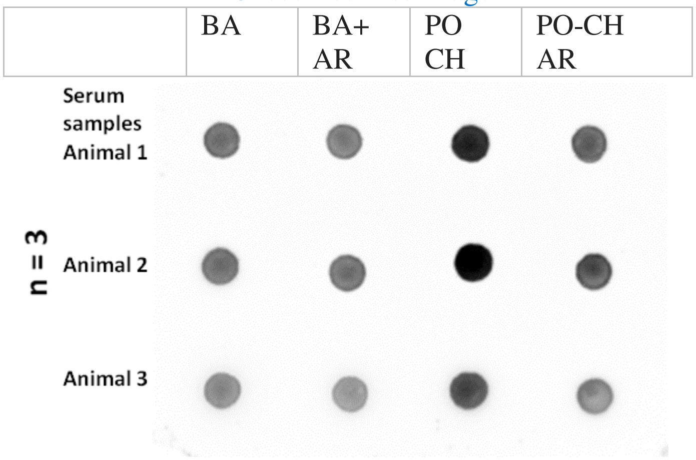

Representative Dot Blot Image (Inverted)

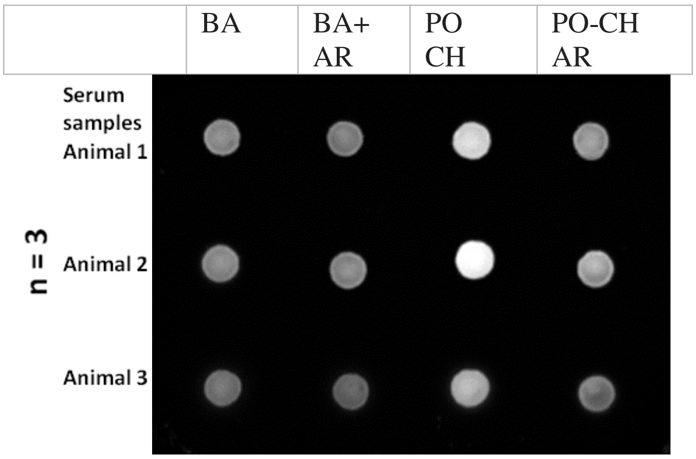

**A-l : Dot blot Original Images for Fig.S2along with the represented (inverted images). Three independent biological replicates in each group. The original images are captured in LAS500 and shown in the inverted mode for better visualization.**
